# Label-free identification of bacterial species using adapted vision transformers

**DOI:** 10.64898/2026.10.09.757828

**Authors:** Erik Hallström, Vinodh Kandavalli, M. Henar Marino Miguélez, Johan Elf, Carolina Wählby

**Author notes:** Corresponding author: Erik Hallström.

## Abstract

Antibiotic resistance is an increasingly serious problem worldwide. Identifying the bacterial species can help guide antibiotic selection and improve patient outcomes. In sepsis, where every minute counts, rapid identification is particularly important, and new approaches are needed.

In this study, deep learning models were trained to identify bacterial species from phase-contrast time-lapse images of cells growing in microfluidic traps, without fluorescent labels. We demonstrated this approach in two previous studies using laboratory strains. Here, the previous work is extended to clinical bacterial isolates, evaluating performance on isolates not seen during training. Furthermore, it was tested whether autofluorescence could aid species identification.

Two validation sets were acquired in separate experiments from the training data, the first containing clinical-isolate traps with five species (*Escherichia coli*, *Enterococcus faecalis*, *Klebsiella pneumoniae*, *Staphylococcus aureus* and *Enterobacter cloacae*), imaged every two minutes, and the second containing clinical isolates of two species (*E. coli* and *K. pneumoniae*) recovered from spiked donor blood, imaged every minute.

Training with the best previous CNN method achieved unsatisfactory results. Therefore, vision transformers were explored, and a self-supervised adaptation procedure was developed for pretrained DINOv3 using a frozen teacher, followed by supervised classification using time-lapse montages. Mean macro-F1 was 93.6 *±* 1.3% for isolates precultured in Mueller–Hinton medium using 60 frames (118 minutes) and 87.2 *±* 2.7% for isolates precultured in blood using 55 frames (54 minutes) (mean *±* sample standard deviation across training seeds).

Performance was evaluated over time by increasing the number of frames available to the model during testing, and Fourier-domain filtering approximated reduced numerical aperture. The results are promising and suggest that the aperture can be decreased while retaining good results. However, further evaluation on independent isolates, experiments and patient samples is needed to establish generalization. Together with previous findings, these results bring us closer to rapid image-based diagnostics that could transform the initial treatment of acute bacterial infections by enabling better-informed antibiotic selection.

## 1 Introduction

### 1.1 Bacterial infections and current diagnostic workflow

Bacterial infections remain a major challenge, with millions of deaths worldwide each year [1]. Antibiotic resistance makes these infections increasingly difficult to treat and was estimated to directly cause 1.27 million deaths in 2019 [2].

One particularly serious condition is sepsis, in which the body’s immune response to infection causes organs to fail [3]. Broad-spectrum antibiotics are often given before the bacteria causing the infection have been identified. The diagnostic workflow typically starts with taking a blood sample for overnight culturing [4]. Species identification is then commonly performed using MALDI-TOF mass spectrometry, a highly accurate method for identifying the bacterial species that typically requires visible colonies or a positive blood culture [5]. After blood culturing, antibiotic susceptibility can be investigated using phenotypic methods, which observe how antibiotics affect bacterial growth, or genotypic methods, which look for known resistance genes [6]. Obtaining species information is important because it can narrow the choice of antibiotics and help select the appropriate breakpoints for interpreting susceptibility results [5, 7]. Obtaining this information earlier can reduce the use of broad-spectrum antibiotics, improve treatment decisions and potentially improve patient outcomes [6]. However, the time required by conventional culture-based workflows is a major limitation. New approaches to rapid bacterial identification are therefore needed to support earlier and better-informed treatment decisions.

### 1.2 Species identification from phase-contrast time-lapse images

In this study, we use a microfluidic device consisting of narrow bacterial traps [8]. The cells become trapped and then grow and multiply as growth medium flows past them, with continued growth pushing cells outwards. The cell at the closed end of the trap gives rise to successive generations of cells, hence the name “mother machine.” Phase-contrast microscopy is used to record an image, typically once every other minute. We have previously demonstrated species identification from phase-contrast time-lapse videos using deep artificial neural networks [9] in two studies involving four and seven bacterial species [10, 11]. However, both studies used laboratory strains, with limited variation among isolates of the same species. Here, the investigation is extended to clinical bacterial isolates. In our previous studies, Video ResNet [12], a convolutional neural network (CNN) adapted for videos, performed best. However, even after training on clinical-isolate data, the model did not generalize well to new clinical isolates. Therefore, a self-supervised adaptation procedure was developed for a pretrained DINOv3 vision transformer [13], processing the video clips as frame montages, followed by supervised fine-tuning for species classification.

### 1.3 Dataset

Images from previous studies [10, 11] and new experiments were used together to form the dataset, including 53 microfluidic experiments with a total of 73,067 labelled, non-empty traps (see “Microfluidic experiments and dataset” in Materials and Methods). Each experiment was performed in one microfluidic chip. Hereafter, individual experiments are referred to as “chips”. Two evaluation sets were used. The first, hereafter named Clear medium, included chips 41–43, containing 5,790 clinical-isolate traps with five species (*E. coli*, *E. faecalis*, *K. pneumoniae*, *S. aureus* and *E. cloacae*), imaged every two minutes. The second, hereafter named Blood, included chips 44–53, containing 560 traps with clinical isolates of *E. coli* and *K. pneumoniae* recovered from spiked donor blood, imaged every minute. Both evaluation sets contained isolates that were not used in training. The blood samples were processed using a protocol developed by Marino Miguélez et al. [4]. The protocol reduced debris, producing clearer images for downstream analysis.

Autofluorescence images were also acquired in the new experiments on chips 25–43. These images were obtained after the phase-contrast sequence by illuminating the cells with 488-nm excitation light and recording their naturally occurring fluorescence, without adding fluorescent labels [14]. We investigated whether this provides additional information for species identification. Examples of the phase-contrast time-lapses and corresponding autofluorescence images are shown in Fig 1.

**Fig 1.**
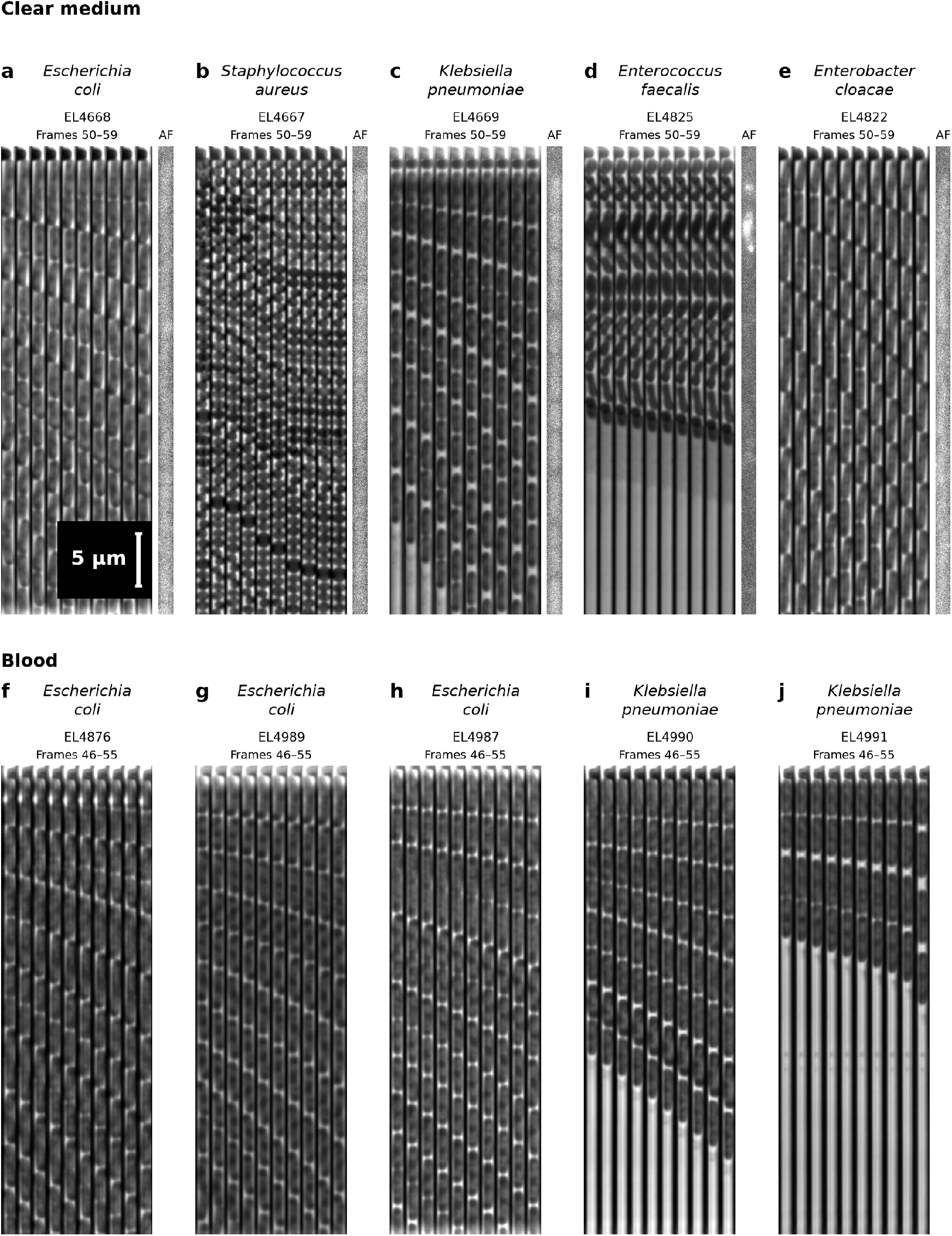
Phase-contrast examples of all evaluation isolates. Each panel shows ten consecutive frames from one selected microfluidic trap with one bacterial isolate. Clear medium (a–e) shows frames 50–59, acquired every two minutes, followed by matching autofluorescence (AF, 488 nm): (a) *E. coli* EL4668, (b) *S. aureus* EL4667, (c) *K. pneumoniae* EL4669, (d) *E. faecalis* EL4825, and (e) *E. cloacae* EL4822. Blood (f–j) shows frames 46–55, acquired every minute, with phase contrast only: (f) *E. coli* EL4876, (g) *E. coli* EL4989, (h) *E. coli* EL4987, (i) *K. pneumoniae* EL4990, and (j) *K. pneumoniae* EL4991. Images are normalized and clipped for visibility. The 5 *µ*m scale bar shown in (a) applies to all panels.

### 1.4 Related work

Apart from our previous studies, we are unaware of other studies that identify bacterial species from single-cell phase-contrast time-lapse videos using deep learning. However, several related approaches exist. A closely related study is that of Kandavalli et al., which provided the data for our seven-species study [8, 11]. It used the same microfluidic platform but identified species using fluorescence in situ hybridization (FISH), requiring species-specific fluorescent probes and fixation that kills the bacteria, preventing further analysis of their growth [8]. Similarly, Chin et al. combined time-lapse imaging for phenotypic AST with sequential FISH for species identification after enriching bacteria from whole blood [15]. In contrast, our study achieves species identification using only spatiotemporal features from phase-contrast time-lapse images (video clips). Other approaches detect and identify growing microcolonies [16, 17], but require several hours for colonies to develop. Further approaches use Raman spectroscopy for species identification or light-scattering measurements of bacteria in droplets to assess antibiotic susceptibility [18, 19]. Many promising approaches are therefore being developed in parallel. An advantage of our approach is its simplicity: species identification uses the phase-contrast images already acquired to monitor bacterial growth, without requiring additional staining or a separate measurement system.

## 2 Results

### 2.1 Testing best models from previous studies on new data

We started the study by testing the previous best-performing models on new data produced in this study. A number of new experiments containing laboratory and clinical isolates were performed, as described in “Microfluidic experiments and dataset” in Materials and Methods. The complete dataset consisted of 53 microfluidic chips, including 29 newly collected chips, each assigned a chip number. The models in both our previous studies [10, 11] had been evaluated on test sets, which effectively served as validation sets when selecting the best models. The selected models had not subsequently been tested on new data. The new experiments allowed us to do this. The best-performing architecture in both previous studies was R(2+1)D Video ResNet [12], achieving macro-F1 scores of 99.8% and 93.6%, respectively.

First, the four-species model was tested on the test set of the seven-species study (chip 16), restricted to its four supported species, and on data from new experiments with laboratory strains (chips 25–27). Then, the seven-species model was tested on the new laboratory data, containing six of its seven species, excluding *P. mirabilis*. The four-species model achieved macro-F1 scores of 53.7% and 89.0% on the seven-species study’s data and the new laboratory data, respectively, suggesting that transfer to new experiments is not trivial. The seven-species model achieved a macro-F1 score of 92.3%, close to the previous validation score of 93.6%.

Then, we compared the results on laboratory strains in new experiments (chips 25–27) and new clinical isolates (chips 28–40). Because no clinical isolates of *P. aeruginosa* were available, we matched three species for the four-species model and five species for the seven-species model (excluding *P. aeruginosa* and *P. mirabilis*) between the laboratory and clinical data. The four-species model achieved a macro-F1 score of 89.0% on laboratory isolates and 74.8% on clinical isolates, a drop of 14.1 percentage points. The seven-species model achieved 92.2% on laboratory isolates and 62.3% on clinical isolates, a drop of 29.9 percentage points. This suggests that performance on laboratory isolates did not fully transfer to clinical isolates, although differences between experiments may also have contributed to the decrease.

Finally, the Video ResNet model was trained using the seven-species study’s training recipe on clinical isolates from chips 28–40 and tested on clinical isolates from chips 41–43. The model achieved a macro-F1 score of 66.6% across the five species present in the test set. Thus, even after training on clinical isolates, performance remained limited on new clinical experiments, motivating us to explore alternative modelling approaches.

The full evaluation, including confusion matrices, is presented in S1 Document.

### 2.2 Testing on autofluorescence

Autofluorescence was also explored for classification of the five species in the Clear medium benchmark, using ResNet18 [20] trained on clinical isolates from chips 28–40 and evaluated on chips 41–43 with 31 paired training seeds per input condition. Autofluorescence alone achieved a mean macro-F1 score of 52.6%, compared with 62.6% using the last phase-contrast frame alone and 65.9% when combining both inputs. Adding autofluorescence thus improved mean macro-F1 slightly, suggesting that it provided additional information for species classification under the tested conditions. These results are exploratory, as the same evaluation chips had been used in previous comparisons when developing the training recipe. Although autofluorescence may be interesting for further investigation, we did not pursue this approach further in the present study.

The full results are available in S2 Document.

### 2.3 Adapting vision transformers for bacterial species identification

All models in this and the following sections were trained 31 times with different random seeds. Results are reported as the mean *±* standard deviation across these runs.

One interesting observation in both our previous studies was that classification performance remained high even after substantial downsampling [10, 11], suggesting that the images were oversampled. After checking the microscope specifications, the images were confirmed to be oversampled (see Materials and Methods). The first step was therefore to downsample the cropped trap images from 42 pixels to 16 pixels in width. The resulting spatial sampling rate remained above the Nyquist limit (see Materials and Methods). Area interpolation was used to approximate sampling with a sensor containing fewer pixels. This preprocessing was separate from the Fourier-domain filtering used later to simulate optics with a lower numerical aperture during evaluation.

Due to the poor performance of CNNs on these data, vision transformers were explored. Ten consecutive time-lapse frames were concatenated horizontally into a montage, following the montage approach used in our previous study [4]. With DINOv3’s 16 *×* 16-pixel patches, each patch spans the width of one frame and a short segment along the trap. Thus, the model can relate image information both along the growing cell column and across time. We first tried an off-the-shelf DINOv3 Small model with approximately 21 million parameters, using either full fine-tuning with layer-wise learning-rate decay or low-rank adaptation (LoRA) [21] applied to selected transformer blocks. These approaches achieved mean macro-F1 scores of 83.0*±*1.2% and 81.5*±*3.6%, respectively, on Clear medium, and 68.3 *±* 1.9% and 61.3 *±* 5.7% on Blood. The lower performance on Blood motivated a further step to adapt the pretrained features to bacterial images.

In this study, a two-stage procedure was developed to adapt a pretrained DINOv3 Small model to the bacterial time-lapses (see “Model and training” in Materials and Methods). In the first stage, a student model received images with added noise and masked patches and learned to match features produced from the clean images by a frozen teacher. This stage used no species labels. In the second stage, the adapted model was trained with species labels to predict one of eight species. Both stages used LoRA [21], keeping the pretrained backbone weights fixed while training a small set of additional parameters. The adapted model achieved mean macro-F1 scores of 93.6 *±* 1.3% on Clear medium using 60 frames (118 minutes) and 87.2 *±* 2.7% on Blood using 55 frames (54 minutes) (Table 1). Confusion matrices and receiver operating characteristic (ROC) curves for the median-performing training run are shown for the Clear medium dataset in Fig 2 and the Blood dataset in Fig 3. Example recordings of each species are shown in S1–S7 Figs, and examples of misclassified traps, with a comment on each, in S8–S32 Figs.

**Fig 2.**
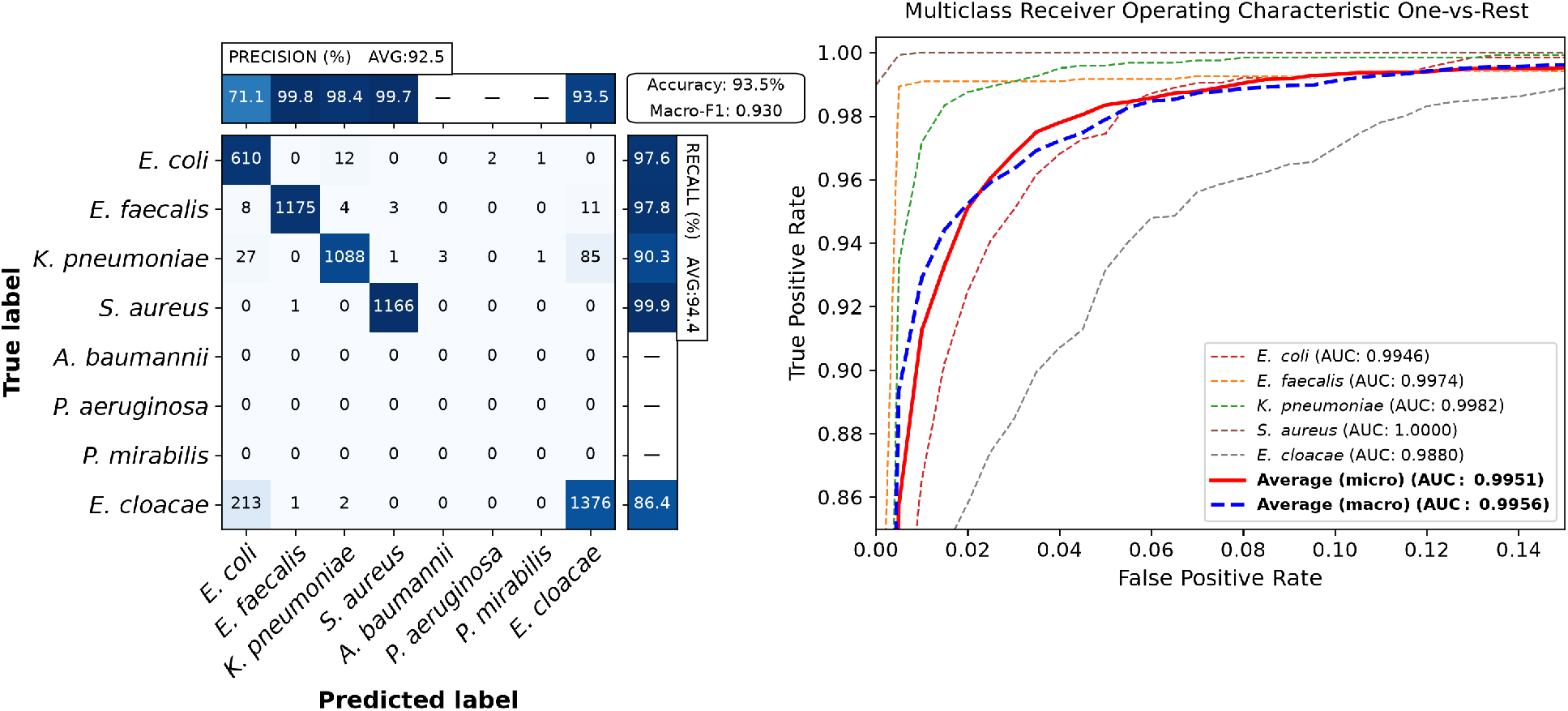
Classification results of the main model on the Clear medium dataset. Results are shown for the median-performing model using 60 frames (118 minutes). Left: confusion matrix showing trap counts, with colors indicating row percentages. The top and right bars show precision and recall, respectively, with both values and colors indicating percentages. Precision and recall are shown only for species present in the evaluation dataset. Right: receiver operating characteristic (ROC) curves. The ROC axes are zoomed for visibility.

**Fig 3.**
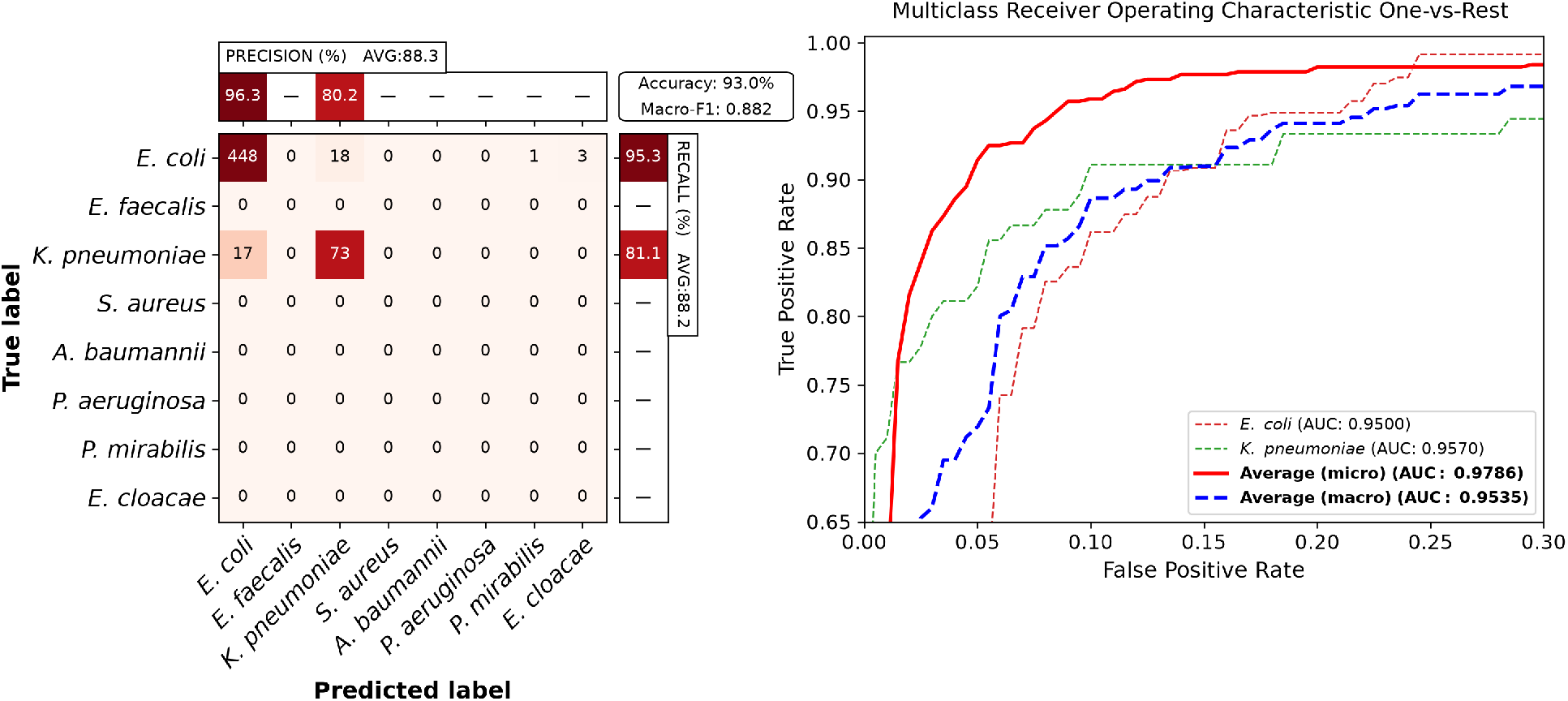
Classification results of the main model on the Blood dataset. Results are shown for the same representative model as in Fig 2, using 55 frames (54 minutes). Left: confusion matrix showing trap counts, with colors indicating row percentages. The top and right bars show precision and recall, respectively, with both values and colors indicating percentages. Precision and recall are shown only for species present in the evaluation dataset. Right: receiver operating characteristic (ROC) curves. The ROC axes are zoomed for visibility.

**Table 1:** Classification performance of the DINOv3 Small models. All procedures used the same training and evaluation sets, as described in Sections 1.3 and 3.3. Values are mean macro-F1 *±* sample standard deviation across training seeds. Predictions retain all eight output classes, while macro-F1 is averaged over the species present in each evaluation set.

| Training procedure | Clear medium (%) | Blood (%) |
| --- | --- | --- |
| Supervised full fine-tuning | $83.0 \pm 1.2$ | $68.3 \pm 1.9$ |
| Supervised LoRA | $81.5 \pm 3.6$ | $61.3 \pm 5.7$ |
| Self-supervised adaptation followed by supervised LoRA | $93.6 \pm 1.3$ | $87.2 \pm 2.7$ |

### 2.4 Species identification with shorter observation times

The performance of our model was tested by varying the number of input frames during testing, without retraining the model. In a clinical situation, it would be important to obtain the information as early as possible. Mean macro-F1 increased from 66.1 *±* 2.4% using only the first frame at time zero to 85.5 *±* 1.5% at 50 minutes on Clear medium, and from 54.9 *±* 6.0% to 86.7 *±* 2.9% on Blood (Fig 4). Note that the Blood test set was sampled at one frame per minute and Clear medium at one frame every two minutes. The full evaluation sequences of 60 frames for Clear medium and 55 frames for Blood correspond to 118 and 54 minutes after the first frame, respectively. Note also that these times exclude sample preparation and earlier imaging stages.

**Fig 4.**
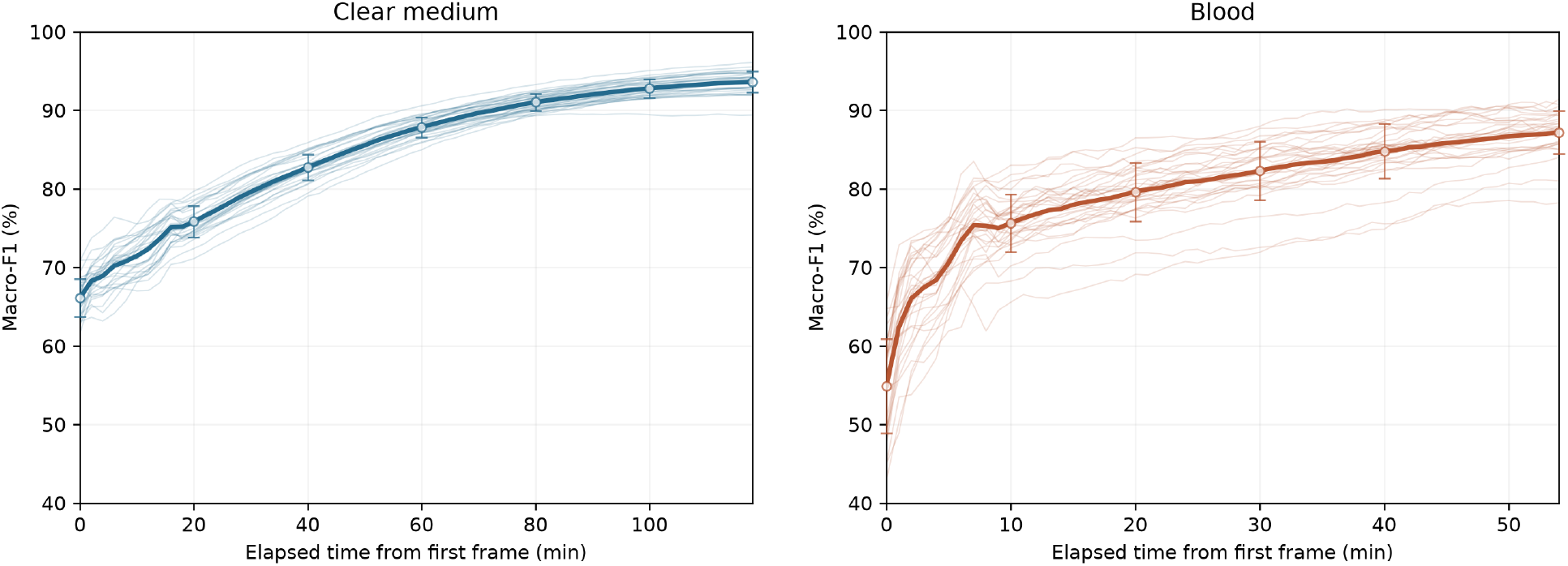
Classification from increasing observation time. Thin lines show individual seed curves across 31 training runs. Thick lines show the mean macro-F1. Hollow circles show the mean, and error bars show the standard deviation at selected time points.

### 2.5 Simulating a lower numerical aperture

Analogous to our previous studies, the model was tested by removing spatial detail from the time-lapse images, mimicking imaging with a microscope in a low-resource setting. Previously, Lanczos downsampling was used. However, this does not closely simulate the images that would be acquired with such a microscope. Instead, a lower numerical aperture (NA) was approximated by filtering the images in the Fourier domain (see Materials and Methods). The model was evaluated across a series of spatial filters and observation times. Interestingly, performance on Clear medium remained relatively stable down to a nominal NA of 0.7, while performance on Blood decreased more gradually. Below this value, performance dropped sharply on Clear medium and continued to decrease on Blood (Fig 5). Examples of the filtered images are shown in S33–S41 Figs and are visualized in S1–S10 Videos.

**Fig 5.**
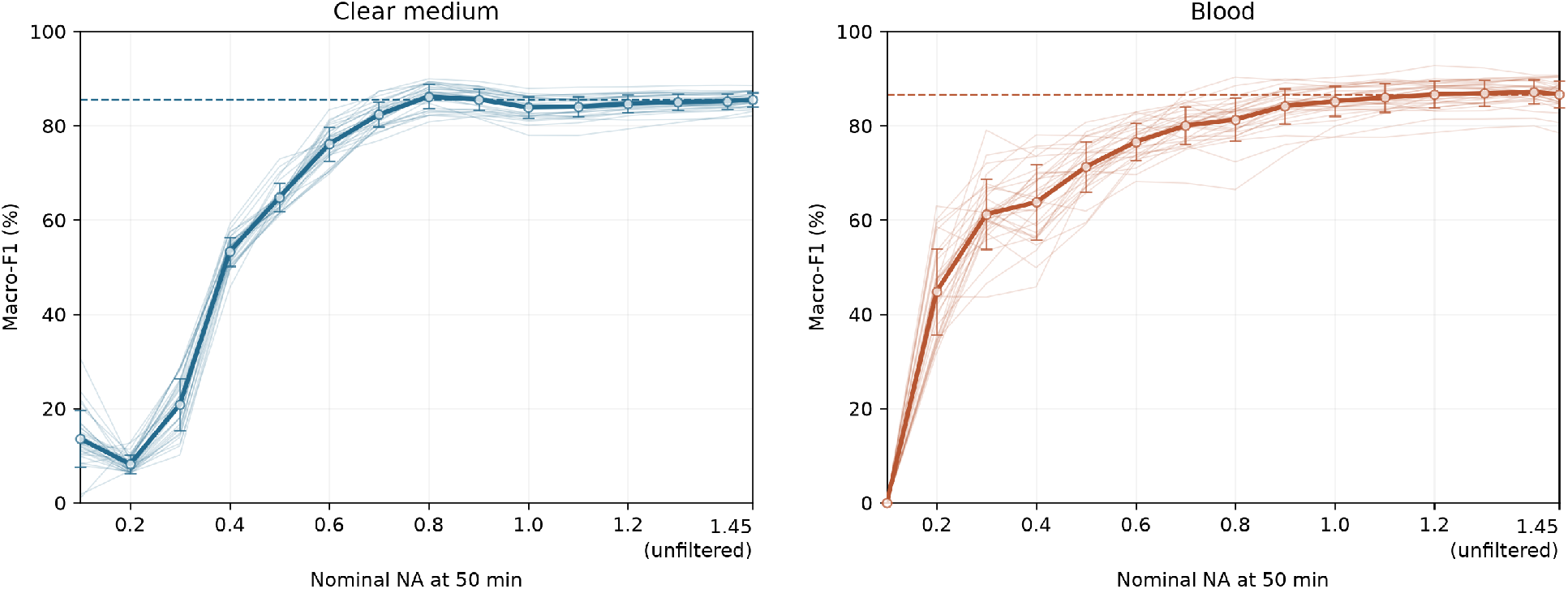
Sensitivity to spatial filtering at 50 minutes. Clear medium is shown on the left and Blood on the right. Thin lines show individual seed curves and thick lines show mean macro-F1 across 31 training seeds. Hollow circles show the mean, and error bars show the standard deviation at each evaluated NA. The point at NA 1.45 shows the unfiltered reference, with no additional Fourier-domain NA filtering and with standard preprocessing. Dashed lines mark the same unfiltered mean. Fifty minutes corresponds to frame 26 for Clear medium and frame 51 for Blood.

### 2.6 Ablation study

A number of ablations were performed during training to investigate what mattered in the model. First, the models were trained with different pretrained backbones, varying between DINO, DINOv2 and DINOv3 [22, 23, 13]. DINOv3 clearly outperformed DINO and DINOv2 on both datasets, suggesting that foundation-model features continue to improve and can transfer to image domains beyond those used for their pretraining.

Then, we changed parts of the self-supervised adaptation (SSL) stage, removing patch masking, noise and regularization. Finally, the augmentations, classifier initialization and warm-up used during supervised fine-tuning were varied (Table 2). Compared with zero initialization, random initialization of the linear classification layer reduced performance on both datasets and increased variation across training runs. Adding 50 warm-up steps recovered some of the performance. Possibly, this is because both approaches allow the classifier to adapt to the already useful pretrained features before the model starts modifying them.

**Table 2:**
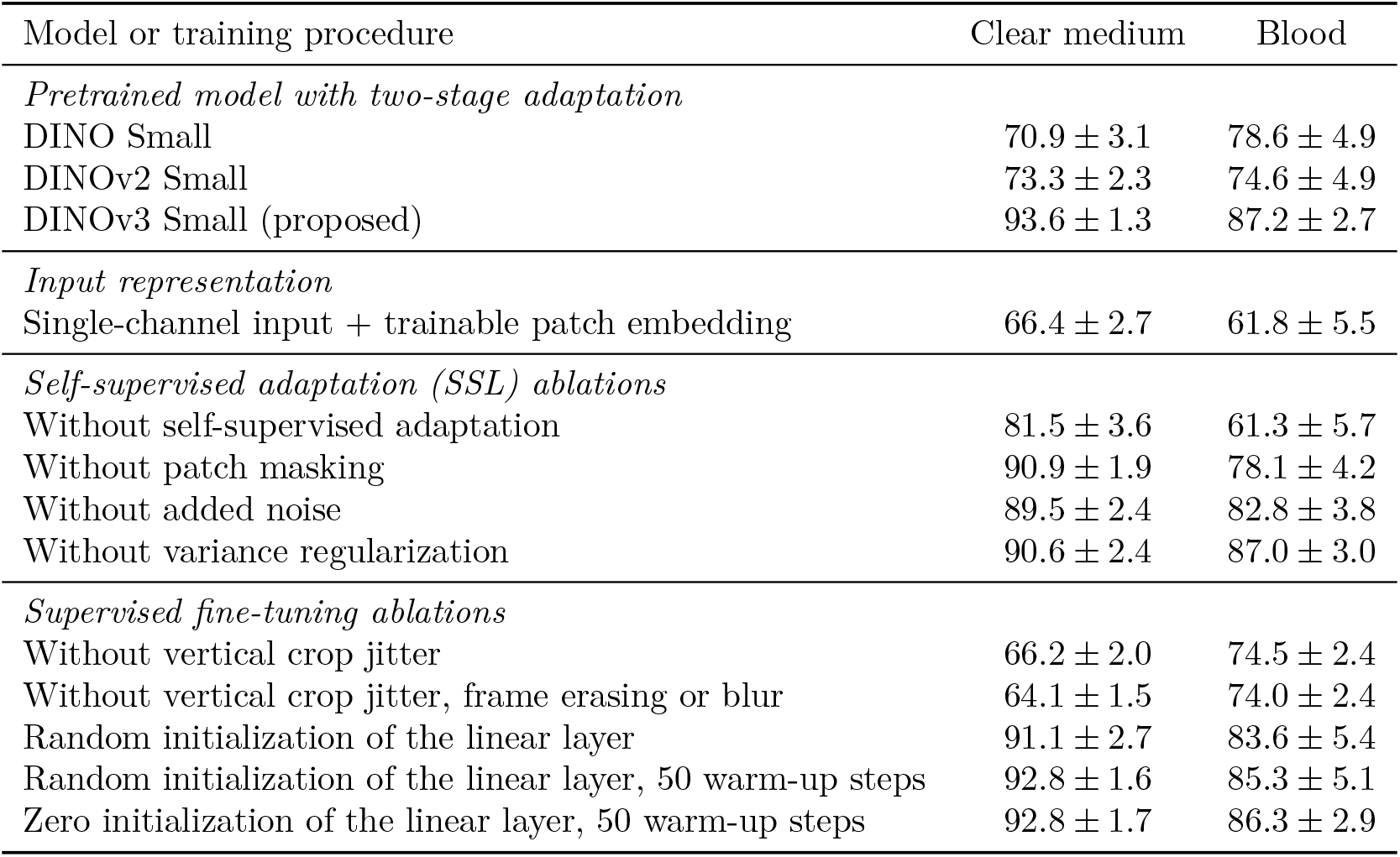
Pretrained-model and training-component ablations. Mean macro-F1 *±* sample standard deviation (%). Clear medium uses 60 frames and Blood uses 55 frames.

Frame erasing augmentation was included when training the model, masking out the last one to nine consecutive frames on the right-hand side of the montage (retaining at least one frame). This was done to improve performance when evaluating fewer than ten frames. During evaluation, sequences longer than ten frames were processed using sliding windows of ten frames with a stride of one, and the resulting logits were averaged. With fewer than ten frames, the remaining frame positions were masked out because the transformer used a fixed-size input. Furthermore, blur augmentation was added to make the model perform better at lower simulated NA. Models without these augmentations were trained and tested at varying evaluation times and simulated NA values to test their benefit. Results are shown in Fig 6. Frame erasing improved performance for the shortest recordings in both datasets, with the difference becoming smaller as the number of available frames approached ten. The blur augmentation was also clearly beneficial.

**Fig 6.**
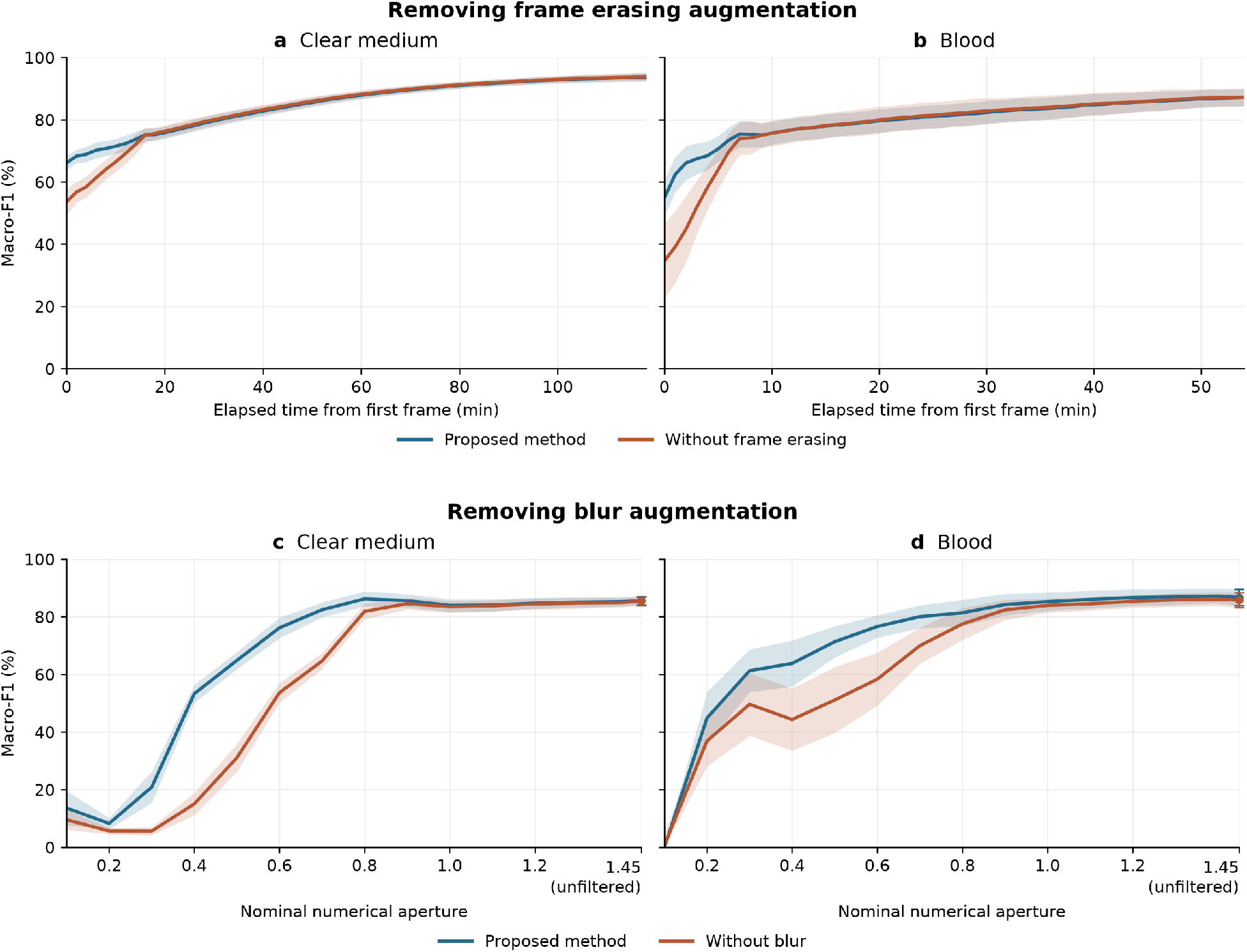
Effects of removing frame erasing and blur augmentation during training. (a, b) Macro-F1 over time with and without frame erasing for Clear medium (a) and Blood (b). (c, d) Macro-F1 at different simulated NA values with and without blur during training for Clear medium (c) and Blood (d), evaluated at 50 minutes. The points at NA 1.45 show the unfiltered reference, with no additional Fourier-domain NA filtering and with standard preprocessing. Lines show the mean macro-F1 across 31 training runs, and shading and reference error bars indicate *±* one sample standard deviation.

Then, the contribution of individual training datasets was tested by excluding each dataset from self-supervised adaptation alone or from both training stages (Table 3). Excluding each dataset from SSL alone had relatively little effect on Clear-medium performance. On Blood, excluding the mixed-isolate experiments from SSL reduced mean macro-F1 from 87.2% to 83.7%. However, removing the entire SSL stage reduced performance, particularly on Blood, as shown earlier (Table 2). Excluding datasets from both stages generally had a larger effect, especially when removing clinical isolates from the single-isolate or mixed-isolate experiments (chips 28–40). This suggests that including variation among clinical isolates during training was important for identifying new isolates.

**Table 3:** Effect of excluding training datasets. Macro-F1 (mean *±* sample SD) at 60 Clear-medium frames and 55 Blood frames.

| Dataset excluded | Clear medium (%) | Blood (%) |
| --- | --- | --- |
| All training data (reference) | 93.6 ± 1.3 | 87.2 ± 2.7 |
| *Excluded from self-supervised adaptation only* |  |  |
| Four-species study | 93.8 ± 1.4 | 86.4 ± 3.1 |
| Seven-species study | 93.3 ± 1.1 | 86.0 ± 2.8 |
| New laboratory strains | 93.4 ± 1.5 | 86.9 ± 2.8 |
| Single-isolate experiments | 93.7 ± 1.4 | 86.8 ± 2.6 |
| Mixed-isolate experiments | 93.0 ± 1.4 | 83.7 ± 4.4 |
| *Excluded from both SSL and supervised fine-tuning* |  |  |
| Four-species study | 94.4 ± 1.1 | 75.5 ± 4.0 |
| Seven-species study | 93.0 ± 1.4 | 82.4 ± 3.5 |
| New laboratory strains | 93.1 ± 1.5 | 84.0 ± 3.0 |
| Single-isolate experiments | 69.0 ± 0.8 | 74.0 ± 4.4 |
| Mixed-isolate experiments | 78.2 ± 2.4 | 79.4 ± 4.0 |

## 3 Materials & Methods

For full details, refer to the replication package [24, 25], which contains all code and data needed to reproduce all deep learning experiments in this paper.

### 3.1 Microfluidic experiments and dataset

The microfluidic chip used in these studies was introduced by Baltekin et al. [26] and contains narrow bacterial traps with a small constriction at one end, allowing growth medium to flow through while stopping bacteria from escaping. Images from our previously published four-species [10] and seven-species [11] studies were used together with new experiments, including 53 microfluidic chips and 73,067 labelled, non-empty traps (Table 4). For the validation sets, only traps containing bacteria from the start and throughout the time-lapse were included, removing traps with empty frames.

**Table 4:**
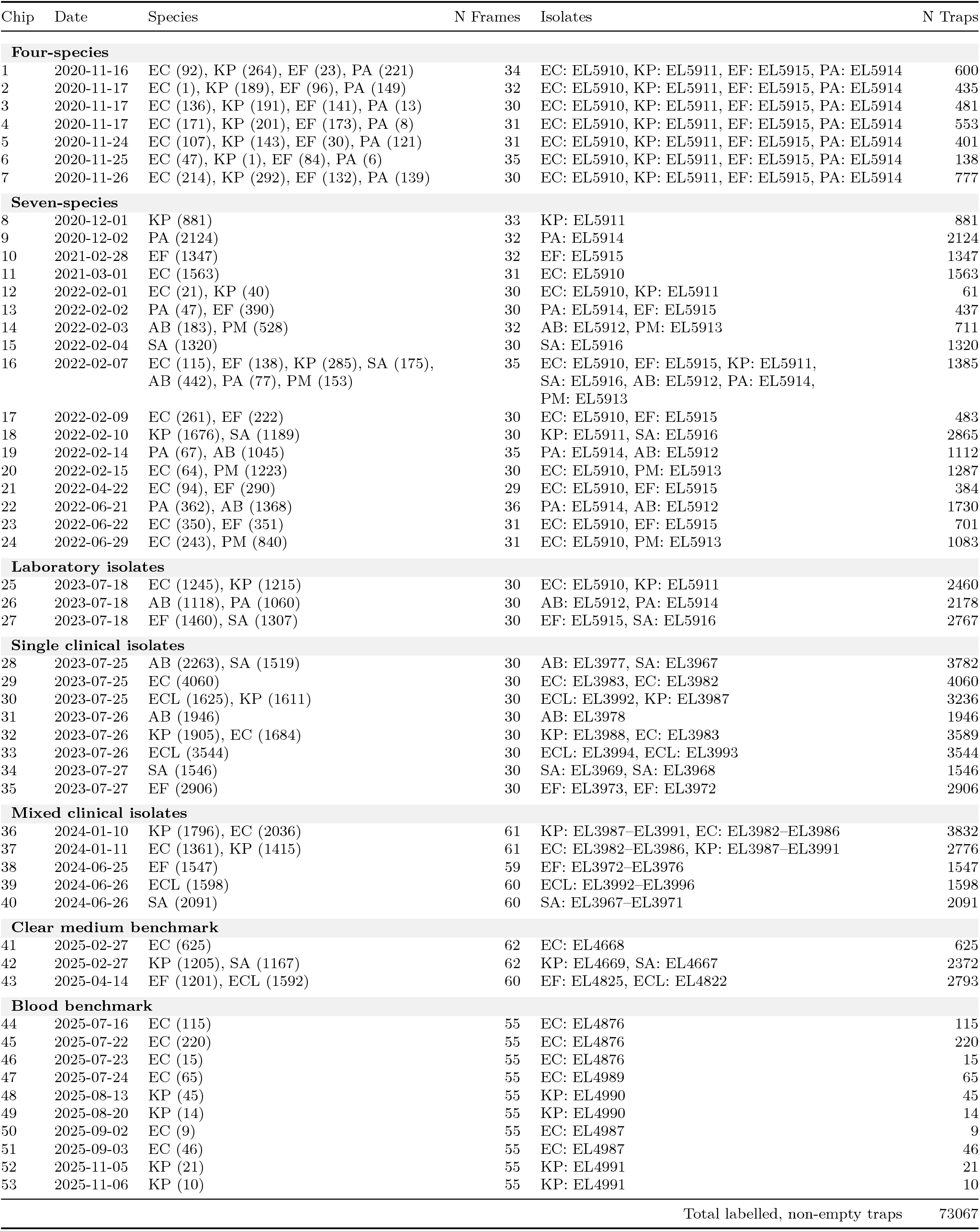
Full dataset inventory. Species abbreviations are EC (*Escherichia coli*), EF (*Enterococcus faecalis*), KP (*Klebsiella pneumoniae*), SA (*Staphylococcus aureus*), AB (*Acinetobacter baumannii*), PA (*Pseudomonas aeruginosa*), PM (*Proteus mirabilis*) and ECL (*Enterobacter cloacae*). N Frames is the number of phase-contrast frames per trap and N Traps the number of labelled, non-empty traps per chip. Numbers in parentheses give the number of traps per species. EL identifiers refer to Johan Elf’s laboratory collection, and strain and origin details are listed in Table 5. Dates are acquisition dates. From chip 25 onward, samples were unstained, and each chip section contained one isolate or a mixture of isolates of one species, listed as top, bottom. The Blood benchmark was imaged at one frame per minute and all other chips every two minutes. The final two frames of chips 41 and 42 were excluded so that all Clear-medium recordings used 60 frames. The Four-species dataset (chips 1–7) is from [10], and the Seven-species dataset (chips 8–24) is from [11].

**Table 5:** Strain and isolate origins. Each strain or isolate is listed once, using the EL identifiers of Johan Elf’s laboratory. DA identifiers refer to the strain collection of Dan I. Andersson’s group (Uppsala University), from which all clinical training isolates were obtained. Provider ID is the designation used by the originating laboratory or the strain designation. Laboratory strains are those described by Kandavalli et al. [8]. Chip assignments are listed in Table 4. Clinical source describes the original specimen, rather than the medium used in the microfluidic experiments. Year is reproduced as recorded in the collection records. *Sample origin:* Abbreviations are UUH (Uppsala University Hospital, Sweden), CCUG (Culture Collection University of Gothenburg, Sweden), EUCAST (European Committee on Antimicrobial Susceptibility Testing), KSH (Karolinska Institutet, Huddinge, Sweden), MAD (Hospital Universitario Ramón y Cajal, Madrid, Spain), TRO (University Hospital of North Norway, Tromsø, Norway) and CPH (Rigshospitalet, Copenhagen, Denmark). EL4876 is isolate C6 from Baltekin et al. [26]. The abbreviation n.s. means not stated in the collection records. The abbreviation n.a. means not applicable. A dash indicates that the information is unavailable.

| EL ID | Species | DA ID | Provider ID / strain | Sample origin | Clinical source | Year |
| --- | --- | --- | --- | --- | --- | --- |
| **Laboratory strains (chips 1–27)** |  |  |  |  |  |  |
| EL5910 | *E. coli* | DA4201 | K-12 MG1655 | Ref. [8] | n.a. | n.a. |
| EL5911 | *K. pneumoniae* | – | ATCC 13883 | Ref. [8] | n.a. | n.a. |
| EL5912 | *A. baumannii* | DA68153 | – | Ref. [8] | n.a. | n.a. |
| EL5913 | *P. mirabilis* | – | ATCC 29906 | Ref. [8] | n.a. | n.a. |
| EL5914 | *P. aeruginosa* | DA6215 | – | Ref. [8] | n.a. | n.a. |
| EL5915 | *E. faecalis* | – | ATCC 51299 | Ref. [8] | n.a. | n.a. |
| EL5916 | *S. aureus* | – | ATCC 29213 | Ref. [8] | n.a. | n.a. |
| **Clinical training isolates (chips 28–40)** |  |  |  |  |  |  |
| EL3967 | *S. aureus* | DA70300 | – | MAD | n.s. | n.s. |
| EL3968 | *S. aureus* | DA70488 | KSH-Sa3 | KSH | Superficial wound | n.s. |
| EL3969 | *S. aureus* | DA70518 | KSH-Sa18 | KSH | Blood | n.s. |
| EL3970 | *S. aureus* | DA70710 | Nr 20 | TRO | Wound | 2020 |
| EL3971 | *S. aureus* | DA70898 | – | CPH | n.s. | n.s. |
| EL3972 | *E. faecalis* | DA70208 | Nr 6 | MAD | n.s. | n.s. |
| EL3973 | *E. faecalis* | DA70222 | Nr 13 | MAD | n.s. | n.s. |
| EL3974 | *E. faecalis* | DA70472 | KSH-Efa17 | KSH | Blood | n.s. |
| EL3975 | *E. faecalis* | DA70602 | Nr 6 | TRO | Urine | 2018 |
| EL3976 | *E. faecalis* | DA70834 | – | CPH | n.s. | n.s. |
| EL3977 | *A. baumannii* | DA72716 | KSH-AB1 | KSH | n.s. | 2021 |
| EL3978 | *A. baumannii* | DA72717 | KSH-AB2 | KSH | n.s. | 2020 |
| EL3982 | *E. coli* | DA62888 | – | UUH | Blood | 2015 |
| EL3983 | *E. coli* | DA62890 | – | UUH | Blood | 2015 |
| EL3984 | *E. coli* | DA62894 | – | UUH | Blood | 2015 |
| EL3985 | *E. coli* | DA62896 | – | UUH | Blood | 2015 |
| EL3986 | *E. coli* | DA62898 | – | UUH | Blood | 2015 |
| EL3987 | *K. pneumoniae* | DA72206 | KSH-KP1 | KSH | n.s. | 2021 |
| EL3988 | *K. pneumoniae* | DA72207 | KSH-KP2 | KSH | n.s. | 2021 |
| EL3989 | *K. pneumoniae* | DA72208 | KSH-KP3 | KSH | n.s. | 2021 |
| EL3990 | *K. pneumoniae* | DA72209 | KSH-KP4 | KSH | n.s. | 2021 |
| EL3991 | *K. pneumoniae* | DA72210 | KSH-KP5 | KSH | n.s. | 2021 |
| EL3992 | *E. cloacae* | DA72734 | KSH-ENT1 | KSH | n.s. | 2021 |
| EL3993 | *E. cloacae* | DA72735 | KSH-ENT2 | KSH | n.s. | 2021 |
| EL3994 | *E. cloacae* | DA72736 | KSH-ENT3 | KSH | n.s. | 2021 |
| EL3995 | *E. cloacae* | DA72737 | KSH-ENT4 | KSH | n.s. | 2021 |
| EL3996 | *E. cloacae* | DA72738 | KSH-ENT5 | KSH | n.s. | 2021 |
| **Clear medium benchmark (chips 41–43)** |  |  |  |  |  |  |
| EL4668 | *E. coli* | – | ALB53 | CCUG | – | – |
| EL4669 | *K. pneumoniae* | – | 50878013 | CCUG | – | – |
| EL4667 | *S. aureus* | – | 6701 | CCUG | – | – |
| EL4825 | *E. faecalis* | – | CCUG 59212 | CCUG | – | – |
| EL4822 | *E. cloacae* | – | CCUG 6323 | CCUG | – | – |
| **Blood benchmark (chips 44–53)** |  |  |  |  |  |  |
| EL4876 | *E. coli* | – | C6 [26] | UUH | – | – |
| EL4989 | *E. coli* | – | 50857972 | EUCAST | – | – |
| EL4990 | *K. pneumoniae* | – | 3560B | EUCAST | – | – |
| EL4987 | *E. coli* | – | 19-597221 | EUCAST | – | – |
| EL4991 | *K. pneumoniae* | – | TU 7 | EUCAST | – | – |

In the four-species and seven-species datasets, bacterial fluorescent staining after the phase-contrast time-lapse provided species labels. This allowed multiple species to be loaded into the same chip, which may reduce overfitting to individual experiments. The seven-species dataset used combinatorial fluorescence in situ hybridization (FISH) originating from Kandavalli et al. [8], in which the fluorescence-staining procedure is also described in detail.

The new experiments did not use fluorescent staining for species identification. Instead, each separately loaded chip section contained one species, providing the species label. The new data consist of single laboratory isolates in chips 25–27, single clinical isolates in chips 28–35, and mixtures of clinical isolates of the same species in chips 36–40. Chips 41–43 also contain single clinical isolates, but these isolates were not used in training. After the final phase-contrast frame, autofluorescence images were acquired using 488-nm excitation. Autofluorescence is the naturally occurring fluorescence of molecules within the cells and requires no added fluorescent labels [14].

For the Clear medium dataset, bacterial cultures were grown overnight in Mueller–Hinton medium at 37 *^◦^*C, diluted into fresh medium and grown for a further two hours before loading. For the new clinical experiments in chips 28–43, cultures were diluted 1:100 and imaged at 30 *^◦^*C. Imaging was performed using a Nikon Ti2-E microscope with a 100*×* oil-immersion objective with a numerical aperture (NA) of 1.45 and a camera with 3.45 *µ*m pixels. Growth medium flowed continuously through the device while phase-contrast microscopy recorded bacterial growth.

The Blood dataset originated from blood processed using a protocol developed by Marino Miguélez et al. [4]. Donor blood was spiked with known bacterial isolates. Centrifugation, selective blood-cell lysis and volume reduction were followed by microfluidic loading. The bacteria were then grown in the device for 60 minutes before automated detection and subsequent imaging of positions containing bacteria. This growth period also allowed residual blood lysate to clear from some traps, providing clearer images for downstream analysis. The lower bacterial concentrations resulted in fewer occupied traps. Chips 44–53 contained 560 non-empty traps from three *E. coli* and two *K. pneumoniae* isolates.

### 3.2 Imaging and preprocessing

Individual traps were extracted from the raw phase-contrast image sequences using the cropping and registration pipeline used in our previous studies. For the four-species and seven-species datasets, the images available in the replication packages on Zenodo were adapted. Chips 25–43 used the cropping and stabilization pipeline from the seven-species study, including translation registration using StackReg [27]. Chips 44–53 used the improved pipeline published in our bacterial detection study [4].

Two modifications were made. Previously, trap images with a width of 52 pixels were used. Here, five pixels from each side were cropped because these contained the trap walls, leaving a width of 42 pixels. In our previous studies, we noticed good performance despite substantial downsampling. This was possibly due to the images being oversampled. The Nikon Ti2-E microscope used a 100*×* oil-immersion objective with an NA of 1.45 and a camera with 3.45 *µ*m pixels, meaning that each pixel covered 0.0345 *µ*m of the sample. Assuming a wavelength of 550 nm, the estimated optical resolution was approximately 0.23 *µ*m. Using at least two pixels per resolvable detail, following the Nyquist sampling criterion, the cropped image width of 1.45 *µ*m required approximately 13 pixels. The trap images were therefore downsampled to 16 pixels per trap, retaining more than two pixels per resolvable detail while matching the transformer’s patch size.

### 3.3 Model and training

The vision-transformer experiments were repeated 31 times using independent random seeds. The autofluorescence experiments also used 31 seeds, with the same seeds paired across the three input conditions. The representative run was selected by averaging the Clear medium and Blood macro-F1 scores for each run and choosing the run with the median average score. The confusion matrices and ROC curves in Figs 2 and 3 show this same model.

Our proposed model is based on the standard ViT Small pretrained using the DINOv3 self-supervised methodology, with 16 *×* 16-pixel patches [13]. Training consists of self-supervised adaptation followed by supervised fine-tuning, both using chips 1–40. Self-supervised adaptation is performed for one epoch and supervised fine-tuning for two epochs, both over non-overlapping windows of ten consecutive frames. Both stages use montages of ten frames, each 384 *×* 16 pixels, giving an input size of 384 *×* 160 pixels. The whole strategy is shown in Fig 7. All LoRA adaptations use rank 8 and a scaling parameter of 8, giving a scaling factor of one. Both stages use a batch size of 256 and AdamW with a learning rate of 10*^−^*^3^, a cosine schedule that decays to zero and no weight decay.

**Fig 7.**
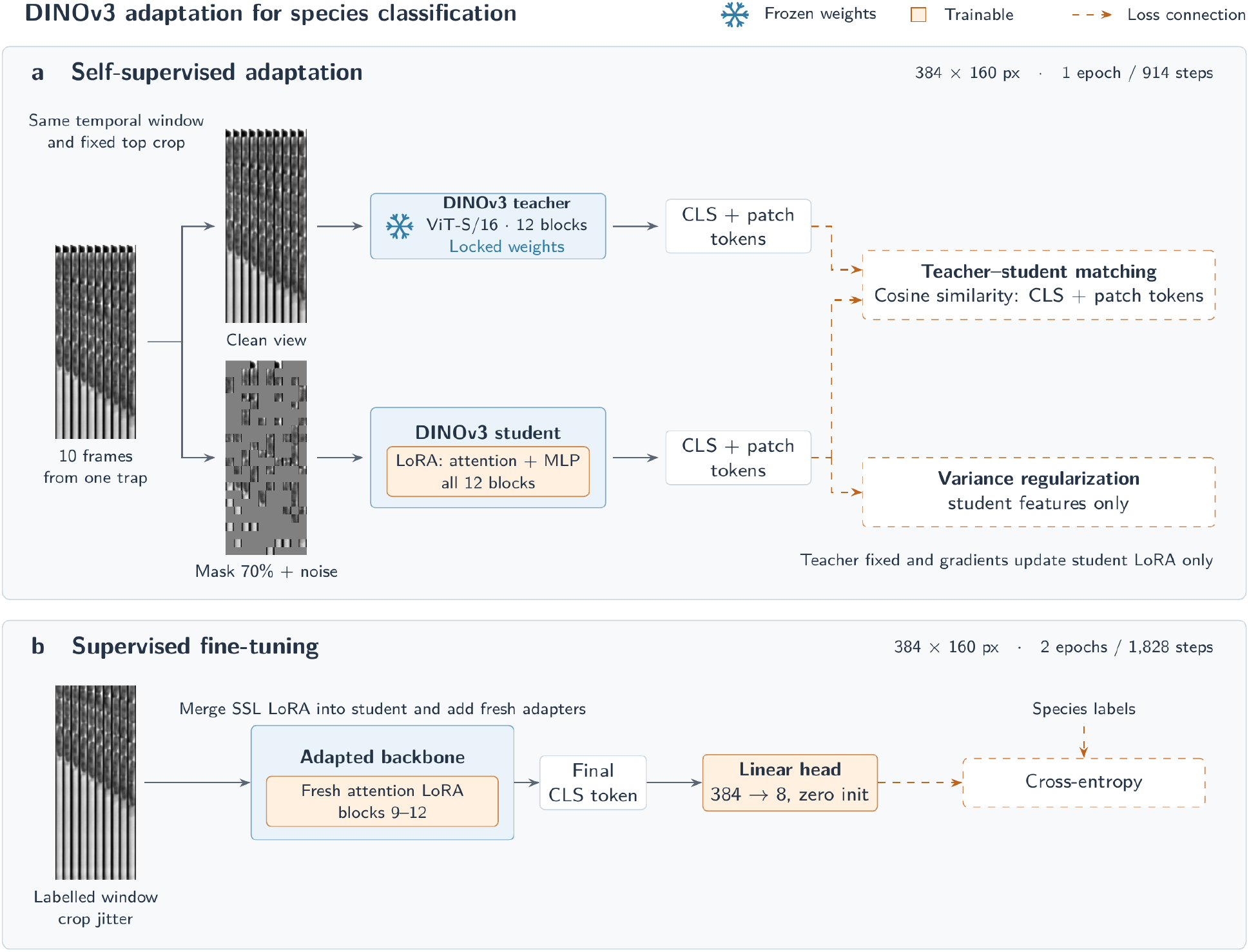
DINOv3 adaptation for bacterial species classification. (a) Self-supervised adaptation with a fixed teacher and a student trained through LoRA adapters using one clean teacher view and one corrupted student view of the same temporal window. Teacher–student matching uses CLS and patch tokens, with variance regularization applied to student features. (b) Supervised fine-tuning after merging the learned SSL LoRA weights into the student and adding fresh attention LoRA adapters and a linear classification head.

The original single-channel grayscale phase-contrast image intensities are divided by 65,535, the maximum value of the raw uint16 images, and replicated across three channels to match the input format of the model (which was pretrained on natural images). Each channel is then normalized using the ImageNet means (0.485, 0.456 and 0.406) and standard deviations (0.229, 0.224 and 0.225). Patch masking (used during SSL), frame erasing (used during supervised fine-tuning) and padding of missing frames (when evaluating time-lapses with fewer than ten frames) are applied after normalization, setting all three channels to zero at the affected positions.

#### 3.3.1 Self-supervised adaptation

The input to the SSL stage consists of ten consecutive frames, each cropped to a fixed height of 384 pixels from the top of each trap and concatenated into a montage, as described in the preceding section and shown on the left in Fig 7a. First, the teacher, a DINOv3 model with frozen weights, receives the clean montage and produces CLS and patch tokens. The CLS token is a feature vector describing the whole montage, while the patch tokens are feature vectors describing each of its 16 *×* 16-pixel input patches [28].

Then the student, a copy of the teacher with trainable parameters, receives a corrupted copy of the same montage. To create this corrupted copy, Gaussian pixel noise with a mean of zero and a standard deviation of 0.005 is added to the grayscale input scaled to [0, 1], without clipping. This noise is added before the image is replicated across three channels and ImageNet-normalized, as described in the preceding section. Patch masking is then applied by independently setting each 16 *×* 16-pixel patch to zero in all three channels with a probability of 70%. These patches follow the transformer model’s patch grid, as shown on the left in Fig 7a.

The student has LoRA adaptations [21] to the attention and feed-forward projections in all 12 transformer blocks. The student is trained to match the teacher’s CLS and patch tokens using cosine similarity. Let *B* denote the batch size, *P* the number of patches per montage, and *D* the number of components in each feature vector. Here, *B* = 256, *P* = 240, and *D* = 384. Student and teacher features are denoted by *s* and *t*, respectively. The matching losses are

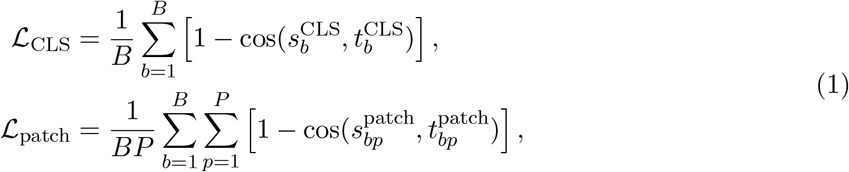

where cos(*·, ·*) denotes cosine similarity. The CLS loss is averaged over the batch, while the patch loss is averaged over all patches in the batch.

Variance regularization is applied separately to the student’s CLS and patch features, following the variance component of VICReg [29]. For a collection *Z* of *N* student feature vectors, the mean and standard deviation of each component *d* are

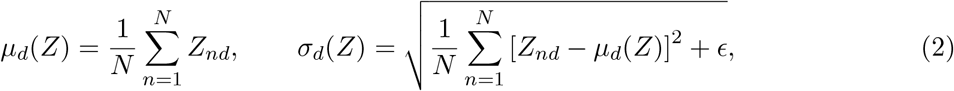

where *ɛ* = 10*^−^*^4^ ensures numerical stability. These calculations use the features before unit-length normalization. The regularization penalizes standard deviations below one, averaged over the *D* components:

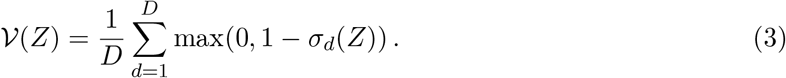

For CLS features, *S*_CLS_ contains the *B* CLS vectors in the batch. For patch features, *S*_patch_ contains all *BP* patch vectors in the batch. The total SSL loss gives equal weight to all four terms:

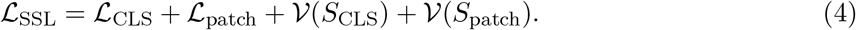

#### 3.3.2 Supervised species classification

After adaptation, the SSL LoRA updates are merged into the student weights. Fresh rank-8 LoRA updates are added to the attention projections of the last four blocks, together with a linear classifier using the final CLS token to predict eight species. The classifier is initialized to zero and trained jointly with the new LoRA parameters from the first optimizer step.

During supervised fine-tuning, the vertical crop offset is sampled uniformly from zero to 100 pixels. Each training window has a 10% probability of Gaussian blur being applied separately to each frame before ImageNet normalization, using a standard deviation of 2 pixels, a 13 *×* 13 kernel and reflection padding. Each training window also has a 10% probability of frame erasing. The number of erased frames is sampled uniformly from one to nine, and that many consecutive frames are erased from the end of the montage by setting all three ImageNet-normalized channels to zero.

The classifier uses cross-entropy without class weights.

#### 3.3.3 Baselines and ablations

The supervised LoRA baseline omits the SSL stage but retains the same classifier and supervised training procedure. The full fine-tuning baseline updates all backbone weights, using a backbone learning rate of 10*^−^*^4^, layer-wise learning-rate decay of 0.25 and a classifier learning rate of 10*^−^*^3^.

Backbone comparisons use DINO, DINOv2 and DINOv3. For DINOv2, one column is cropped from each side of each prepared frame to fit its 14 *×*14-pixel patches, and the image height is padded by reflection to a multiple of 14. Chips 1–40 (66,717 traps) were used for both self-supervised adaptation and supervised training. This includes chip 16, which served as a benchmark in the earlier seven-species study. Neither fitting stage uses Clear medium (chips 41–43) or Blood (chips 44–53). These datasets contain different isolate identifiers from the training set but informed method development, so they serve as development benchmarks rather than independent external validation sets.

### 3.4 Training and inference times

Training time averaged 4.0 minutes per seed on a single NVIDIA GH200 GPU for the SSL and supervised stages combined. Training was fast because only one epoch was used for SSL and two epochs for supervised fine-tuning, with LoRA training only a small number of additional parameters. Most computation time was spent evaluating simulated numerical apertures and different observation times. Model-forward inference time averaged 0.143 ms per ten-frame montage across the 31 runs when processed in batches of 256, excluding preprocessing, data transfer and logit aggregation, suggesting that model inference would add little time compared with bacterial growth and image acquisition.

### 3.5 Evaluation

With fewer than ten frames, the remaining frame positions were masked by setting all three normalized channels to zero. For longer recordings, ten-frame windows were evaluated with a stride of one, and their logits were averaged to obtain one prediction per trap. Clear medium was evaluated using up to 60 frames (118 minutes), and Blood using up to 55 frames (54 minutes).

Performance was measured using macro-F1 calculated from the confusion matrix, averaging the F1 scores of the species present in each evaluation set. All eight model outputs remained available, so predictions of species absent from the evaluation set counted as errors. ROC curves and AUC were calculated using a one-versus-rest approach. Results are reported as mean *±* sample standard deviation across training seeds.

Lower NA was approximated by filtering each frame in the Fourier domain. The filter was circularly symmetric, with coefficients decreasing smoothly from 1 at the centre to 0 at the target cutoff, progressively removing fine spatial detail. This profile was calculated from a standard optical model for a circular aperture, accounting for the original NA of 1.45 and assuming an illumination wavelength of 550 nm [30]. Target NA values ranged from 1.4 to 0.1, and the same trained model was evaluated without retraining. This approximated reduced spatial resolution rather than the complete image formation of a lower-NA phase-contrast microscope.

## 4 Discussion

The results suggest that bacterial species identification from phase-contrast time-lapse images is possible for clinical isolates grown in clear medium and recovered from spiked blood. The confusion matrices and ROC curves indicate that the rod-shaped species, *E. coli*, *K. pneumoniae* and *E. cloacae*, were more difficult to distinguish than *E. faecalis* and *S. aureus* (Figs 2–3). Confusion was particularly apparent for *E. cloacae* predicted as *E. coli* and *K. pneumoniae* predicted as *E. cloacae*. This is reasonable given their similar appearance.

Autofluorescence provided additional information for species identification, although the improvement over phase contrast alone was small (mean macro-F1 from 62.6% to 65.9%). Acquiring autofluorescence images also requires detection equipment and an additional imaging step. When phase-contrast images are already available for monitoring bacterial growth, using these images alone may therefore be a simpler approach.

Self-supervised adaptation was important for performance in the present study, particularly on Blood. It raised mean macro-F1 from 81.5% to 93.6% on Clear medium and from 61.3% to 87.2% on Blood. Both sets consisted of clinical isolates not seen during training and came from separate experiments recorded 8 to 17 months after the latest training experiments, reducing the risk of data leakage between experiments (which could arise if experiments were performed close together and grouped by species). The two validation sets were also collected by two different researchers. Blood additionally tested bacteria recovered from spiked blood, a different sample condition from clear medium. That the method worked on both suggests that it is not limited to one set of isolates or one sample condition. However, the evaluation datasets were examined during method development and therefore effectively served as validation sets. Confirming the results independently on new clinical isolates not seen during method development is an important next step and will help establish how well the method generalizes.

The limited performance of the previous CNN method (macro-F1 66.6% on Clear medium after retraining) also shows that generalization between different bacterial isolates was not straightforward. With a larger and more varied training dataset, supervised training alone might perform better, and the benefit of SSL adaptation could be different.

Performance improved as more frames became available, but useful species information was already present in shorter recordings. Performance was also retained after moderate removal of spatial detail, suggesting that lower-resolution imaging could be worth investigating experimentally. In a clinical application, decisions would be made from the sample as a whole rather than from individual traps. Combining predictions across traps could improve reliability, but this requires evaluation on independent experiments and patient samples.

In summary, this study, together with our previous studies on bacterial detection [4] and species identification [10, 11], leads us to conclude that spatiotemporal features provide valuable information for deep learning-based pathogen detection and identification. These approaches could help transform infection care by enabling faster, better-informed treatment decisions and may extend to other pathogens, opening new possibilities for rapid, label-free diagnostics.

## Acknowledgements

This work was supported by the Swedish Foundation for Strategic Research (SSF), grant ARC19-0016, awarded to Carolina Wählby and Johan Elf. The funders had no role in designing the study. Computations were carried out on the Alvis GPU cluster at Chalmers University of Technology and the GPU partition of the Arrhenius supercomputer at Linköping University, through the National Academic Infrastructure for Supercomputing in Sweden (NAISS), with funding from the Swedish Research Council. Additionally, computations were carried out on the Berzelius AI supercomputer, operated by NAISS and funded by the Knut and Alice Wallenberg Foundation.

## Author information

Vinodh Kandavalli and M. Henar Marino Miguélez contributed equally to this work.

## Author contributions

E.H. designed all deep learning experiments, developed the software and the two-stage training procedure, performed the computational experiments and data analysis, prepared the figures, and wrote and revised the manuscript. V.K. conducted the Clear medium experiments, and M.M. conducted the Blood experiments. Both datasets were generated in J.E.’s laboratory. J.E. and C.W. acquired funding and revised the manuscript.

## Corresponding author

Correspondence to Erik Hallström.

## Code and data availability

A replication package is available at [24], containing the experiment results, the phase-contrast image data and the code to reproduce all deep learning experiments in this paper in addition to generating all plots. It also includes the trained deep learning model from the representative median run used to generate the confusion matrices presented in the paper. The code and experiment results described in S1 and S2 Documents are also included. The autofluorescence data, the sample gallery and the dependency Apptainer image are available at [25].

## Supporting information

### Supporting documents

**S1 Document.** Full evaluation of previous models on new laboratory and clinical data, including confusion matrices and the clinical-training experiment.

**S2 Document.** Autofluorescence, phase-contrast and combined-input classification: experimental setup, seed-level summaries and confusion matrices.

**S3 Document.** Blood dataset results by clinical isolate and chip, using the same representative run as the main confusion matrices.

### Supporting figures

#### Sample examples

Each image shows the full recording of one trap at NA 1.45 (unfiltered), with the frames side by side from left to right and the frame number above each frame. Frames were acquired every 2 min in Clear medium (118 min in total) and every minute in Blood (54 min in total). Frames beyond the 60 used for evaluation, where present, are shown dimmed.

**S1 Fig. Clear medium validation sample: *E. coli*, Isolate EL4668.**

**S2 Fig. Clear medium validation sample: *K. pneumoniae*, Isolate EL4669. S3 Fig. Clear medium validation sample: *E. faecalis*, Isolate EL4825.**

**S4 Fig. Clear medium validation sample: *S. aureus*, Isolate EL4667.**

**S5 Fig. Clear medium validation sample: *E. cloacae*, Isolate EL4822.**

**S6 Fig. Blood validation sample: *E. coli*, Isolate EL4876.**

**S7 Fig. Blood validation sample: *K. pneumoniae*, Isolate EL4991.**

#### Misclassification examples

In each image, the outline marks the part of the recording that the model saw at the stated time, namely the frames up to that time and the upper 384 pixels of each frame. The rest of the recording is dimmed and shown for context. The predicted class probabilities (%) are listed below each image, removing classes below 1%.

**S8 Fig. At 32 min in clear medium, the model misclassified *E. coli* as *S. aureus* with low confidence (35%).** Only a few cells are visible near the top of the trap, and their outlines are too unclear to distinguish rods from cocci.

**S9 Fig. At 15 min in Blood, the model misclassified *E. coli* as *P. mirabilis* with high confidence (98%).** *E. coli* and *P. mirabilis* are both rods, and the cell outlines are difficult to distinguish in this early recording.

**S10 Fig. At 30 min in clear medium, the model misclassified *S. aureus* as *E. faecalis* with high confidence (91%).** *S. aureus* and *E. faecalis* are both cocci, and the closely packed rounded cells make them difficult to distinguish by shape.

**S11 Fig. At 15 min in Blood, the model misclassified *K. pneumoniae* as *E. cloacae* with high confidence (85%).** *K. pneumoniae* and *E. cloacae* are both rods, with little visible growth or detail to distinguish them at this timepoint.

**S12 Fig. At 40 min in clear medium, the model misclassified *E. coli* as *E. faecalis* with low confidence (32%).** Smaller rounded cells appear alongside longer cells, suggesting a mixed population and possible contamination.

**S13 Fig. At 118 min in clear medium, the model misclassified *E. cloacae* as *E. faecalis* with high confidence (99%).** The cells near the top of the trap appear rounded, like cocci rather than the rod-shaped *E. cloacae* stated on the label, which suggests contamination by a coccus species.

**S14 Fig. At 118 min in clear medium, the model misclassified *E. faecalis* as *E. cloacae* with high confidence (92%).** The cells appear elongated rather than coccal as expected for *E. faecalis*, suggesting possible contamination by a rod-shaped species.

**S15 Fig. At 118 min in clear medium, the model misclassified *E. faecalis* as *K. pneumoniae* with low confidence (44%).** The trap is mostly empty, with a few cells near the top and very little visible growth.

**S16 Fig. At 54 min in Blood, the model misclassified *K. pneumoniae* as *E. coli* with high confidence (98%).** *K. pneumoniae* and *E. coli* are both rods, and the cells look similar.

**S17 Fig. At 54 min in Blood, the model misclassified *K. pneumoniae* as *E. coli* with low confidence (42%).** *K. pneumoniae* and *E. coli* are both rods with similar outlines, and the model scores are nearly tied.

**S18 Fig. At 118 min in clear medium, the model misclassified *E. coli* as *P. aeruginosa* with moderate confidence (61%).** *E. coli* and *P. aeruginosa* are both rods. Only a few cells are visible, with little growth, and they appear to have detached from the top of the trap.

**S19 Fig. At 54 min in Blood, the model misclassified *E. coli* as *K. pneumoniae* with low confidence (44%).** *E. coli* and *K. pneumoniae* are both rods with similar shapes, and the model scores are nearly tied.

**S20 Fig. At 54 min in Blood, the model misclassified *E. coli* as *K. pneumoniae* with high confidence (89%).** *E. coli* and *K. pneumoniae* are both rods, and the cells grew slowly.

**S21 Fig. At 54 min in Blood, the model misclassified *E. coli* as *K. pneumoniae* with high confidence (91%).** *E. coli* and *K. pneumoniae* are both rods with similar shapes, making them difficult to distinguish.

**S22 Fig. At 118 min in clear medium, the model misclassified *E. coli* as *K. pneumoniae* with high confidence (93%).** *E. coli* and *K. pneumoniae* are both rods with similar shapes, making them difficult to distinguish.

**S23 Fig. At 118 min in clear medium, the model misclassified *E. coli* as *K. pneumoniae* with moderate confidence (58%).** The trap remains mostly empty, with one unusually long cell showing slight growth toward the end.

**S24 Fig. At 118 min in clear medium, the model misclassified *K. pneumoniae* as *E. coli* with high confidence (85%).** *K. pneumoniae* and *E. coli* are both rods with similar shapes, making them difficult to distinguish.

**S25 Fig. At 118 min in clear medium, the model misclassified *K. pneumoniae* as *E. coli* with high confidence (94%).** *K. pneumoniae* and *E. coli* are both rods with similar shapes, making them difficult to distinguish.

**S26 Fig. At 118 min in clear medium, the model misclassified *K. pneumoniae* as *E. coli* with moderate confidence (53%).** One cell, followed by a few cells, shows very little growth near the top of the trap, leaving little detail to distinguish *K. pneumoniae* from *E. coli*.

**S27 Fig. At 118 min in clear medium, the model misclassified *E. cloacae* as *E. coli* with low confidence (44%).** The trap is initially almost empty, then cells appear farther down and shift position, making the species difficult to distinguish.

**S28 Fig. At 118 min in clear medium, the model misclassified *E. faecalis* as *E. coli* with low confidence (39%).** Only a few rounded cells are visible near the top, with very little growth. Their coccal appearance differs from the rod-shaped prediction.

**S29 Fig. At 118 min in clear medium, the model misclassified *E. faecalis* as *E. coli* with low confidence (47%).** A small group of cocci appears near the top of the trap, while rods are visible farther down, suggesting possible contamination.

**S30 Fig. At 118 min in clear medium, the model misclassified *E. faecalis* as *S. aureus* with moderate confidence (76%).** A few cocci remain near the top with no visible growth, making the two coccal species difficult to distinguish.

**S31 Fig. At 118 min in clear medium, the model misclassified *K. pneumoniae* as *S. aureus* with low confidence (42%).** The trap appears empty and may have been missed during cleaning.

**S32 Fig. At 118 min in clear medium, the model misclassified *K. pneumoniae* as *A. baumannii* with moderate confidence (56%).** Cocci appear near the top of the trap despite the rod-shaped species label, suggesting possible contamination.

#### Simulated NA comparison

S33–S38 Figs show time-lapse montages of the full trap recordings at simulated NA 0.70 or 0.30. The original images were acquired at NA 1.45.

**S33 Fig. Clear medium validation sample: *E. coli*, Isolate EL4668, simulated NA 0.70.** Same trap as in S1 Fig.

**S34 Fig. Clear medium validation sample: *E. coli*, Isolate EL4668, simulated NA 0.30.** Same trap as in S1 Fig.

**S35 Fig. Blood validation sample: *E. coli*, Isolate EL4876, simulated NA 0.70.** Same trap as in S6 Fig.

**S36 Fig. Blood validation sample: *E. coli*, Isolate EL4876, simulated NA 0.30.** Same trap as in S6 Fig.

**S37 Fig. Clear medium validation sample: *S. aureus*, Isolate EL4667, simulated NA 0.70.** Same trap as in S4 Fig.

**S38 Fig. Clear medium validation sample: *S. aureus*, Isolate EL4667, simulated NA 0.30.** Same trap as in S4 Fig.

#### Multiple samples

S39–S41 Figs show the final frame of the growth overview of 60 traps across both datasets and all species and isolates, at 54 min in Blood and 118 min in Clear medium.

**S39 Fig. Multiple samples from both datasets, NA 1.45 unfiltered.**

**S40 Fig. Multiple samples from both datasets, simulated NA 0.70.**

**S41 Fig. Multiple samples from both datasets, simulated NA 0.30.**

### Supporting videos

#### Multiple samples

S1–S3 Videos show growth in 60 traps across both datasets and five species. Images were acquired every minute in Blood and every two minutes in Clear medium.

**S1 Video. Multiple samples from both datasets, NA 1.45 unfiltered.**

**S2 Video. Multiple samples from both datasets, simulated NA 0.70.**

**S3 Video. Multiple samples from both datasets, simulated NA 0.30.**

#### Individual isolates at different simulated numerical apertures

S4–S10 Videos each show one isolate across all 15 NA levels, from NA 1.45 unfiltered to simulated NA 0.10.

**S4 Video. *E. coli* in Clear medium.** Isolate EL4668.

**S5 Video. *K. pneumoniae* in Clear medium.** Isolate EL4669.

**S6 Video. *E. faecalis* in Clear medium.** Isolate EL4825.

**S7 Video. *S. aureus* in Clear medium.** Isolate EL4667.

**S8 Video. *E. cloacae* in Clear medium.** Isolate EL4822.

**S9 Video. *E. coli* in Blood.** Isolate EL4876.

**S10 Video. *K. pneumoniae* in Blood.** Isolate EL4991.

