## Supplementary material for "Label-free identification of bacterial species using adapted vision transformers": S1 Document

### S1 Document: Testing the best models from the four-species and seven-species papers on new data

In our two previous studies, we trained models to classify four[1] and seven[2] bacterial species, respectively. However, both studies used a single laboratory strain per species in both training and testing. Since clinical isolates of the same species can differ greatly in appearance and growth, performance on these laboratory strains may not reflect performance on clinical isolates. R(2+1)D Video ResNet[3] achieved the best performance in both studies. However, because the reported scores were also used to select the best models, they effectively served as validation scores. To obtain an unbiased test result for the selected models, we needed to evaluate them on a completely new dataset, which became possible in this study. In this study, we collected data from both laboratory and clinical isolates. We therefore started by testing how well our previous models transferred to these new datasets, without further training.

**Input preparation.** We used the software available in the replication package of the seven-species article. The seven-species model was evaluated using its standard tiled crops of 114 × 52 pixels, while the four-species model used a single top crop of 352 × 52 pixels. For the four-species model, we normalised the input as described in the original four-species article. Both models were evaluated using the first 30 frames of each time-lapse.

##### Task 1: Testing previous models on new laboratory isolates

In this study, we collected new data from laboratory isolates of six species on chips 25–27. No new data were collected for *P. mirabilis*. We therefore tested the four-species model on all four species it was trained to classify, and the seven-species model on six of its seven species. Additionally, we tested the four-species model on the test set from the seven-species study, excluding traps containing *P. mirabilis*, *S. aureus*, or *A. baumannii*, since these species were not represented among its outputs. As shown in Table 1, both models performed well on the new laboratory data, but the four-species model performed poorly on the seven-species study's test set. This is consistent with our observation that the species in this test set were harder to distinguish, suggesting that transfer between experiments can be challenging.

[1] *Label-free deep learning-based species classification of bacteria imaged by phase-contrast microscopy.* doi:10.1371/journal.pcbi.1011181.

[2] *Rapid label-free identification of seven bacterial species using microfluidics, single-cell time-lapse phase-contrast microscopy, and deep learning-based image and video classification.* doi:10.1371/journal.pone.0330265.

[3] Tran et al. (2018). *A Closer Look at Spatiotemporal Convolutions for Action Recognition.* CVPR. Original paper.

| Model | Test data | Species | Traps | Macro F1 (%) |
|---|---|---|---|---|
| Four-species | Seven-species study test set, chip 16 | 4 | 623 | 53.7 |
| Four-species | New laboratory | 4 | 4,984 | 89.0 |
| Seven-species | New laboratory | 6 | 7,410 | 92.3 |

Table 1: Transfer to laboratory data. Macro F1 averages the per-species F1 scores over the species present in each test, giving each species equal weight. The models are evaluated on different species sets.

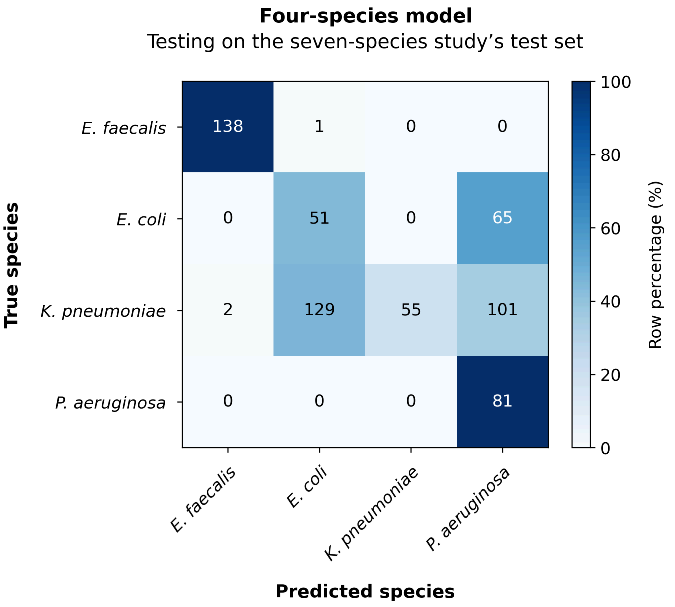


Figure 1: Four-species model on the seven-species study's test set (chip 16), restricted to its four supported species. Rows show actual species and columns show predictions. Cells show trap counts.

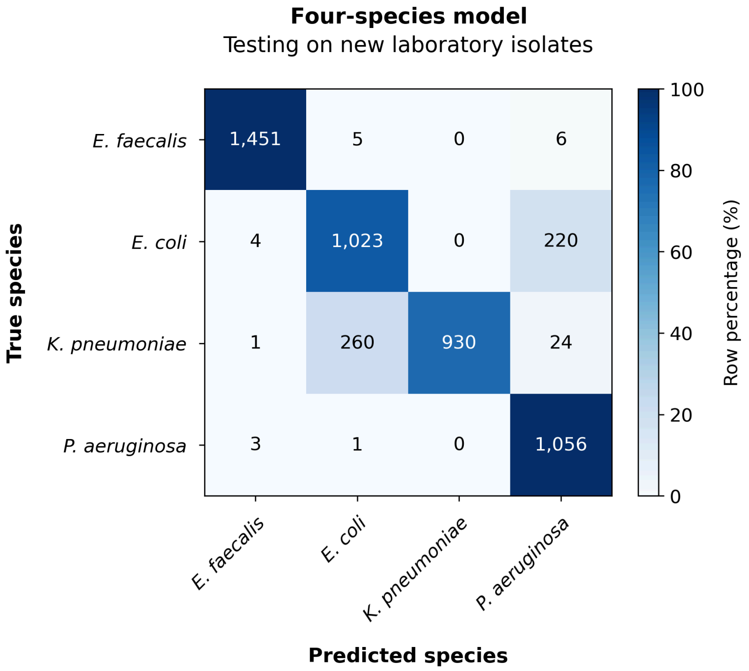


Figure 2: Four-species model on new laboratory data from chips 25-27, using all four supported species. Cells show trap counts. Colours show percentages within each actual-species row.

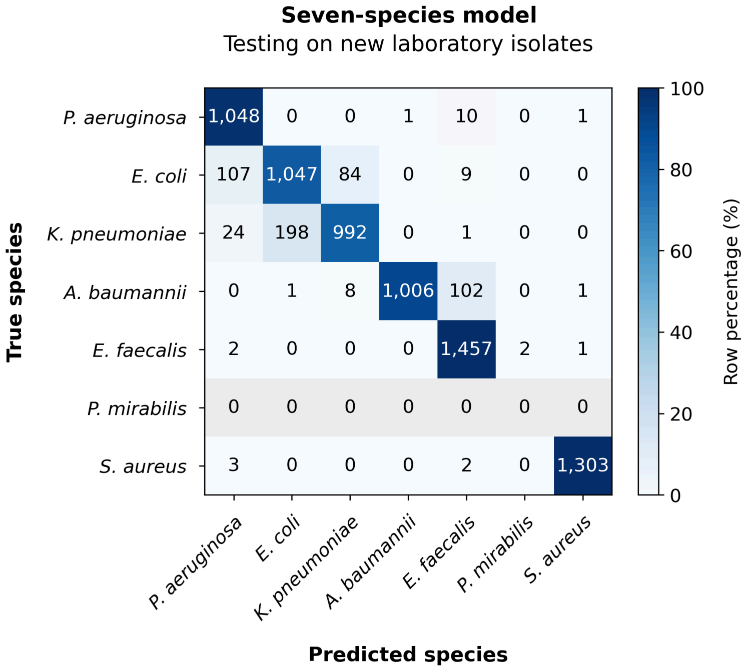


Figure 3: Seven-species model on new laboratory data from chips 25-27, using six species. All seven outputs are retained. Grey rows indicate species with no true examples. Cells show trap counts. Colours show percentages within each actual-species row.

In the chip 16 test, recall was 44.0% for *E. coli* and 19.2% for *K. pneumoniae*, compared with 99.3% for *E. faecalis* and 100% for *P. aeruginosa*.

##### Task 2: Comparing the performance of previous models on laboratory versus clinical isolates

In our new study, chips 28–40 contained clinical isolates. We compared how well our previous models performed on the new laboratory isolates in chips 25–27 versus the new clinical isolates. We were not able to obtain clinical isolates of *P. aeruginosa*, so there were three common species for the four-species model and five common species for the seven-species model. Additionally, the new clinical data contained *E. cloacae*, which was excluded because neither model was trained to classify this species.

| Model | Test data | Species | Traps | Macro F1 (%) |
|---|---|---|---|---|
| Four-species | New laboratory | 3 | 3,924 | 89.0 |
| Four-species | New clinical | 3 | 20,396 | 74.8 |
| Seven-species | New laboratory | 5 | 6,350 | 92.2 |
| Seven-species | New clinical | 5 | 29,768 | 62.3 |

Table 2: Matched laboratory and clinical comparisons. Each model uses the same species in both groups. Macro F1 averages the per-species F1 scores over those species.

The macro F1 score decreased from 89.0% on laboratory isolates to 74.8% on clinical isolates for the four-species model, and from 92.2% to 62.3% for the seven-species model. Both models therefore performed worse on the clinical isolates. This suggests that their performance on laboratory isolates did not fully transfer to clinical isolates. However, differences between experiments may also have contributed to the decrease.

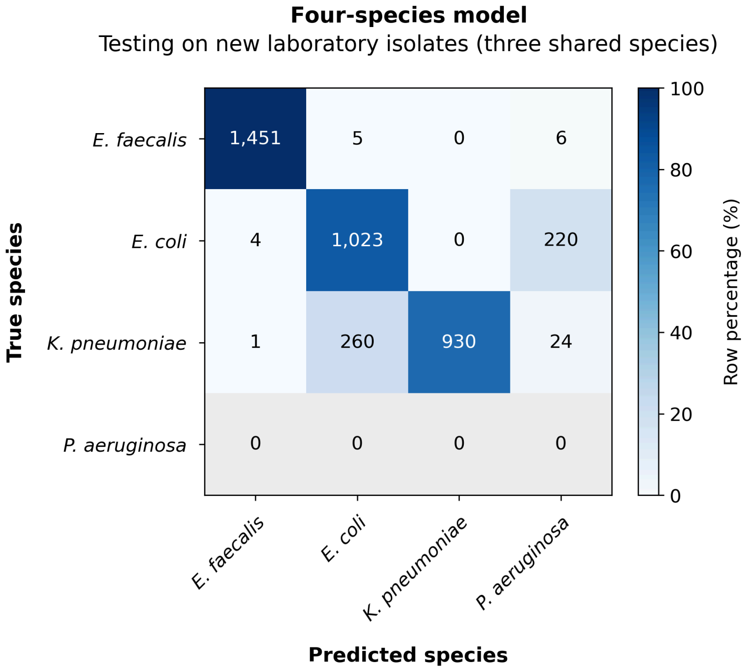


Figure 4: Four-species model on new laboratory data, restricted to the three species shared with the clinical comparison. All four outputs are retained. Cells show trap counts. Colours show row percentages, with recall on the diagonal. Grey rows indicate species with no true examples.

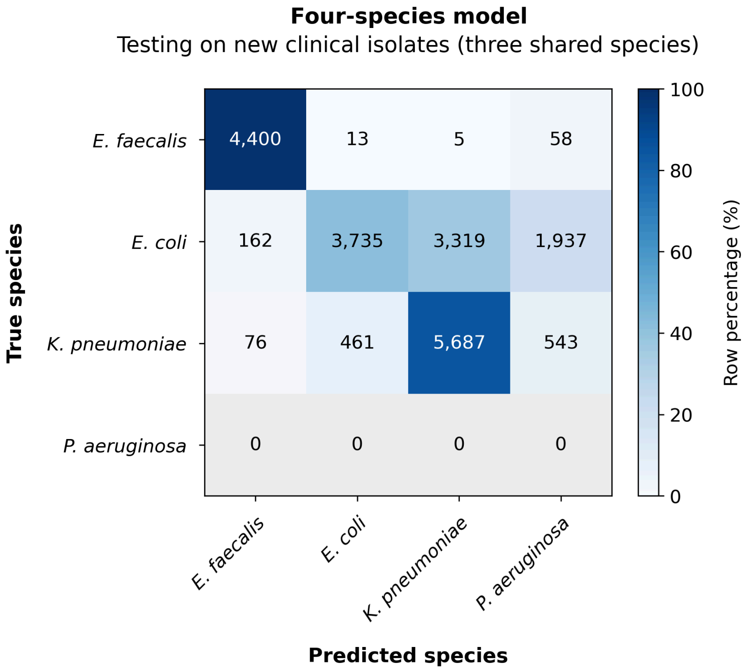


Figure 5: Four-species model on pooled clinical data from chips 28-40, restricted to the same three species as Figure 4. Unsupported true species are excluded. Cells show trap counts. Colours show row percentages, with recall on the diagonal.

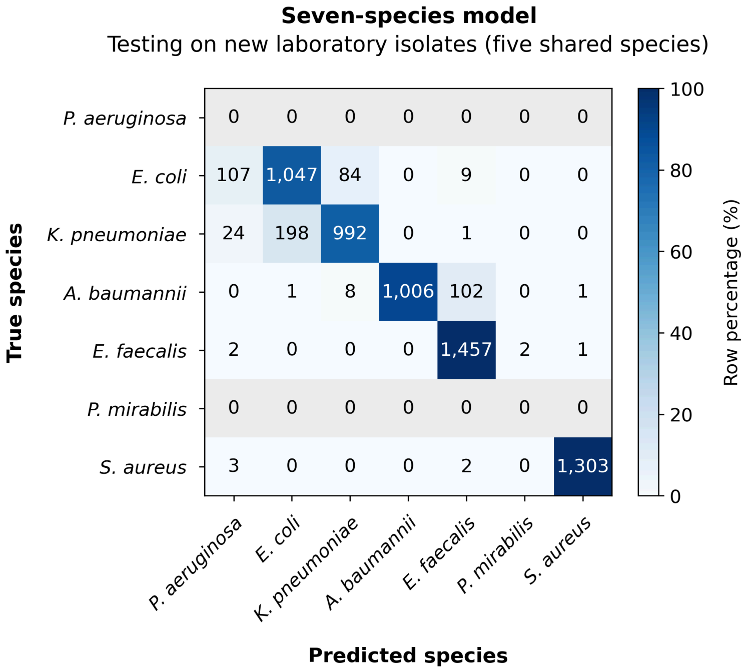


Figure 6: Seven-species model on new laboratory data, restricted to the five species shared with the clinical comparison. All seven outputs are retained. Cells show trap counts. Colours show row percentages, with recall on the diagonal. Grey rows indicate species with no true examples.

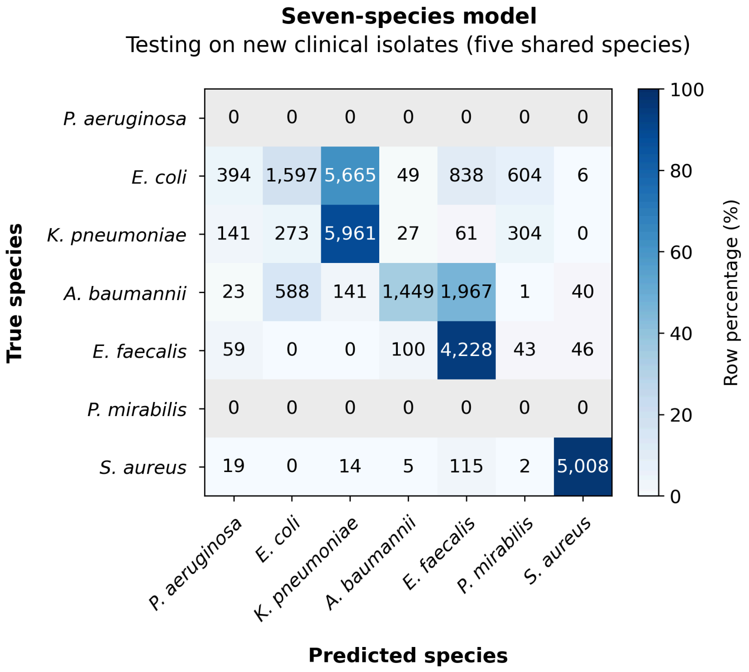


Figure 7: Seven-species model on pooled clinical data from chips 28-40, restricted to the same five species as Figure 6. Unsupported true species are excluded. Cells show trap counts. Colours show row percentages, with recall on the diagonal.

##### Task 3: Training Video ResNet on clinical isolates

We next tested whether the previous Video ResNet approach could classify new clinical isolates after training on clinical data. We used the seven-species study's R(2+1)D-18 model and training recipe, with pretrained initialisation, 114 × 52 pixel crops and 10-frame training clips. Training used chips 28-40. We increased the quality-based cap from 2,600 to 4,000 traps per species, giving 24,000 training traps across six species, including *E. cloacae*. We trained for 150 epochs with one random seed (557), and evaluated the final model using the first 30 frames and summed model scores per trap.

The clinical isolates in chips 41-43 (the clear medium validation set) were held out for testing, containing five species. All six model outputs were retained, although no *A. baumannii* traps were present in the test set. The final macro F1 score was 66.6% (Table 3). Recall was 97.7% for *S. aureus* and 88.4% for *E. faecalis*, but only 47.3-51.0% for *E. cloacae*, *E. coli* and *K. pneumoniae* (Figure 8). Training on clinical isolates therefore did not give consistently high performance across the species in these new experiments. This is a single-seed, trap-level result. It cannot be compared directly with Task 2 because the test chips and species differ.

The previous CNN models were evaluated with the labeling and empty-trap flagging from the seven-species study, giving 6,026 Clear medium traps, compared with 5,790 in the main evaluation. The original labeling tool automatically flagged and removed traps with little or no bacterial content, but still allowed partly empty traps, in which bacteria appeared only towards the end of the time-lapse. In the present study, such traps were flagged as empty and removed from the main evaluation, since the aim is to evaluate species identification over time, not trap loading efficiency.

| Model | Test data | Species | Traps | Macro F1 (%) |
|---|---|---|---|---|
| Video ResNet | Clear medium, chips 41-43 | 5 | 6,026 | 66.6 |

Table 3: Final model after 150 epochs of training on clinical chips 28-40. Macro F1 averages the five species present in the test set. Predictions into all six output classes count when calculating the scores.

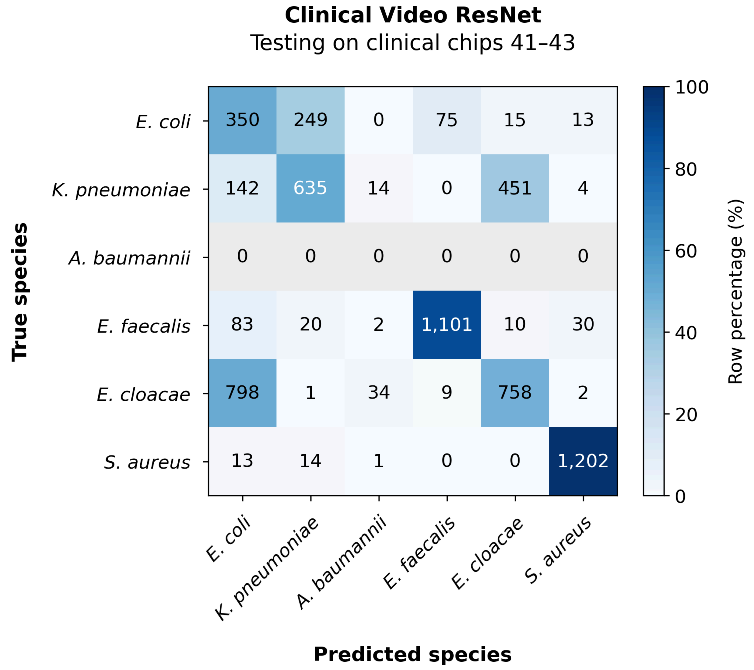


Figure 8: Video ResNet trained on clinical chips 28-40 and tested on chips 41-43. Cells show trap counts and colours show row percentages. All six outputs are retained. The grey row has no true examples.
