## Supplementary material for "Label-free identification of bacterial species using adapted vision transformers": S2 Document

### S2 Document: Can autofluorescence aid classification?

We tested whether autofluorescence could improve bacterial species classification. After acquiring the final frame of each phase-contrast sequence, we illuminated the cells with laser light. Naturally occurring fluorophores absorb excitation light and emit fluorescence as they return to a lower-energy state. This emitted light is recorded by the imaging detector.

We trained a simple ResNet18 from random initialization using either the final phase-contrast image, the autofluorescence image, or both. Each crop was 52 pixels wide and 114 pixels high. Matching crops were placed side by side in one input channel. For the single-modality controls, the same crop was duplicated, keeping the input size identical (104 wide × 114 high).

Mean macro-F1 across 31 training seeds was **0.526 ± 0.017** for autofluorescence, **0.626 ± 0.008** for phase contrast, and **0.659 ± 0.02** for the combination (mean ± sample SD).

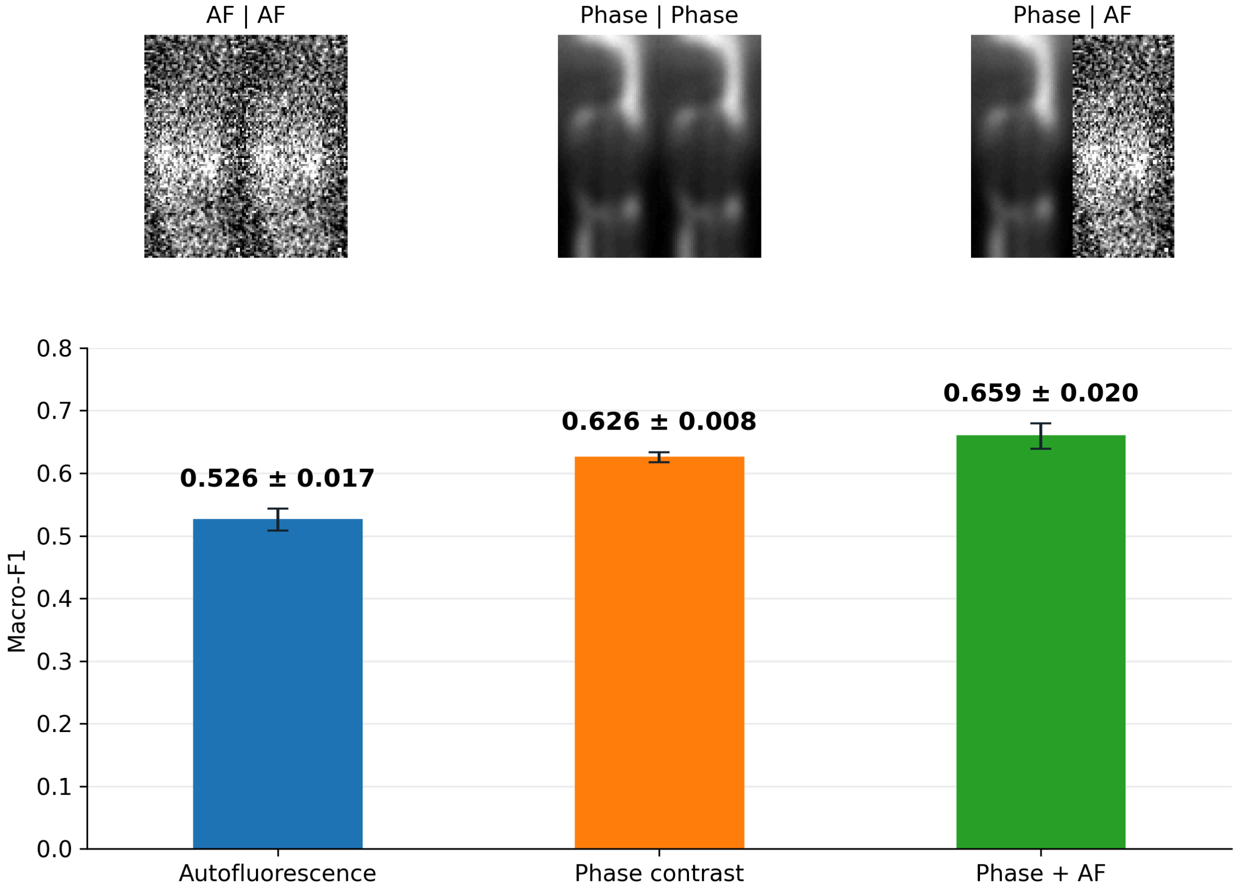


Figure 1: Bars show mean macro-F1 across 31 training seeds; error bars show sample SD. Example inputs show the same held-out *E. faecalis* trap at rows 0–113, without augmentation. Phase is min–max normalized; AF uses P5/P95 clipping for display only (training and evaluation used P1/P99). The trap was selected visually to illustrate the input arrangement.

Training used 32,244 traps from chips 28–40, and testing used 5,790 traps from chips 41–43, covering five species: *Enterobacter cloacae*, *Escherichia coli*, *Enterococcus faecalis*, *Klebsiella pneumoniae* and *Staphylococcus aureus*. Each input condition was trained with the same 31 random seeds. Full preprocessing, training and evaluation details are given under "Training recipe".

### Where do the models make errors?

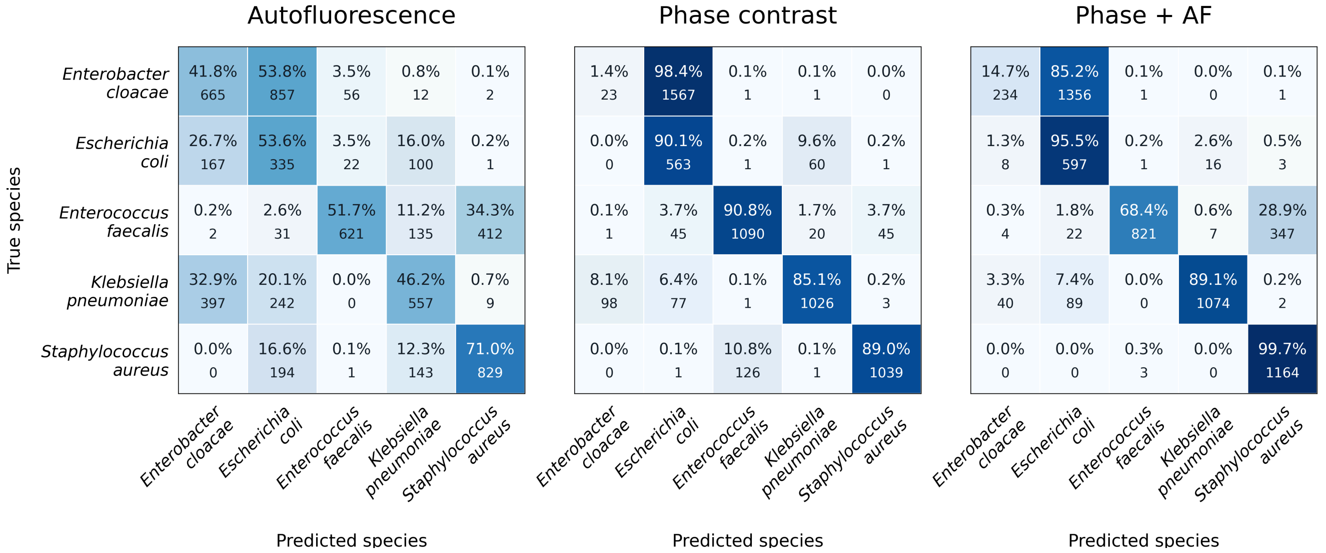


Figure 2: Confusion matrices for the run with median macro-F1 within each input condition. The larger number is the percentage of the true-species row; the smaller number below is the absolute number of traps. All panels use the same 0–100% color scale. Rows are true species and columns are predictions. The summary on the previous page includes all 31 seeds, rather than only these representative runs.

Phase contrast outperformed autofluorescence alone. Adding autofluorescence increased mean macro-F1 by 3.4 percentage points, with an improvement in 30 of 31 paired seeds. Thus, autofluorescence provided additional information under this experimental setup.

The improvement was not uniform across species. Mean recall for *E. cloacae* remained low: 2% with phase contrast and 7.1% with the combination. Larger recall gains were seen for *K. pneumoniae* (77.8% to 88.6%) and *S. aureus* (91.9% to 99.7%).

These results describe the tested chips. Seed variation measures training variability, not uncertainty across new chips; the recipe was developed through previous exploratory comparisons using these same test chips.

#### Training recipe

We trained ResNet18 from random initialization, with all weights trainable, one input channel and five output classes. Training used 32,244 traps from chips 28–40; testing used 5,790 traps from chips 41–43. Acinetobacter was excluded, removing training chip 31. The three inputs were AF + AF, phase + phase and phase + AF. Phase images were the last stored frames. Each modality supplied a 114-pixel-high × 52-pixel-wide crop, giving a 114 × 104 single-channel image after side-by-side concatenation, without resizing.

Growth extents were read from `data/growth_extent.csv` and matched by trap path. The same extent was used for both modalities, capped at 1,300 pixels. During training, one crop was sampled per trap per epoch. Its top coordinate was drawn uniformly from zero to max(0, min(growth extent, 1,300) − 114), inclusive. Phase and AF used the same coordinate. When growth was shorter than 114 pixels, the top 114 pixels of the original image were used, including any area below the measured growth extent.

Normalization was performed separately for each crop before augmentation. AF was clipped at its 1st and 99th percentiles and scaled to 0–1; phase used min–max scaling to 0–1. Constant crops became zeros. Both modalities received the same random affine transformation with probability 0.5: shifts up to ±3%, scale changes up to ±10% and rotations up to ±1°, with bilinear interpolation and reflected borders. No flips or brightness/contrast changes were applied. The images were joined after augmentation; single-modality controls duplicated the same processed crop.

Each condition was trained for five epochs with batch size 32 and seeds 8103–8133. AdamW used weight decay 0.01 and a cosine learning rate schedule from 0.001 at epoch 1 to 0.00001 at epoch 5, without warmup. Training used class-weighted cross-entropy: each example of class c had weight N / (5 × N_c), where N is the total number of training traps and N_c is the number in that class. Weighted losses were averaged over the batch. The final epoch was used for evaluation, without early stopping or test-based checkpoint selection.

At evaluation, 114-pixel-high tiles covered the growth extent with a stride of 57 pixels (50% overlap). A final boundary-aligned tile was included when needed, with duplicate positions removed. Short extents yielded one top tile. Normalization was unchanged and no augmentation was applied. Raw logits were averaged across all tiles in each trap; the class with the highest mean logit was the prediction. Softmax was applied to the mean logits to obtain probabilities for one-vs-rest ROC-AUC.

Macro-F1 was calculated across all five species and summarized as the mean and sample standard deviation over 31 training seeds. Confusion matrices show the run with median macro-F1 within each condition. One-vs-rest ROC-AUC was calculated separately for each species.
