## Supplementary material for "Label-free identification of bacterial species using adapted vision transformers": S3 Document

### S3 Document: Blood dataset, results breakdown by isolate and chip

Confusion matrices for the selected median run, the same run shown in the Blood and Clear medium confusion matrices in the main paper, evaluated using 55 frames (54 minutes) of the original images, without simulated NA degradation. Each panel identifies the clinical isolate (EL number) and chip.

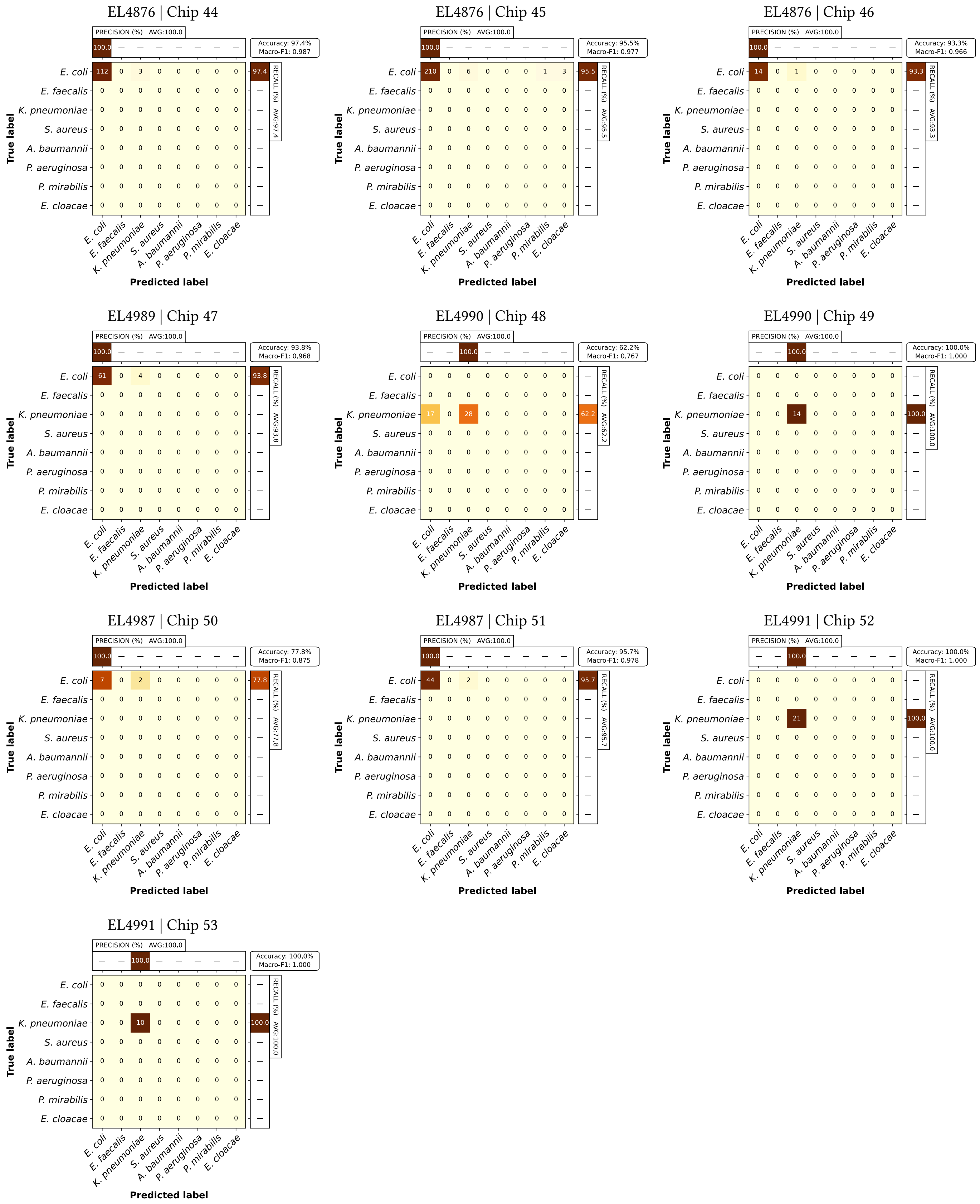


**Supporting figure.** Blood test results for chips 44–53 (560 traps). Matrix entries are trap counts; colours indicate percentages within each true-label row. The upper and right margins show precision and recall, respectively, in percent. Accuracy and macro-F1 are reported for each chip. Repeated EL numbers represent separate chips from the same isolate.
