## Supplementary figures and images for "Label-free identification of bacterial species using adapted vision transformers"

### S1 Fig

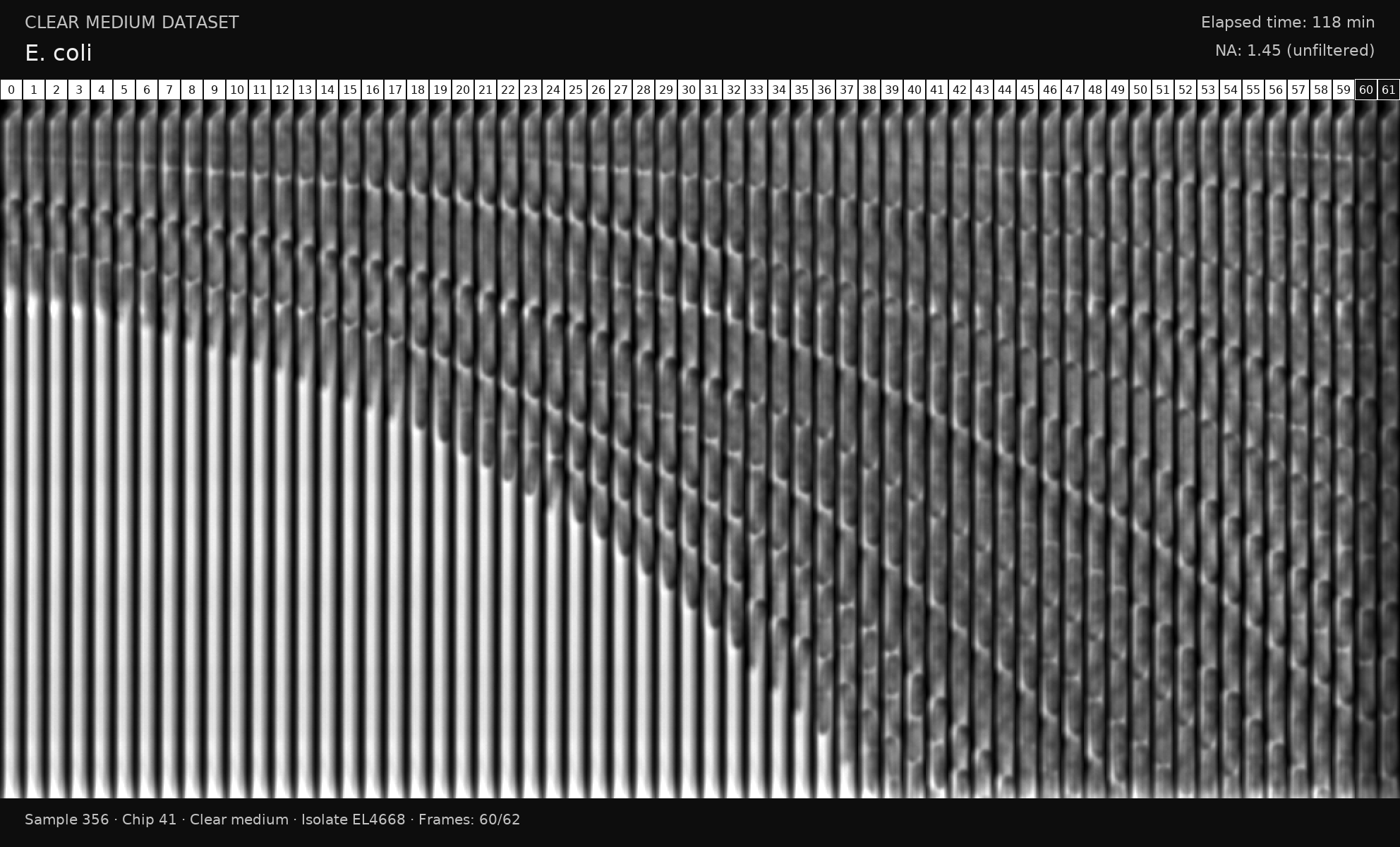

### S2 Fig

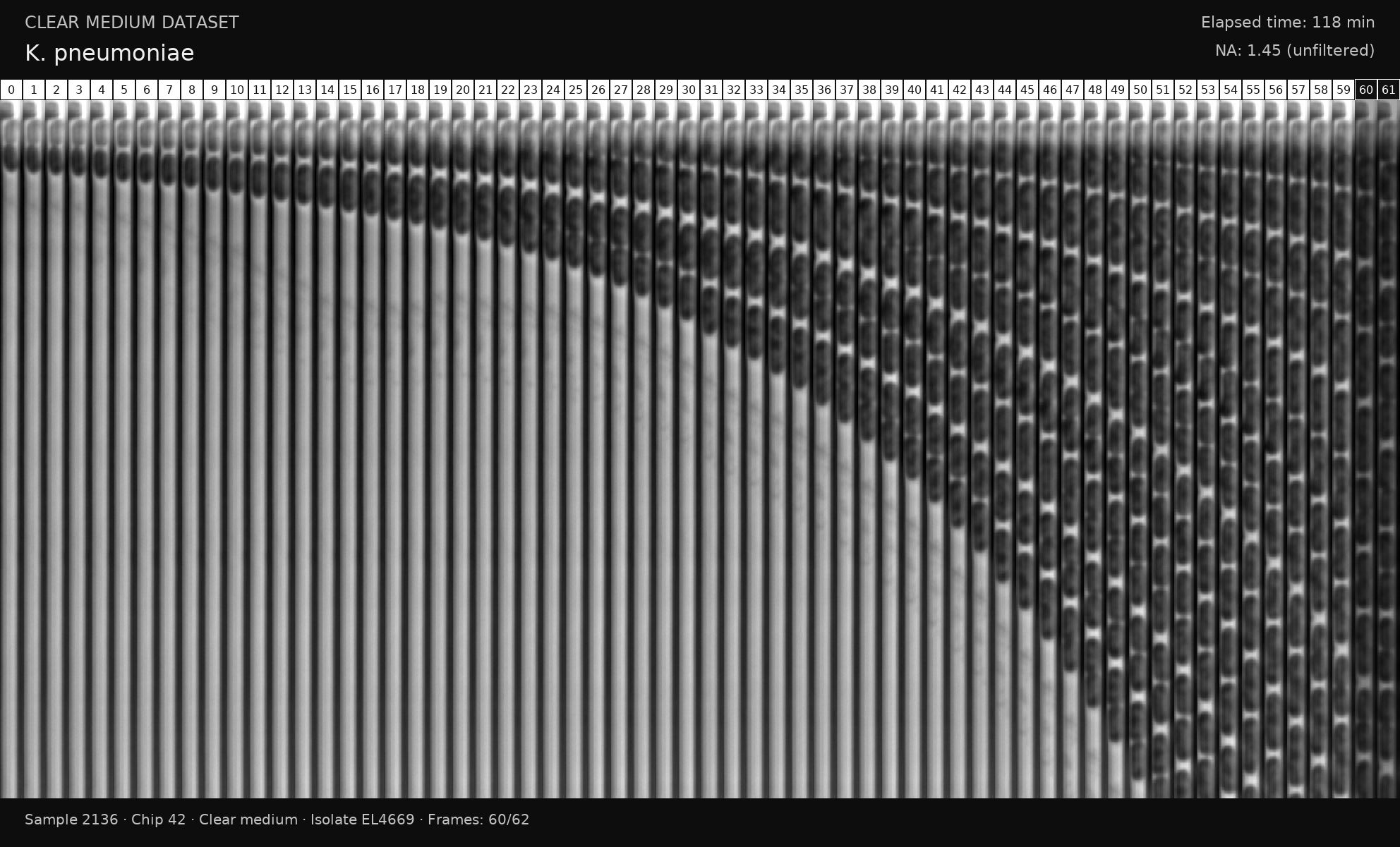

### S3 Fig

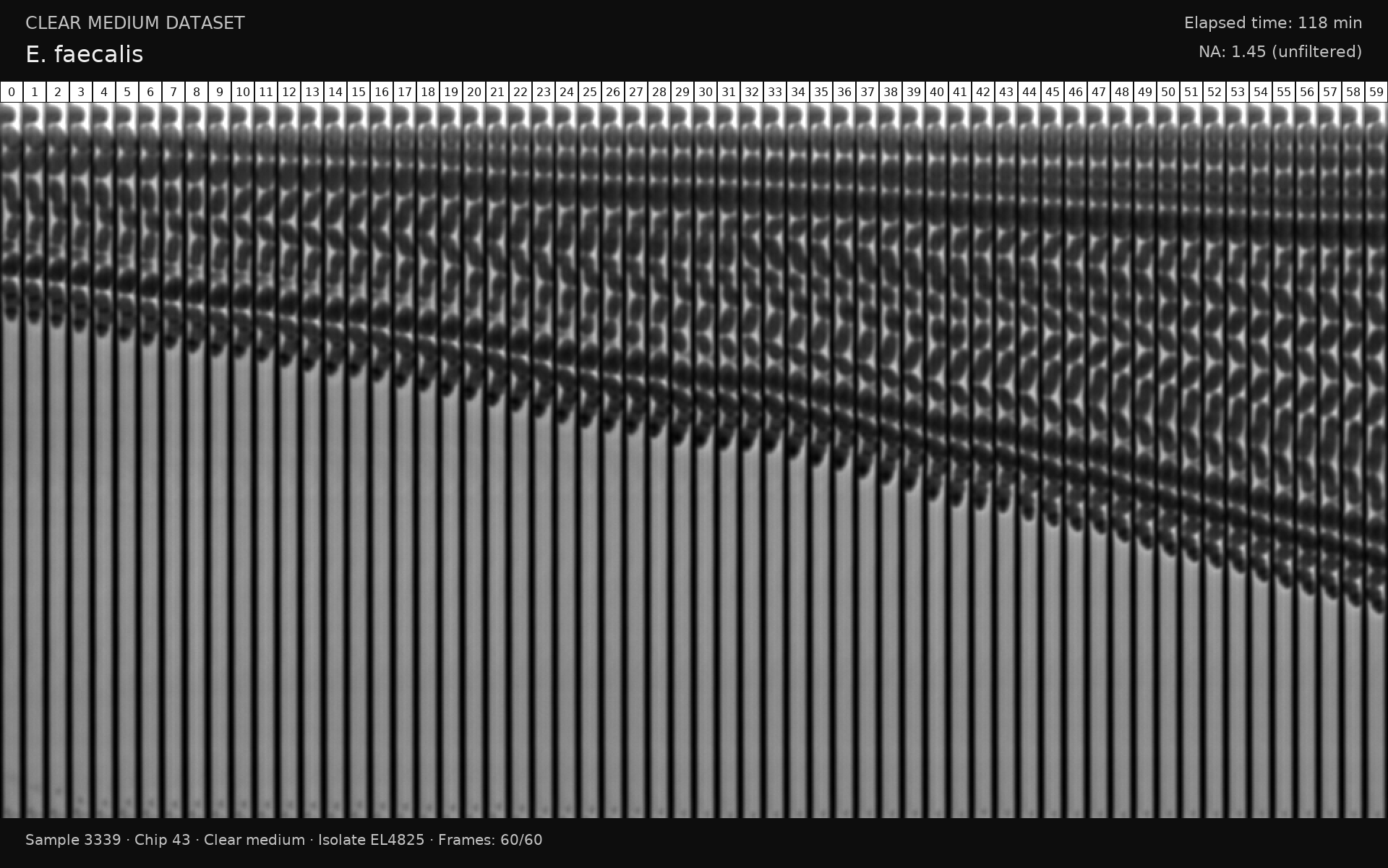

### S4 Fig

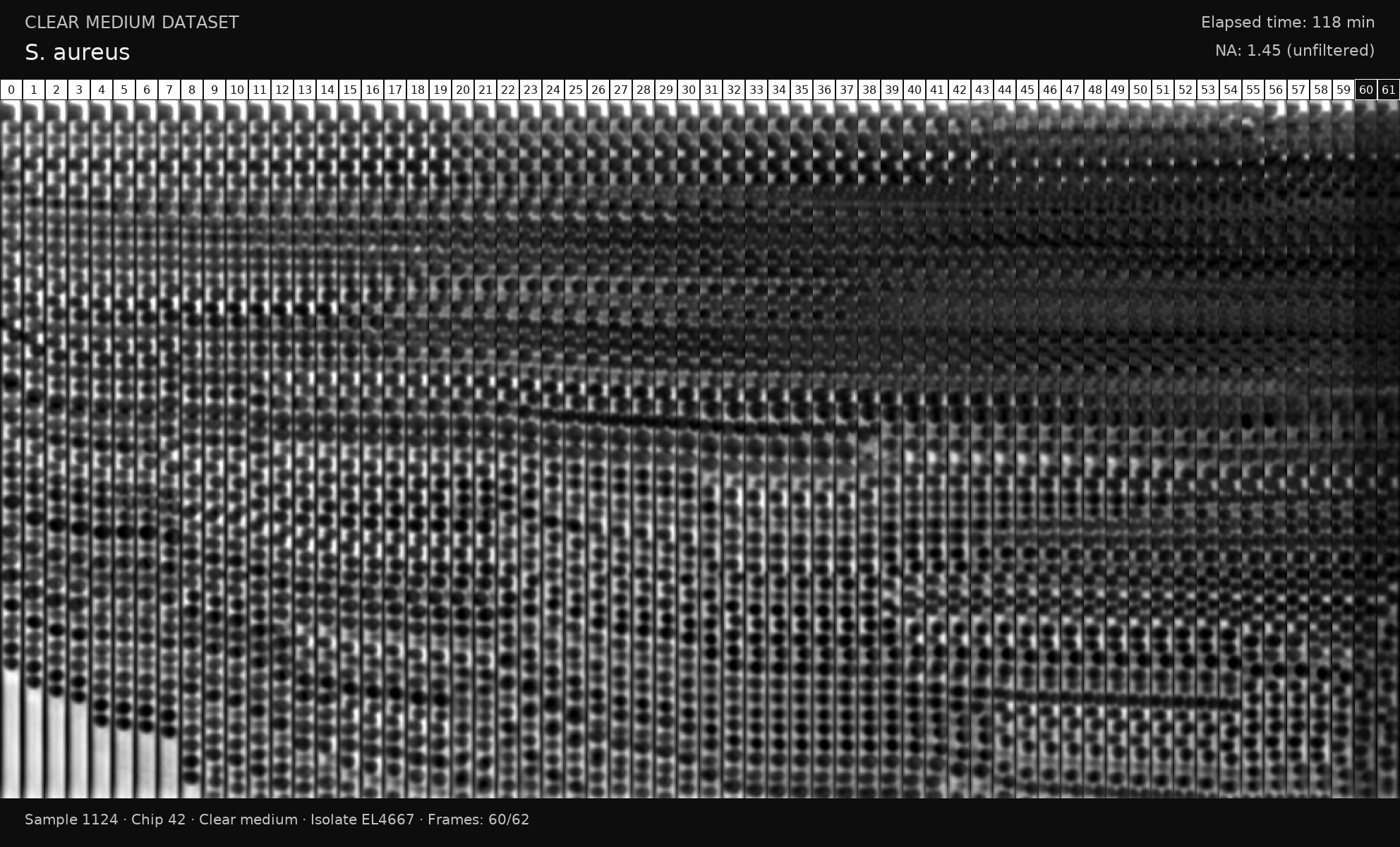

### S5 Fig

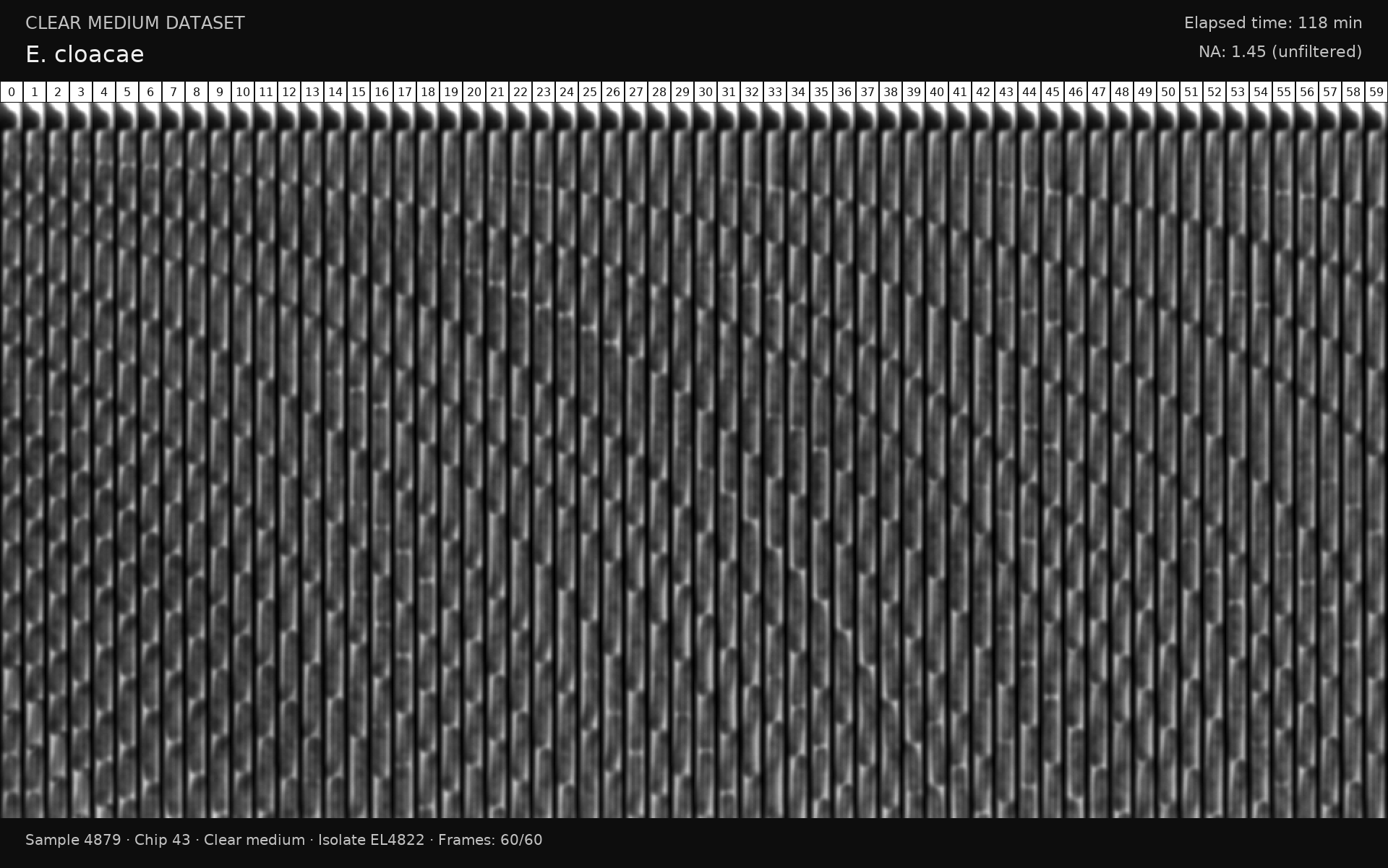

### S6 Fig

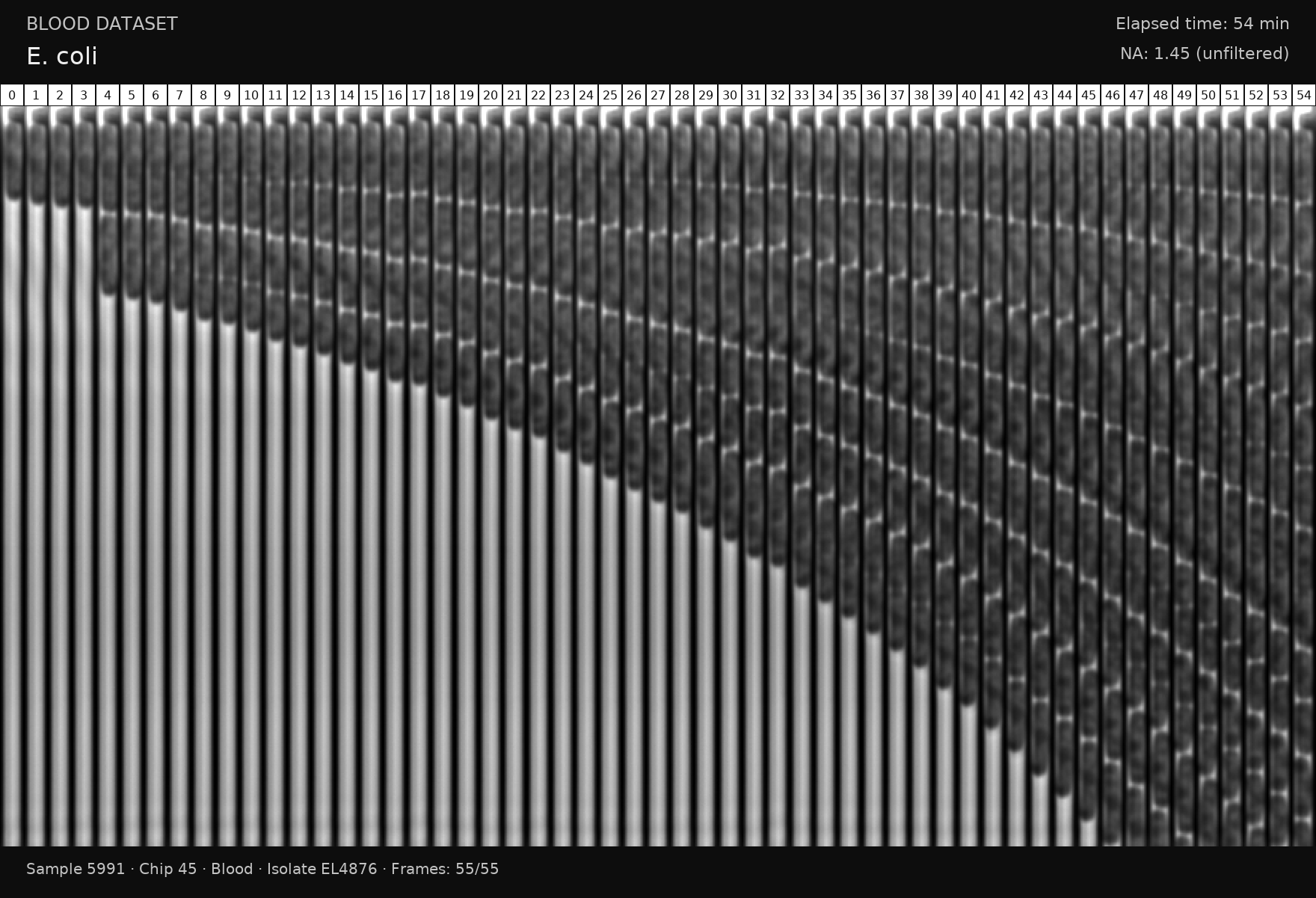

### S7 Fig

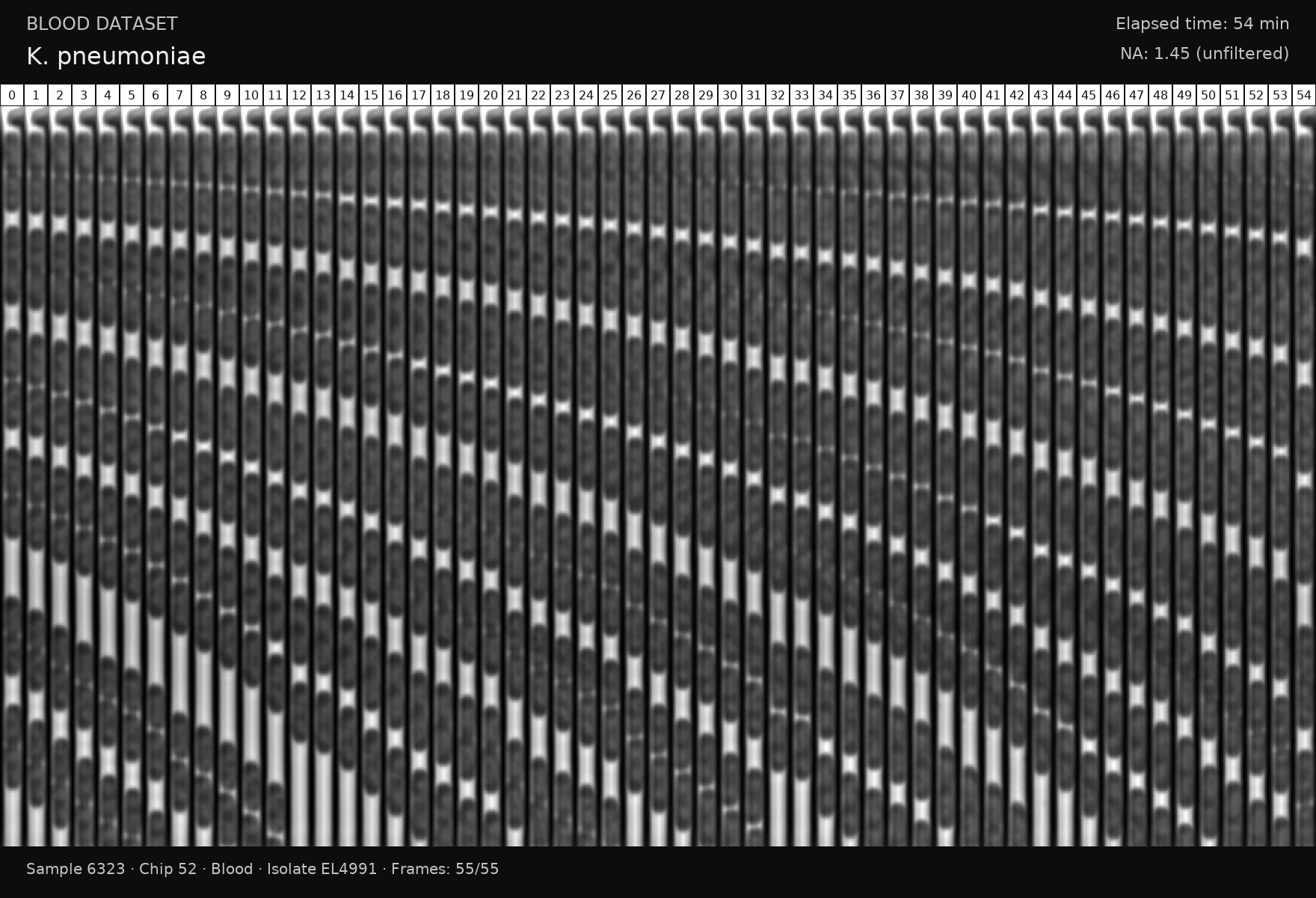

### S8 Fig

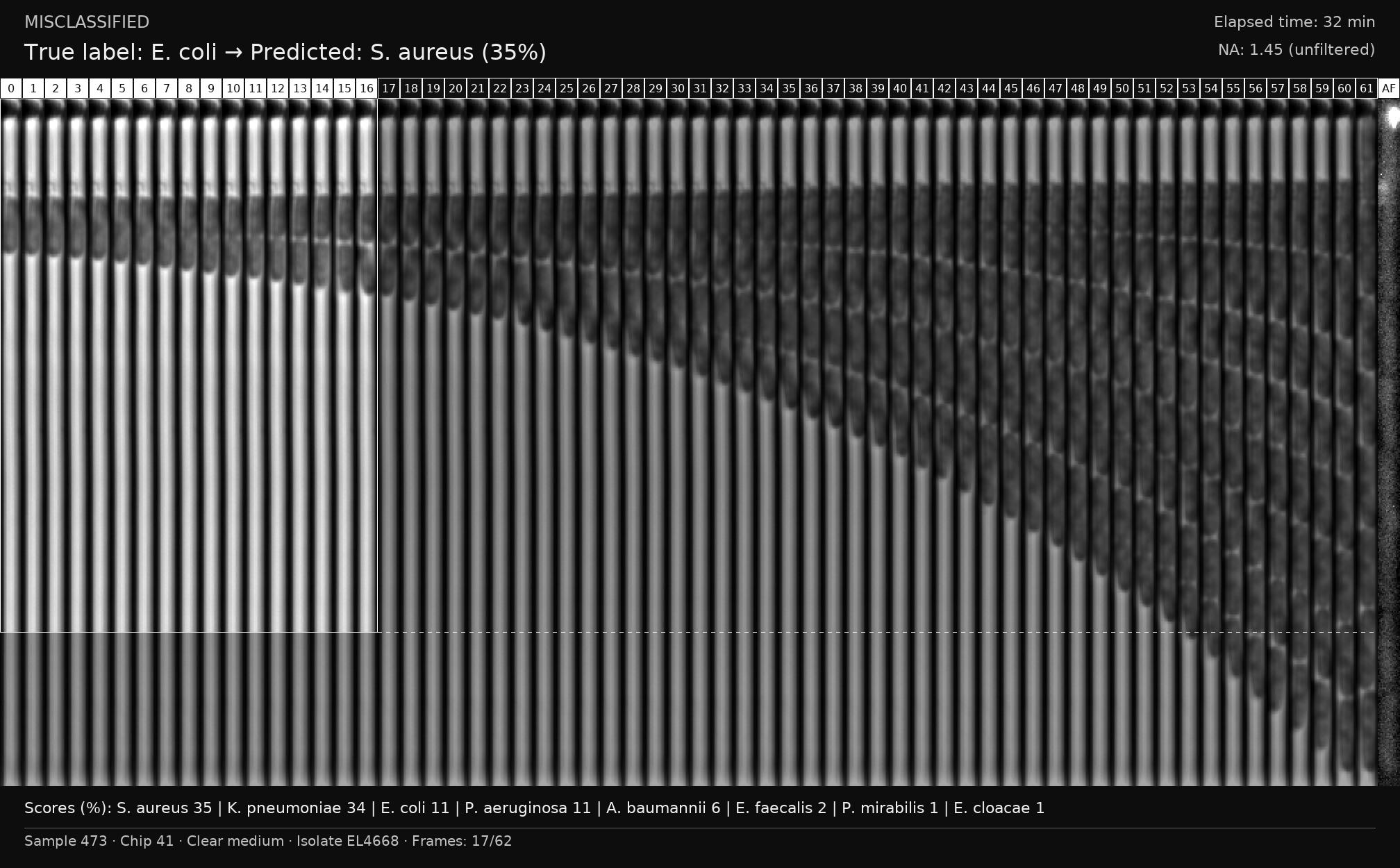

### S9 Fig

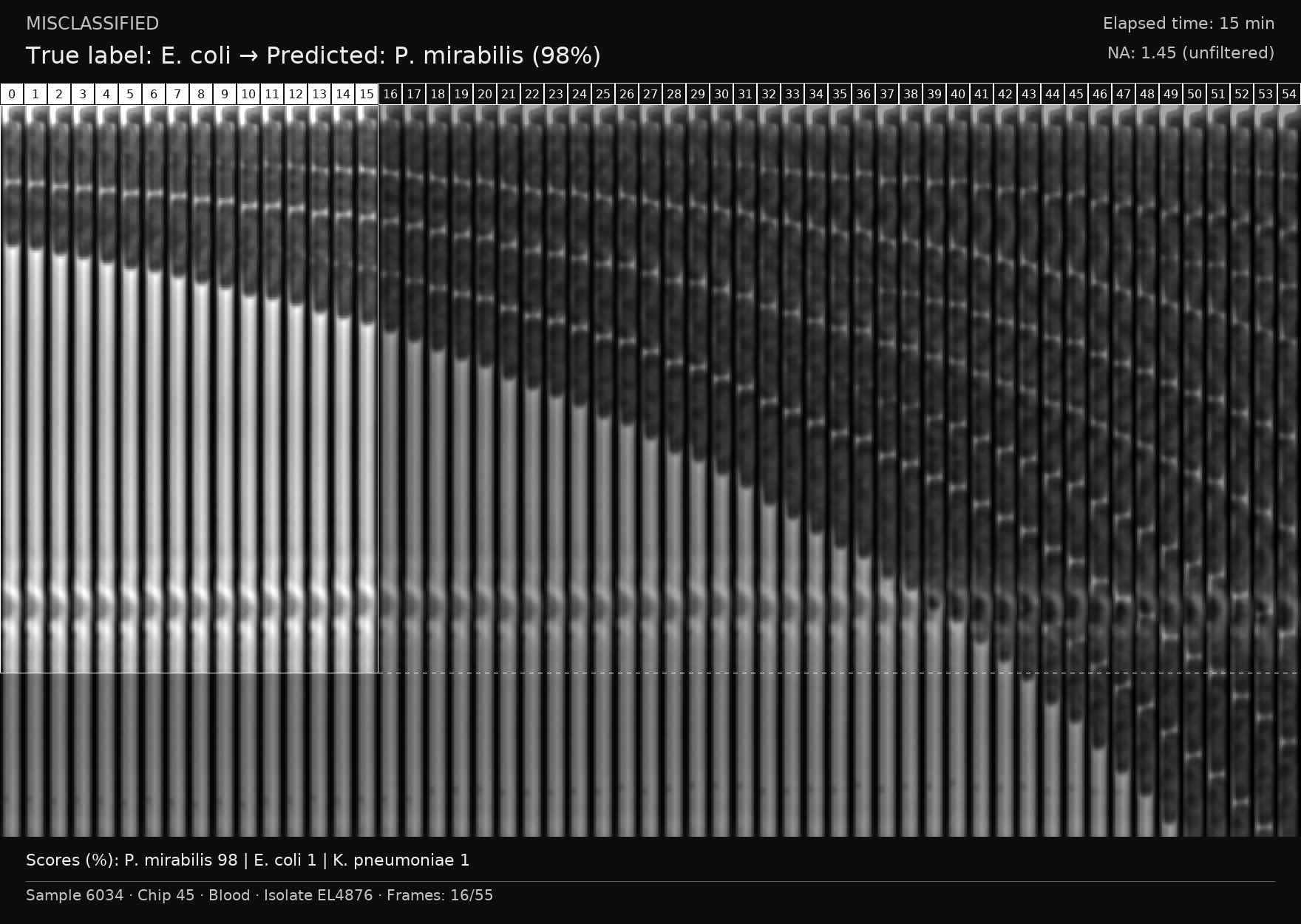

### S10 Fig

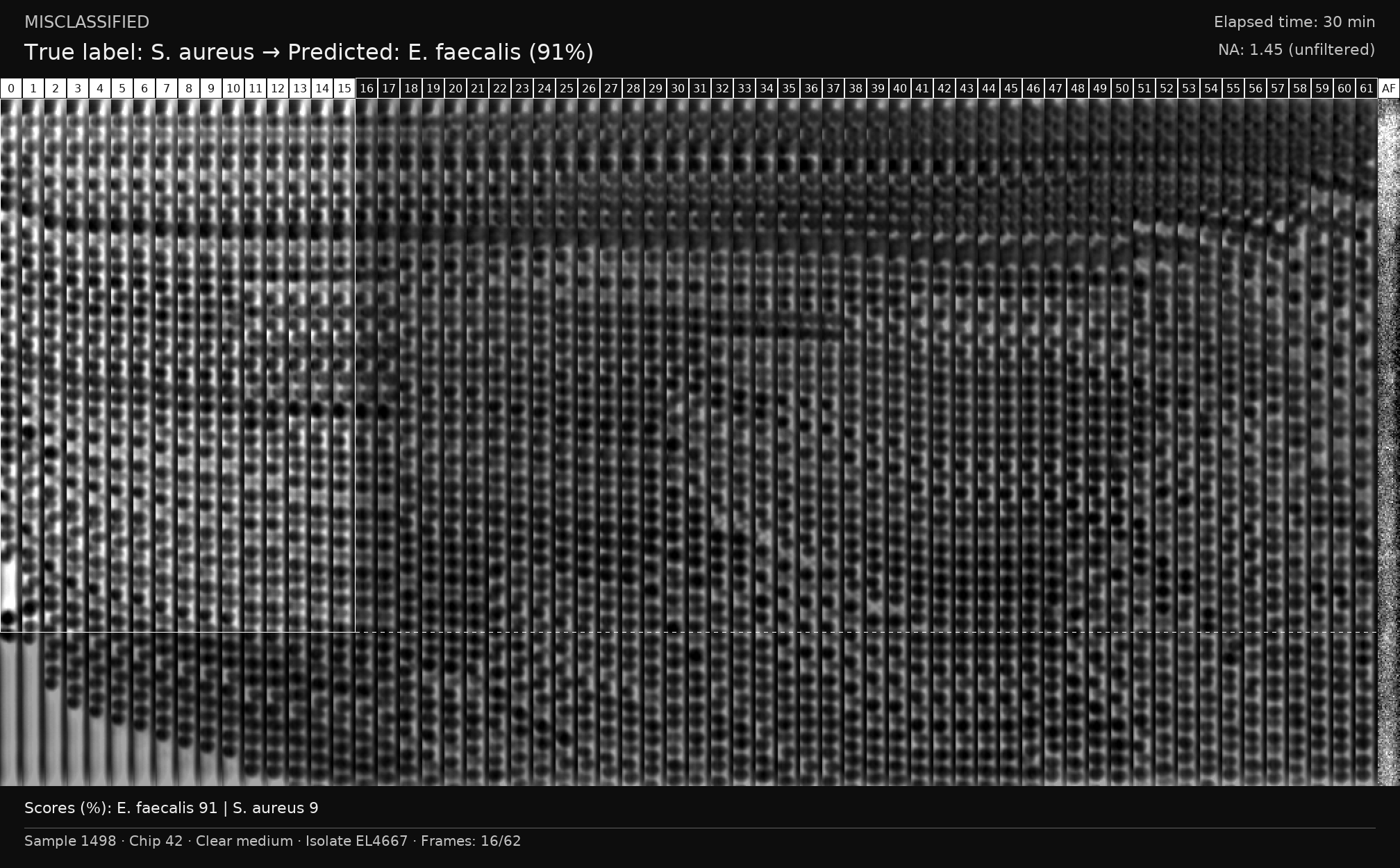

### S11 Fig

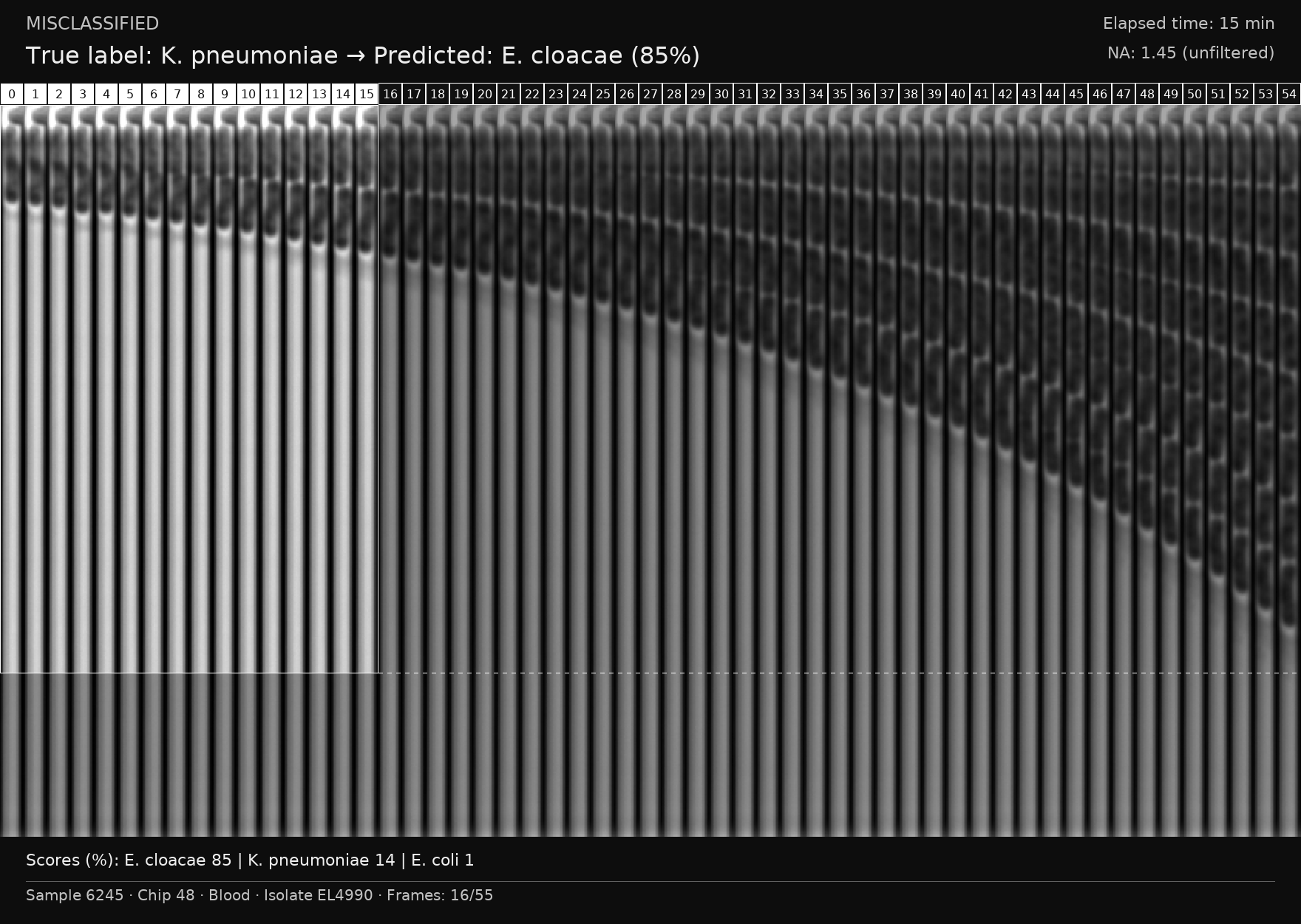

### S12 Fig

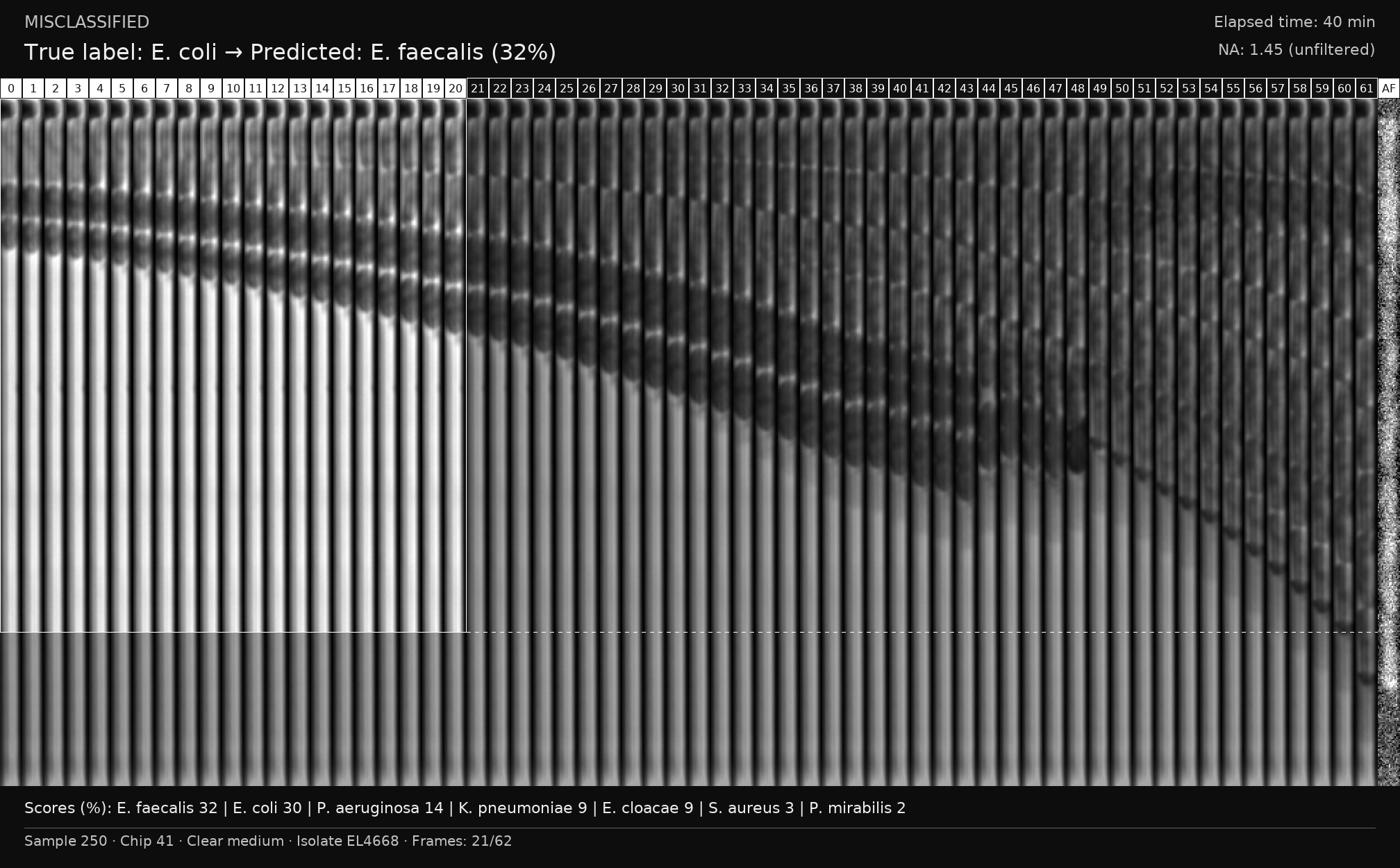

### S13 Fig

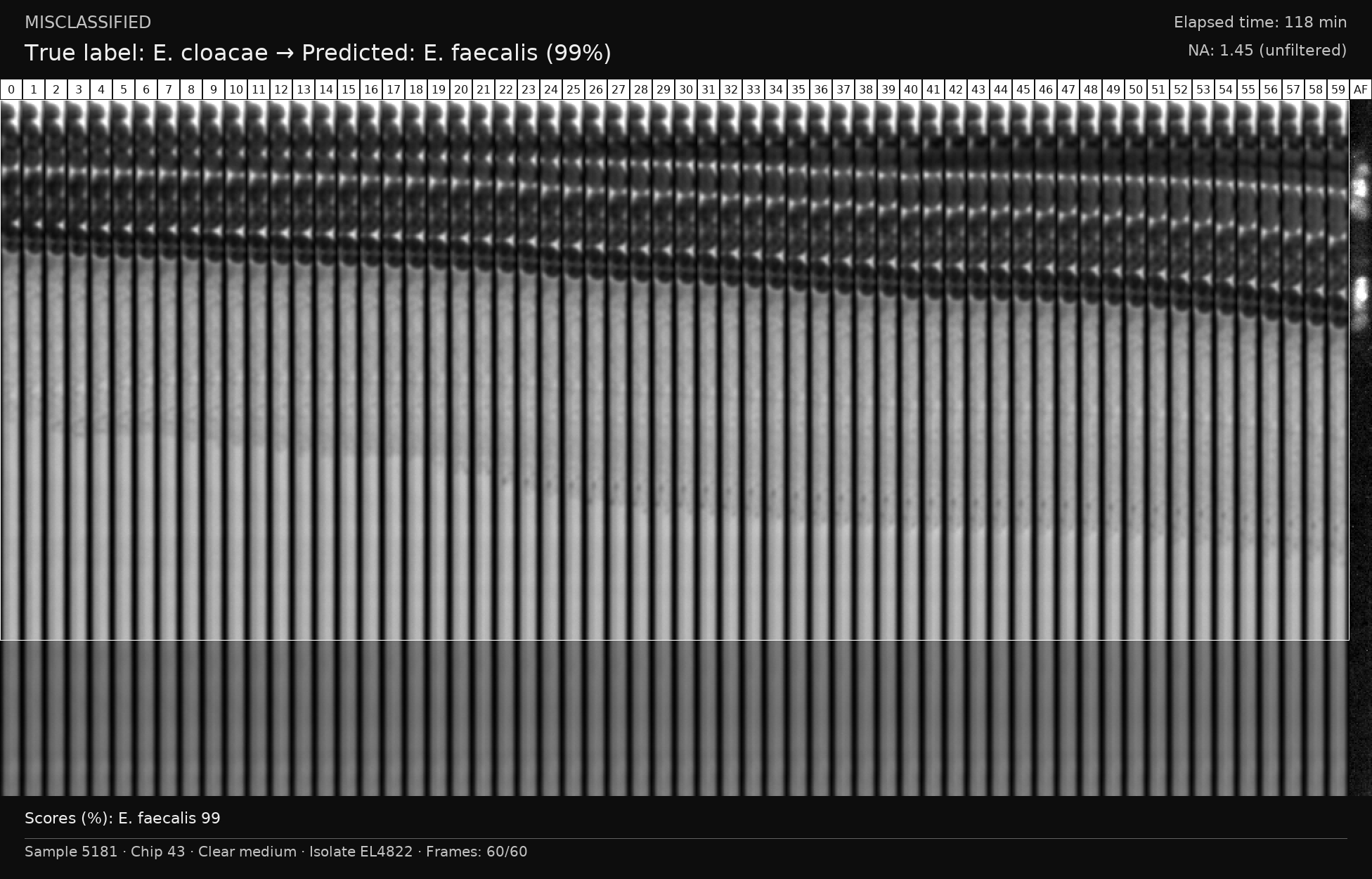

### S14 Fig

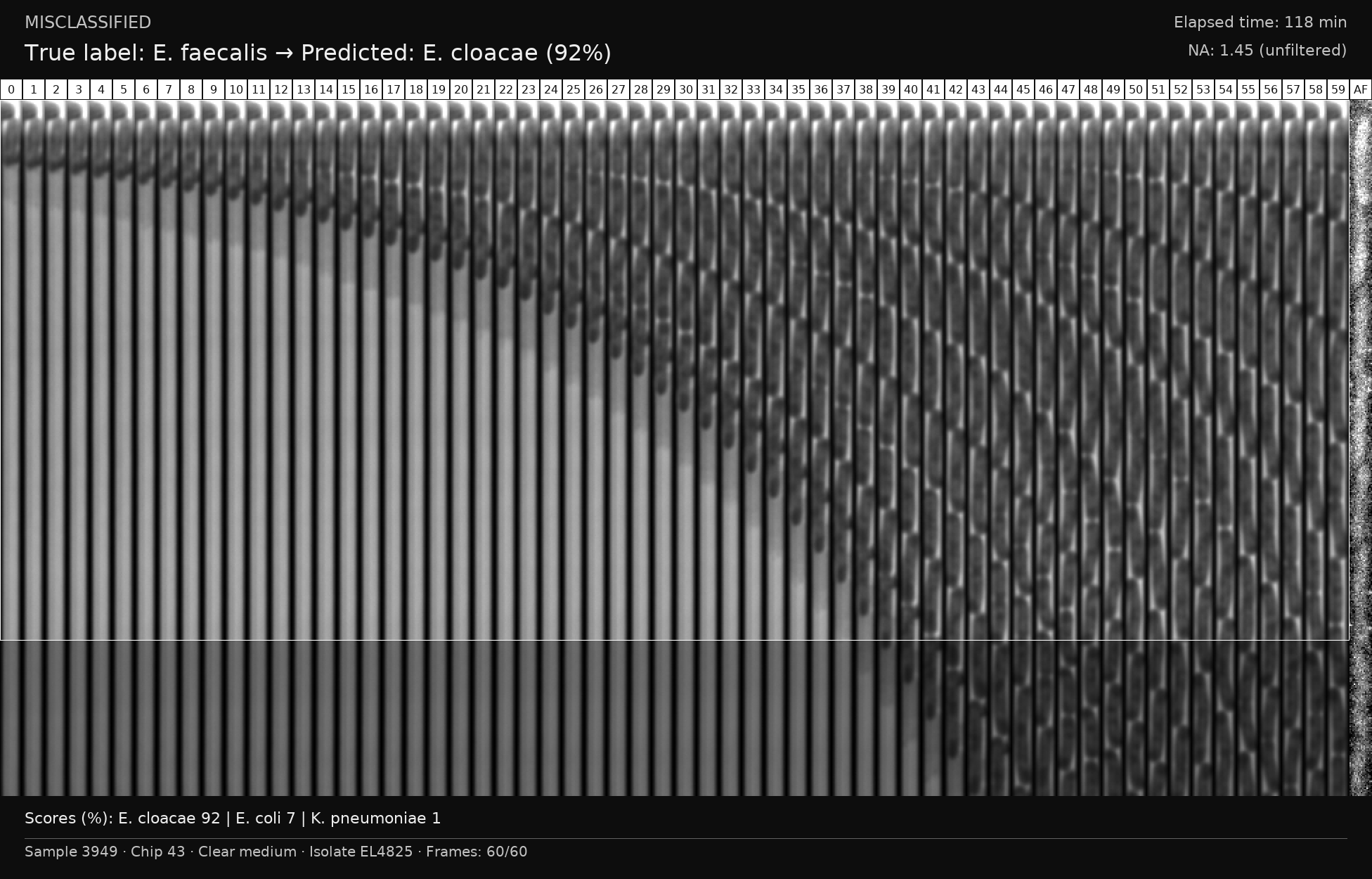

### S15 Fig

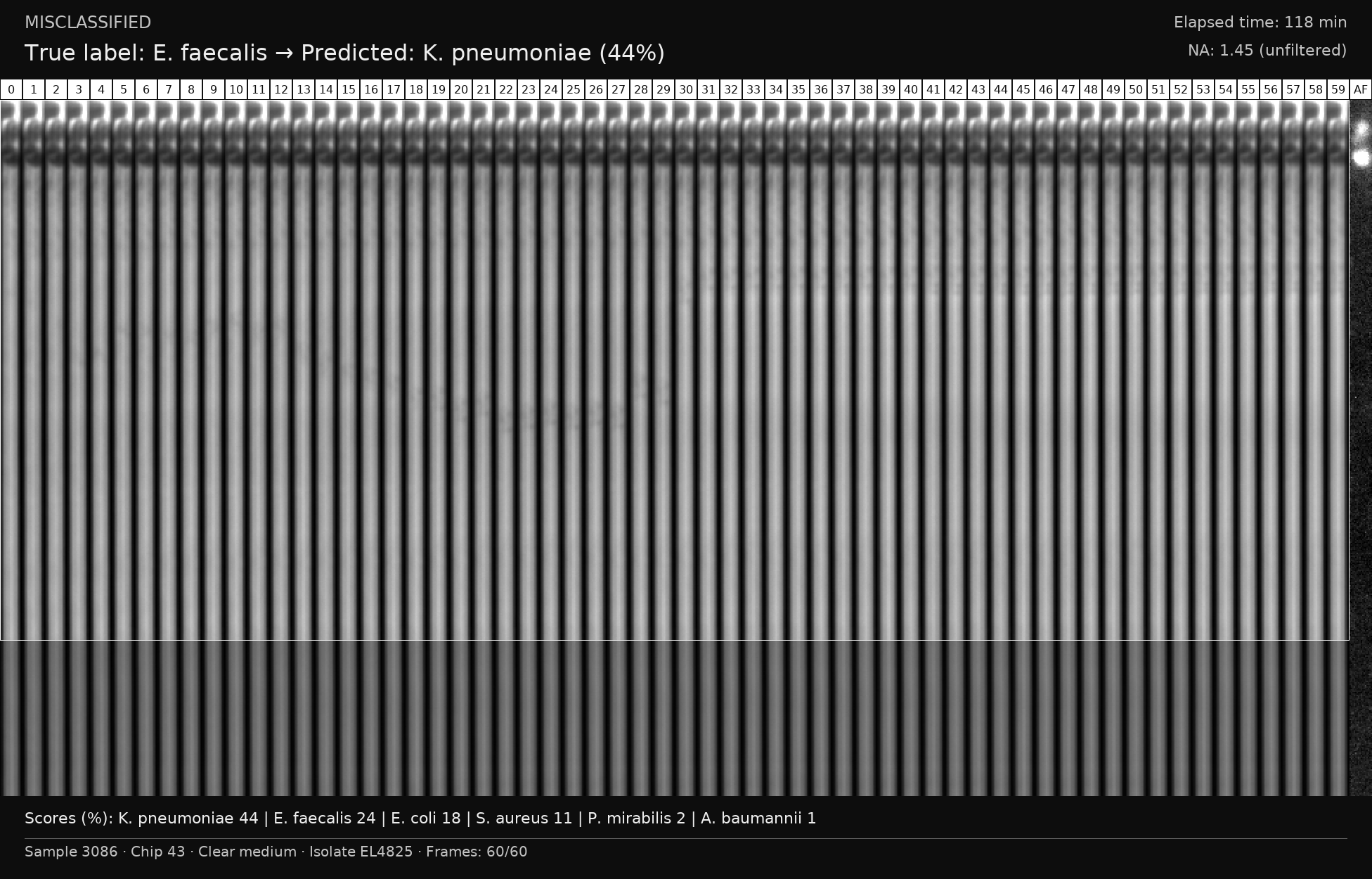

### S16 Fig

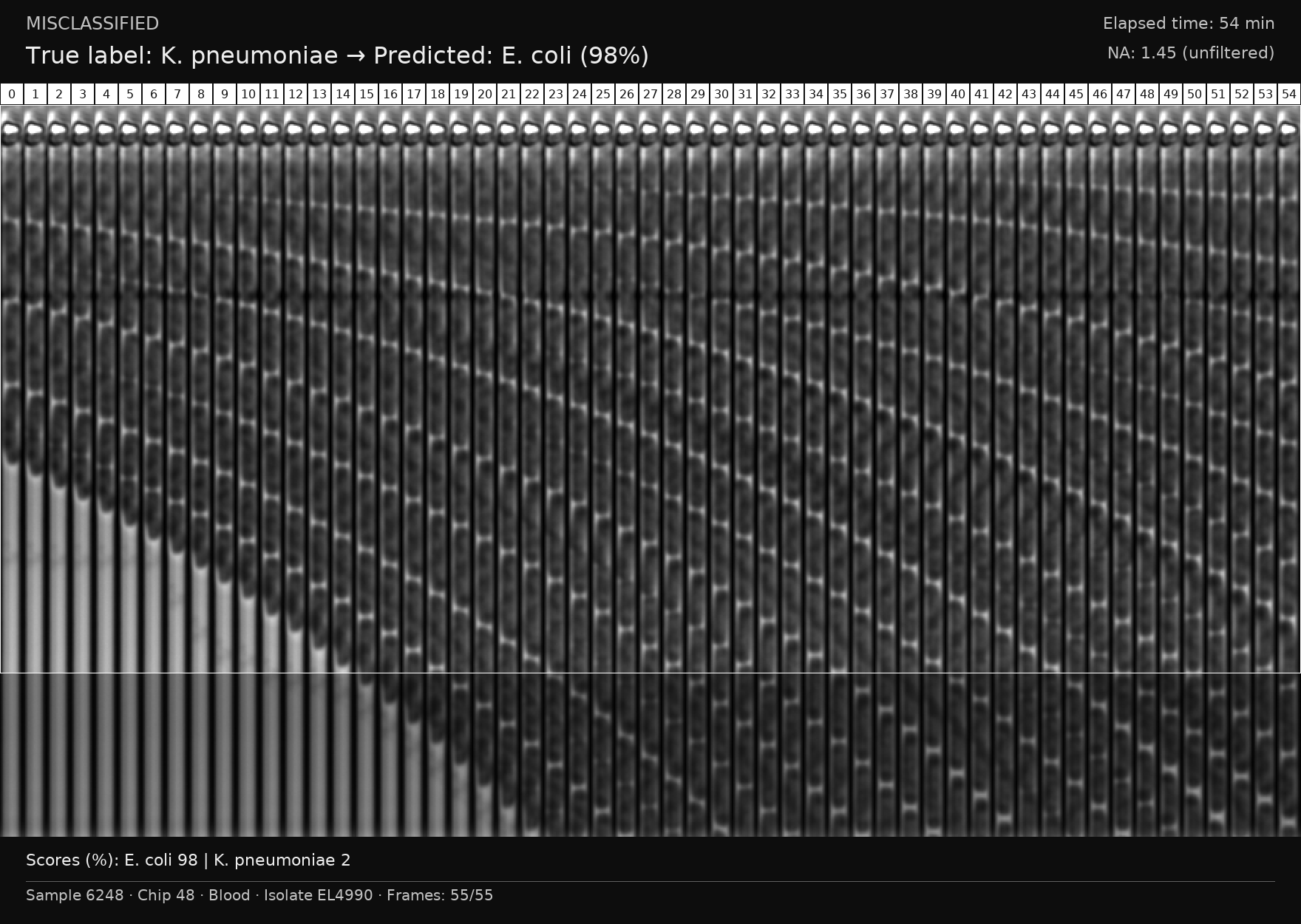

### S17 Fig

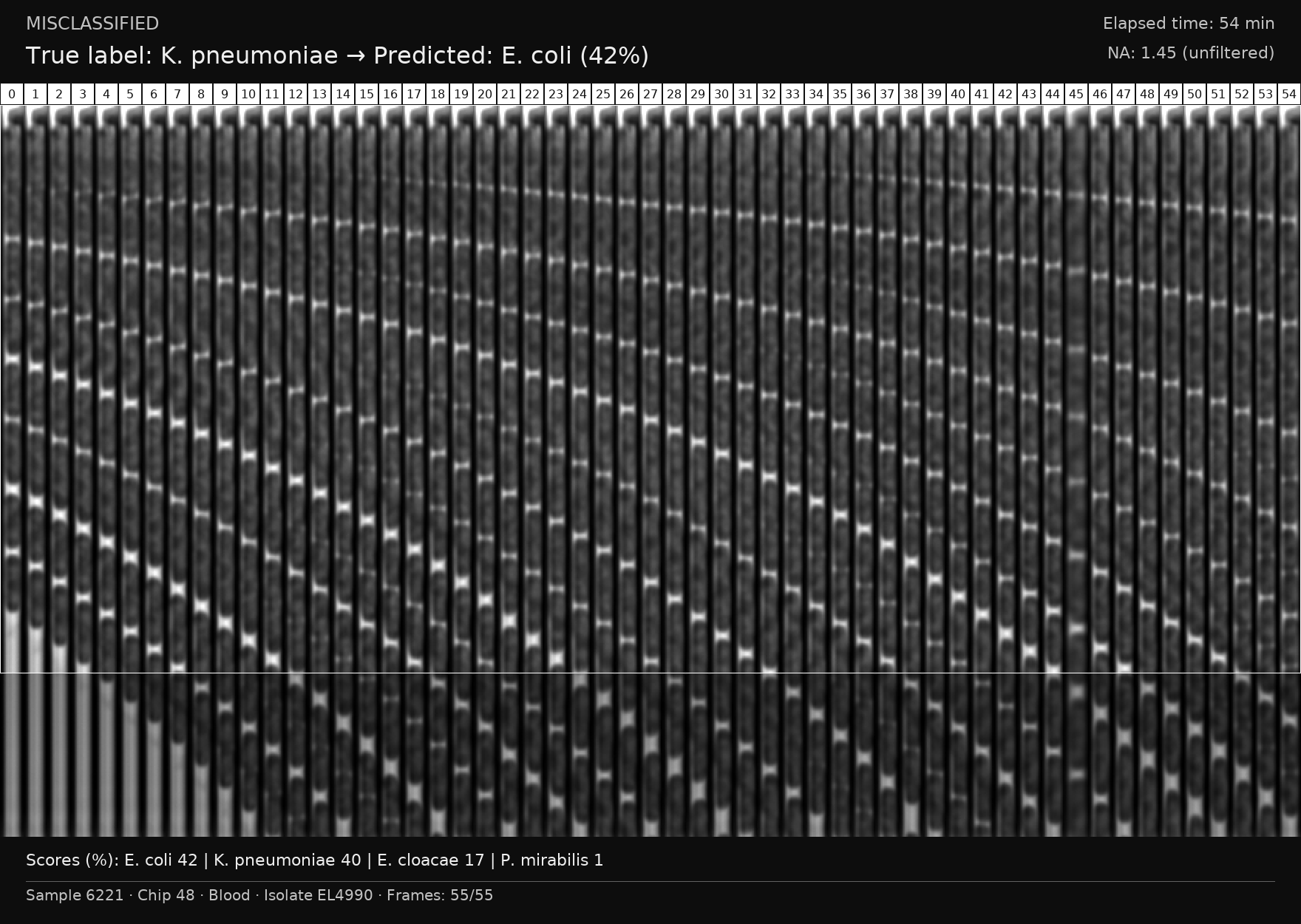

### S18 Fig

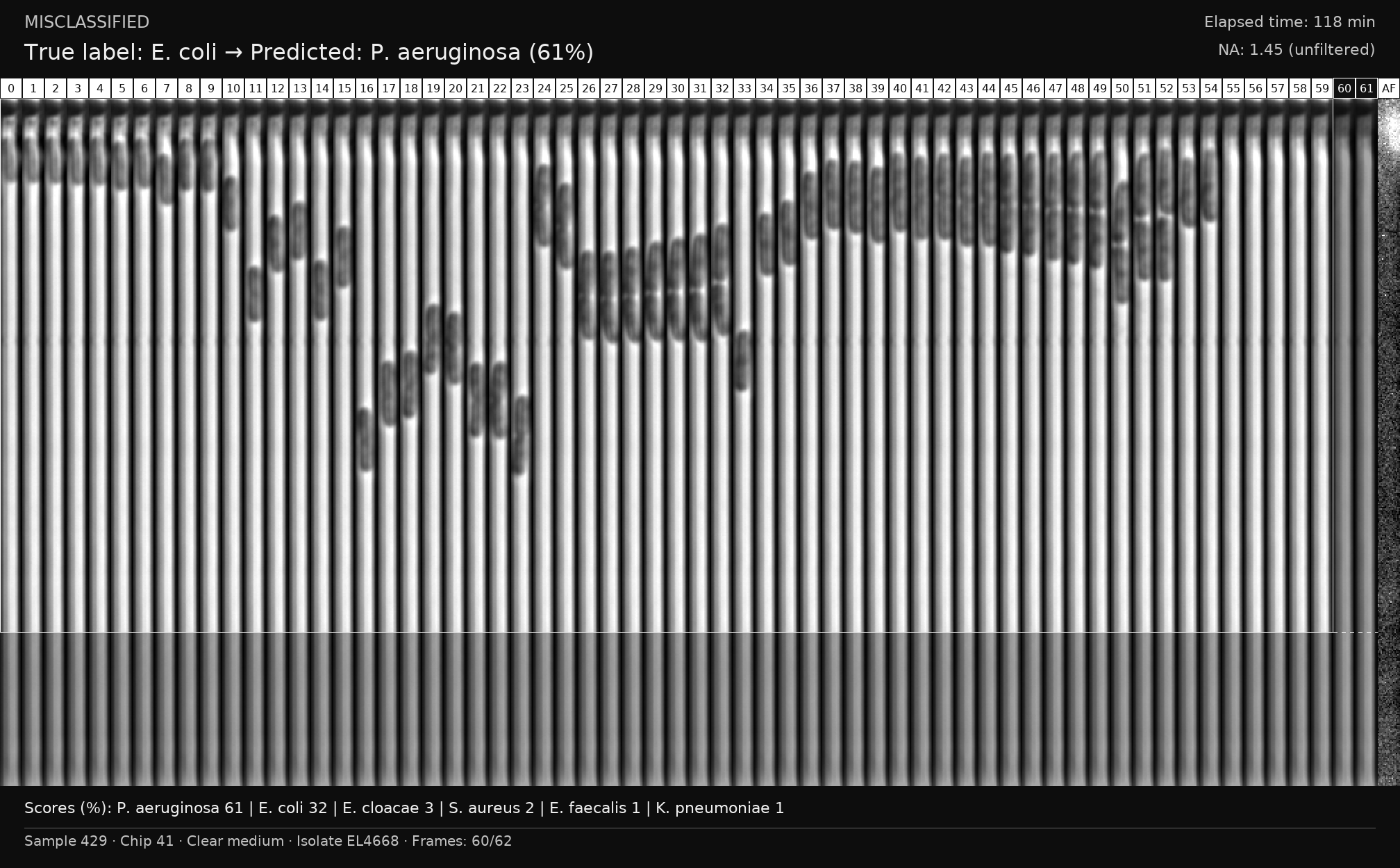

### S19 Fig

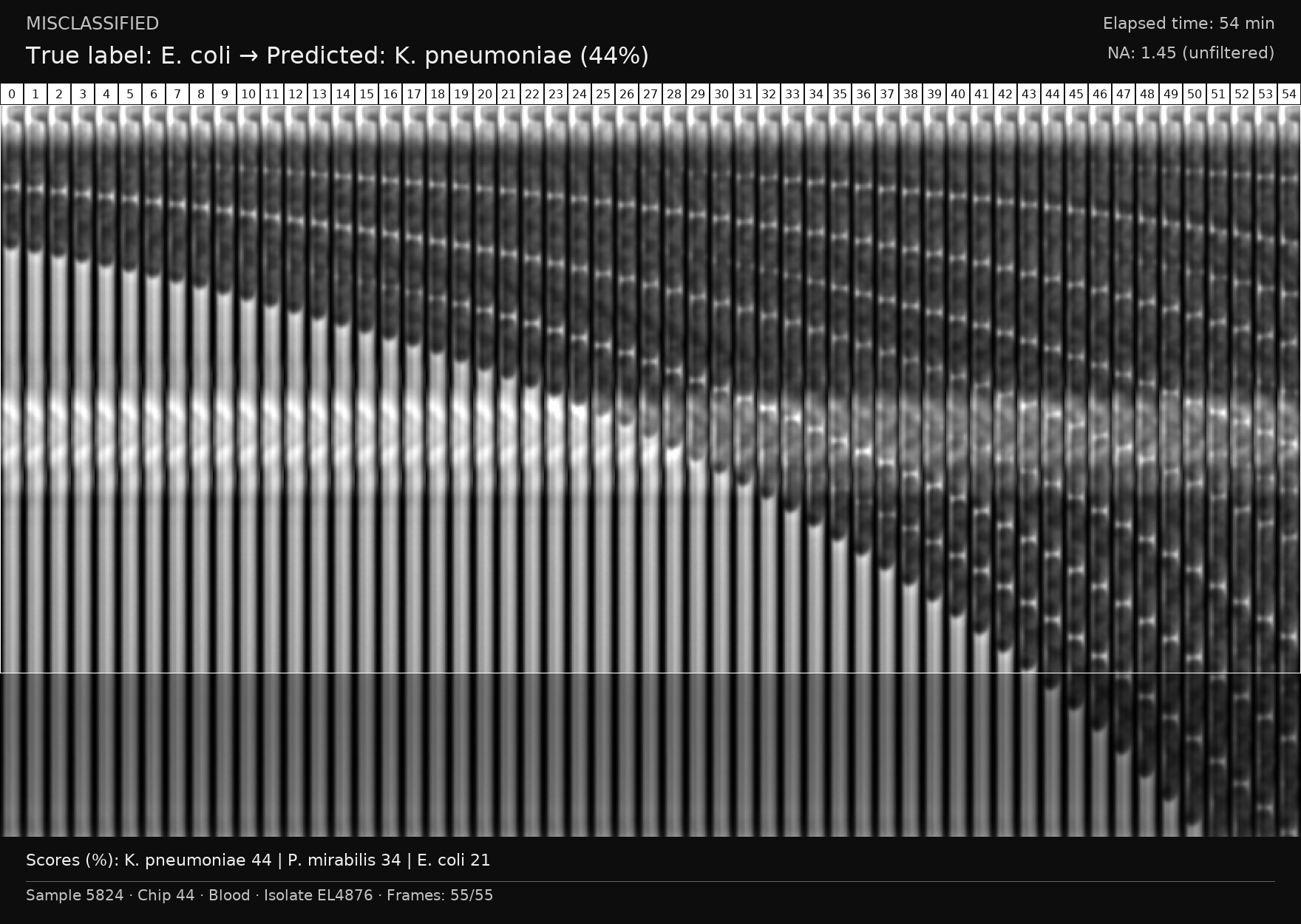

### S20 Fig

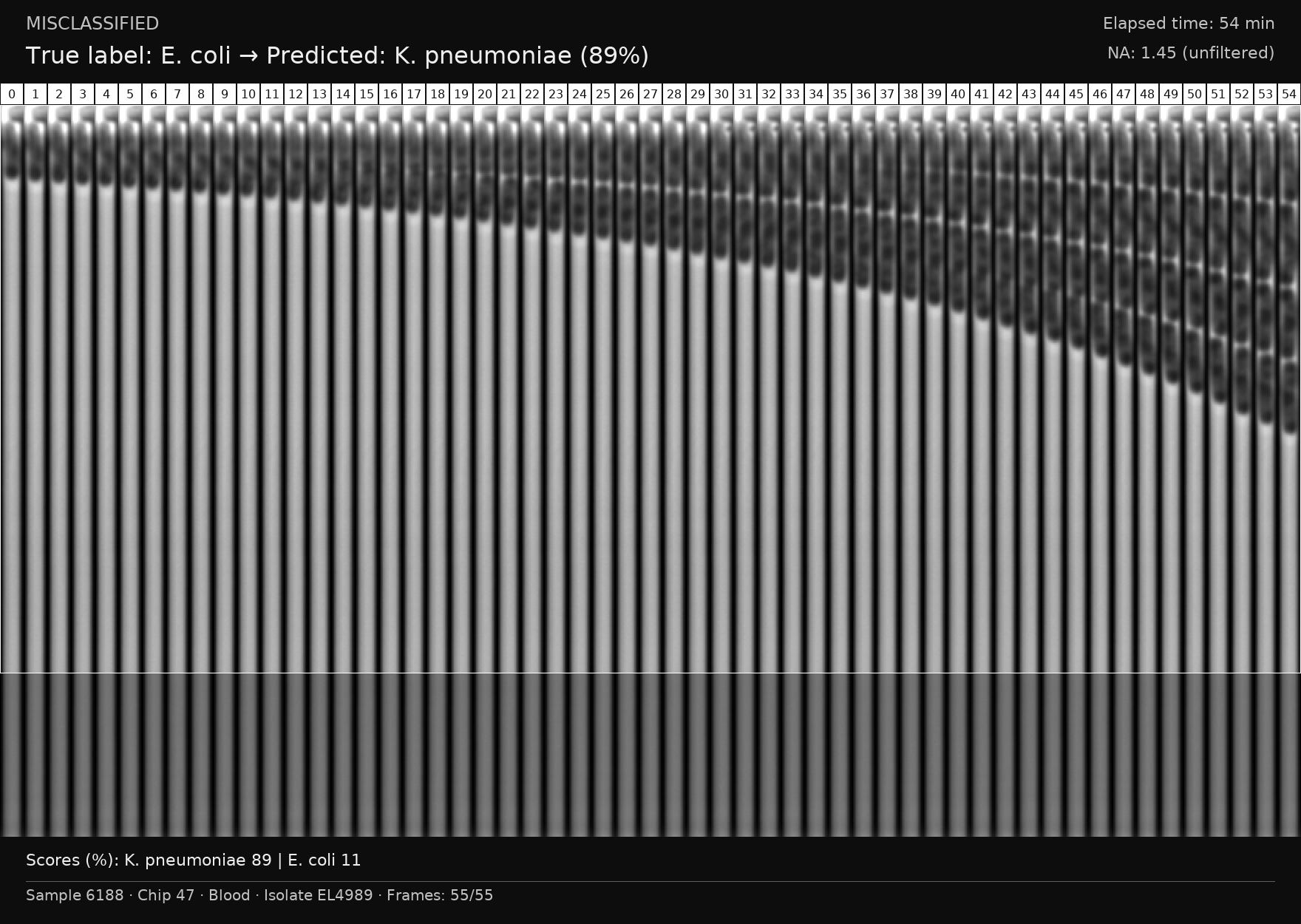

### S21 Fig

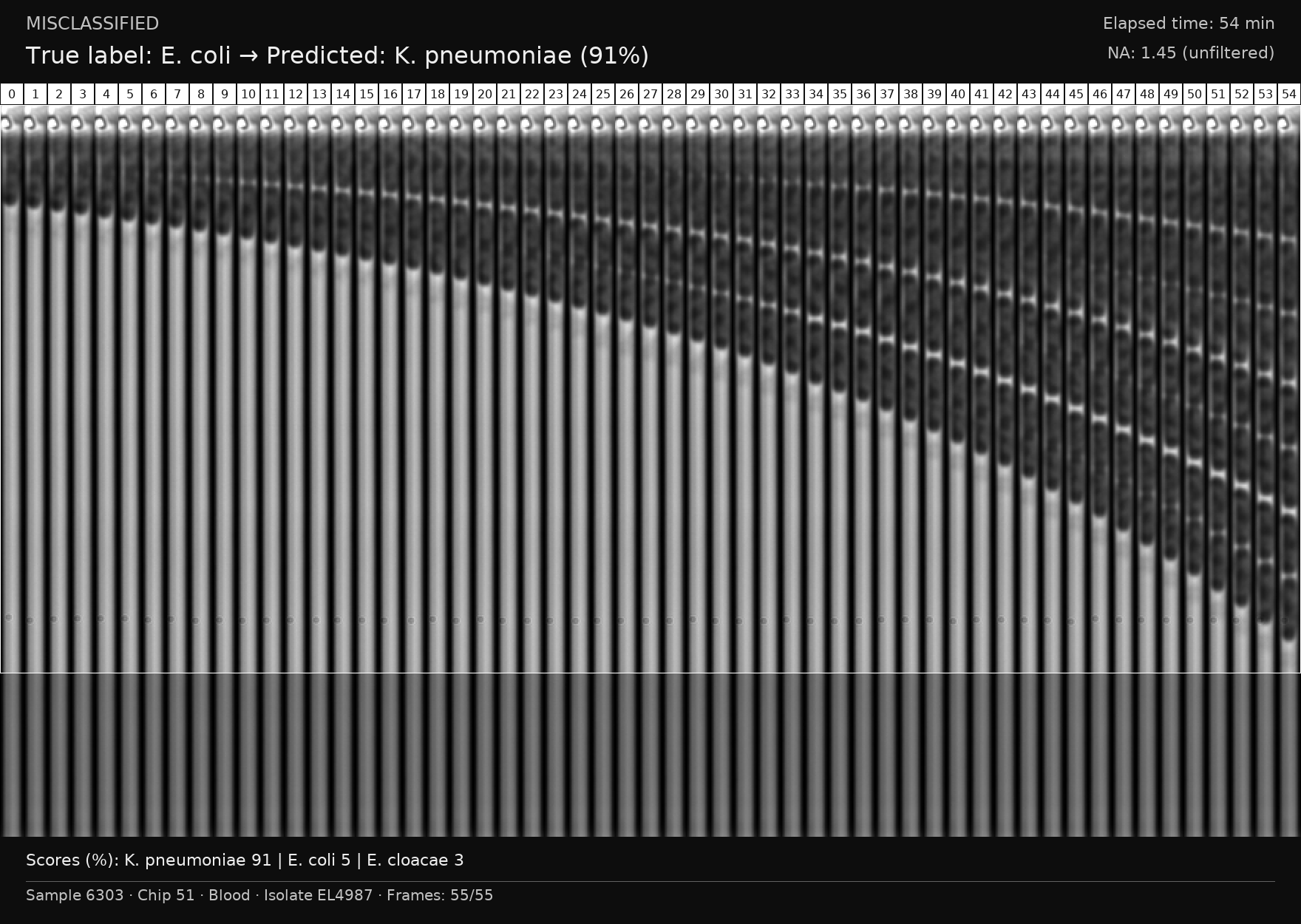

### S22 Fig

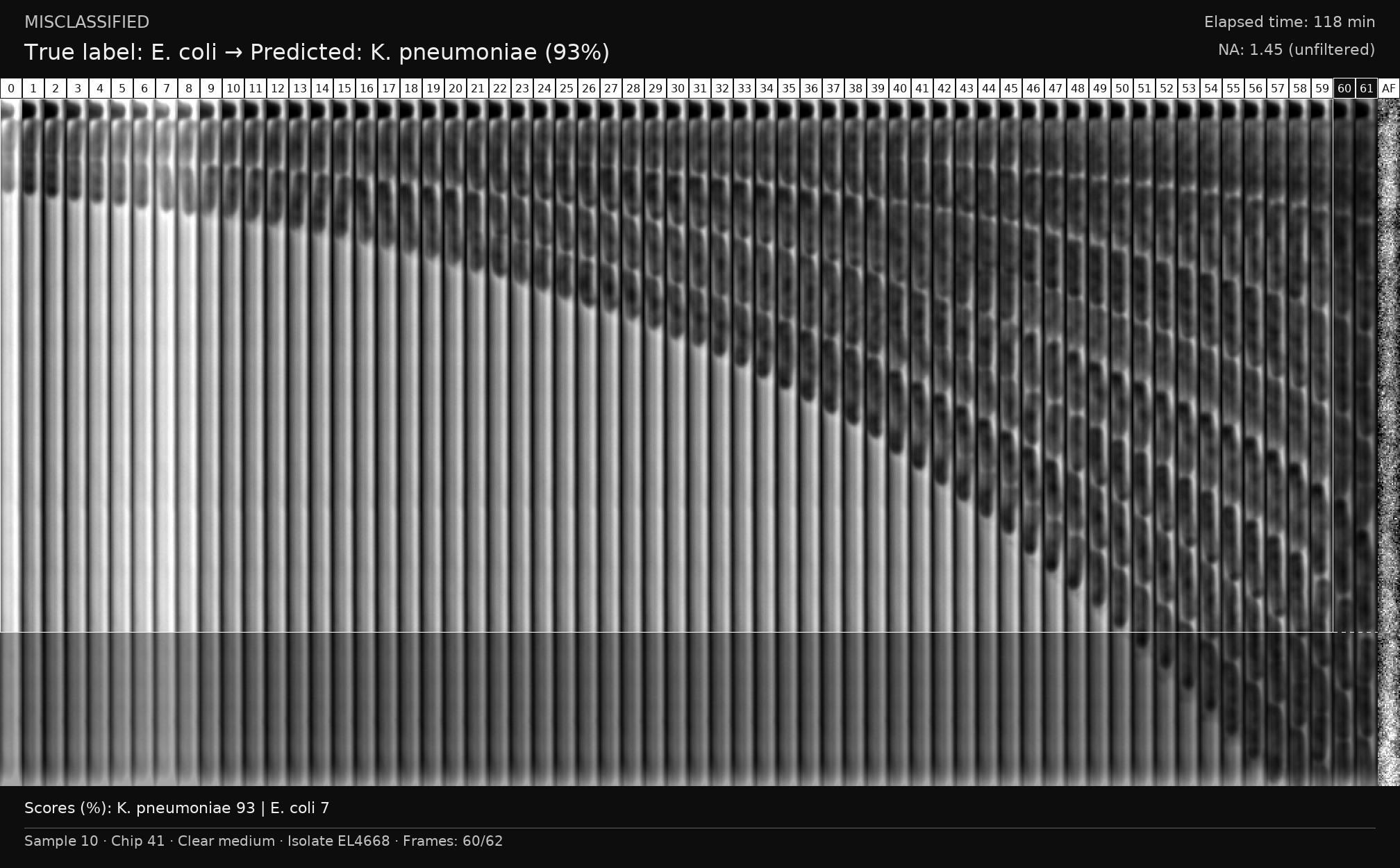

### S23 Fig

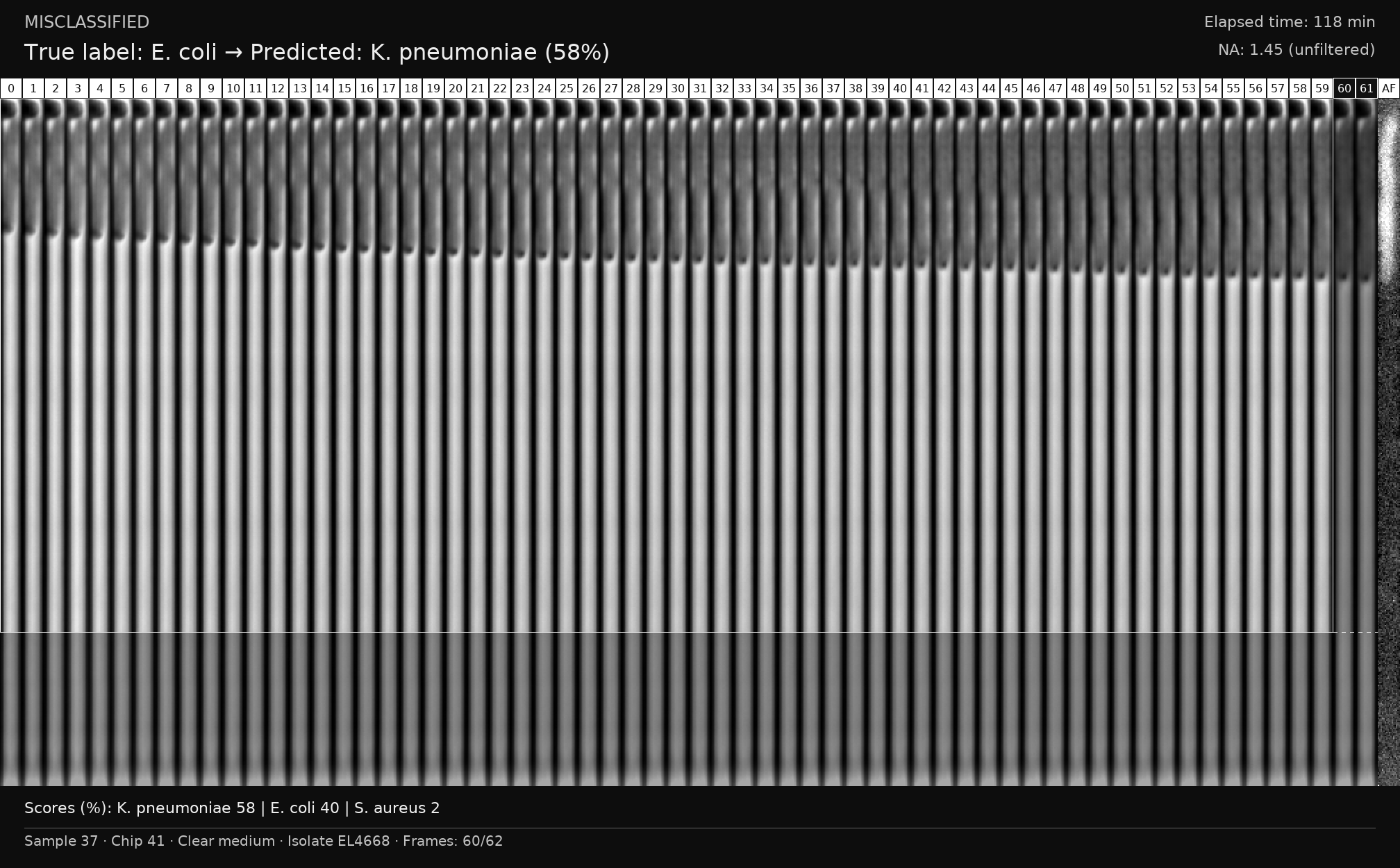

### S24 Fig

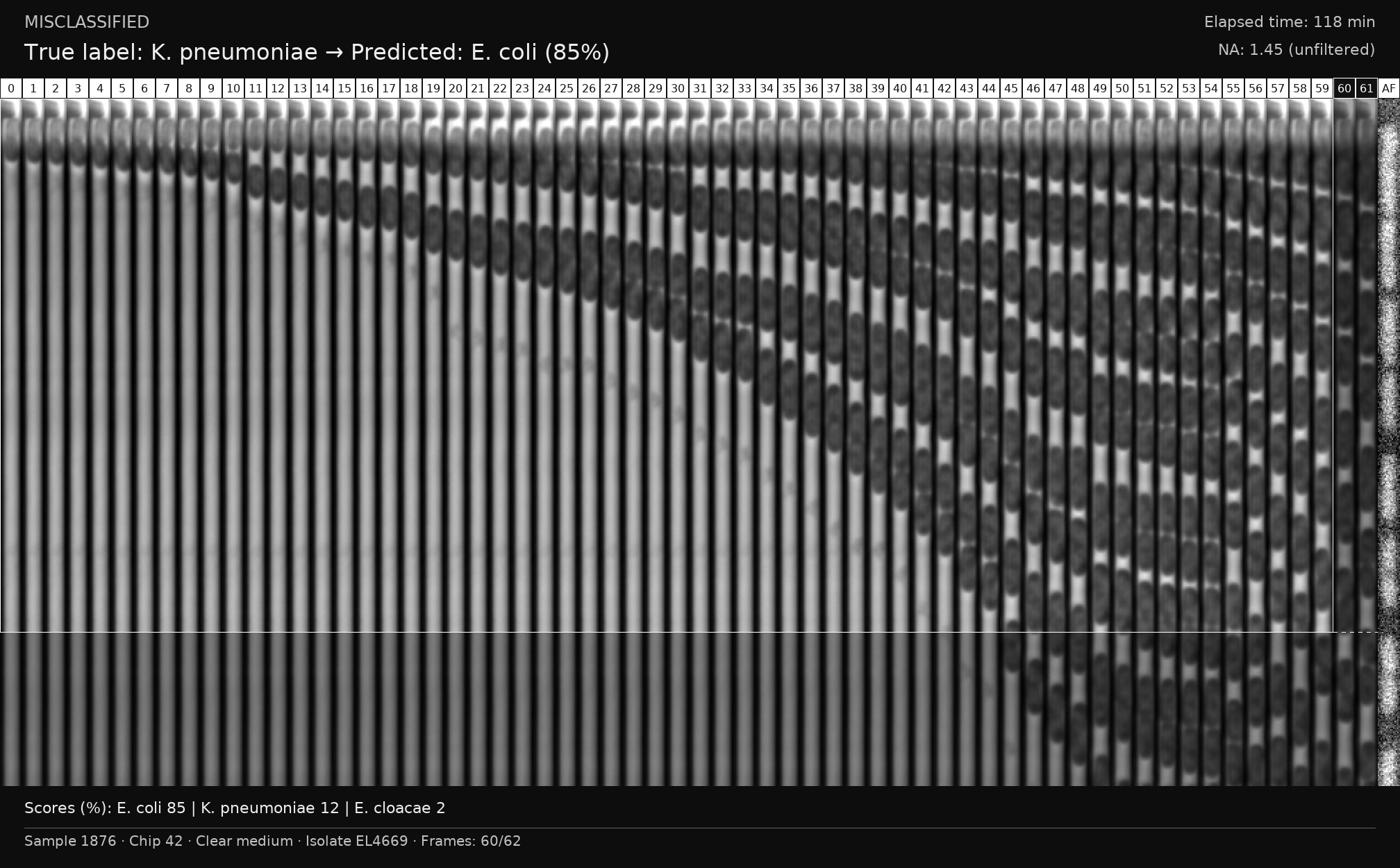

### S25 Fig

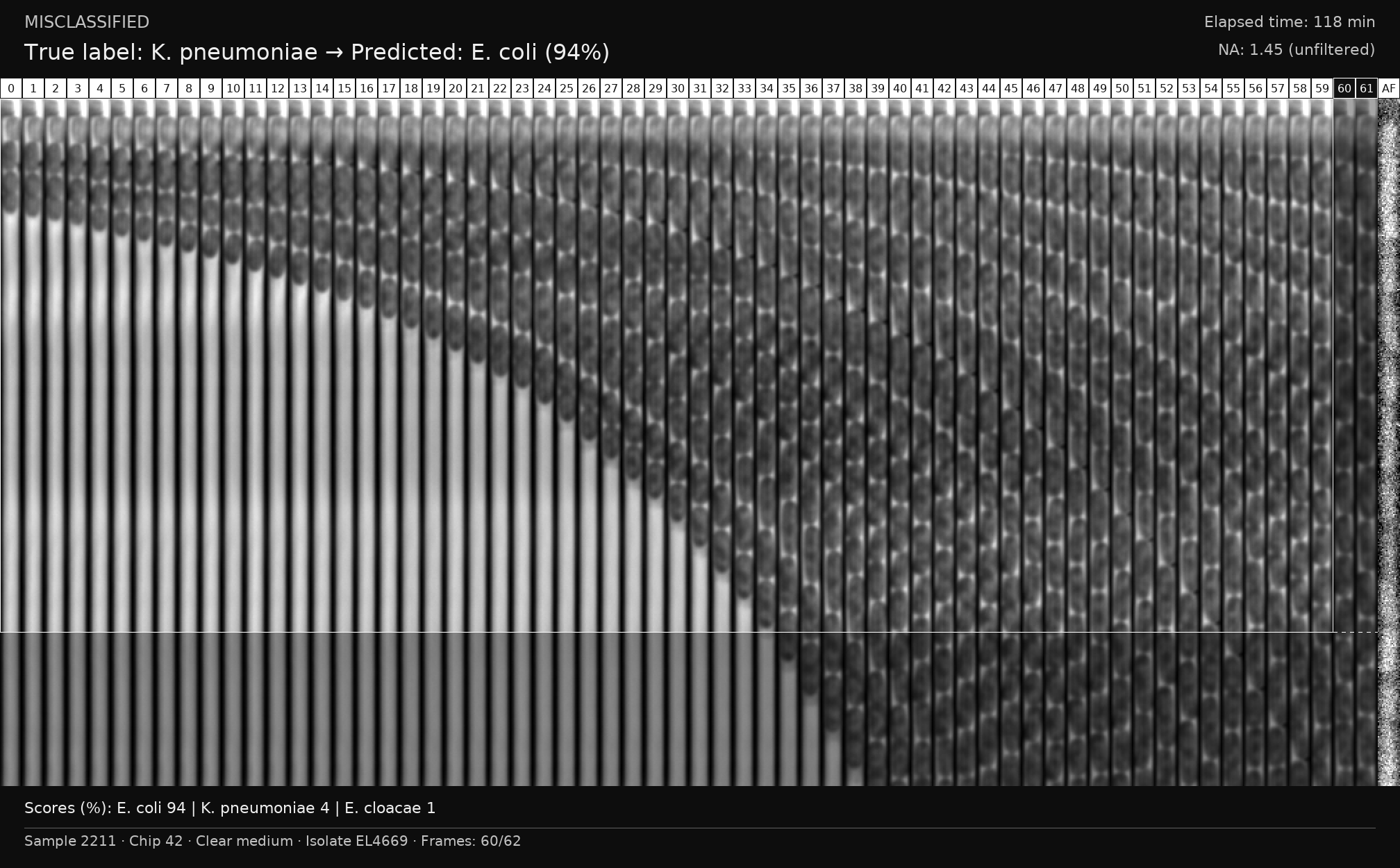

### S26 Fig

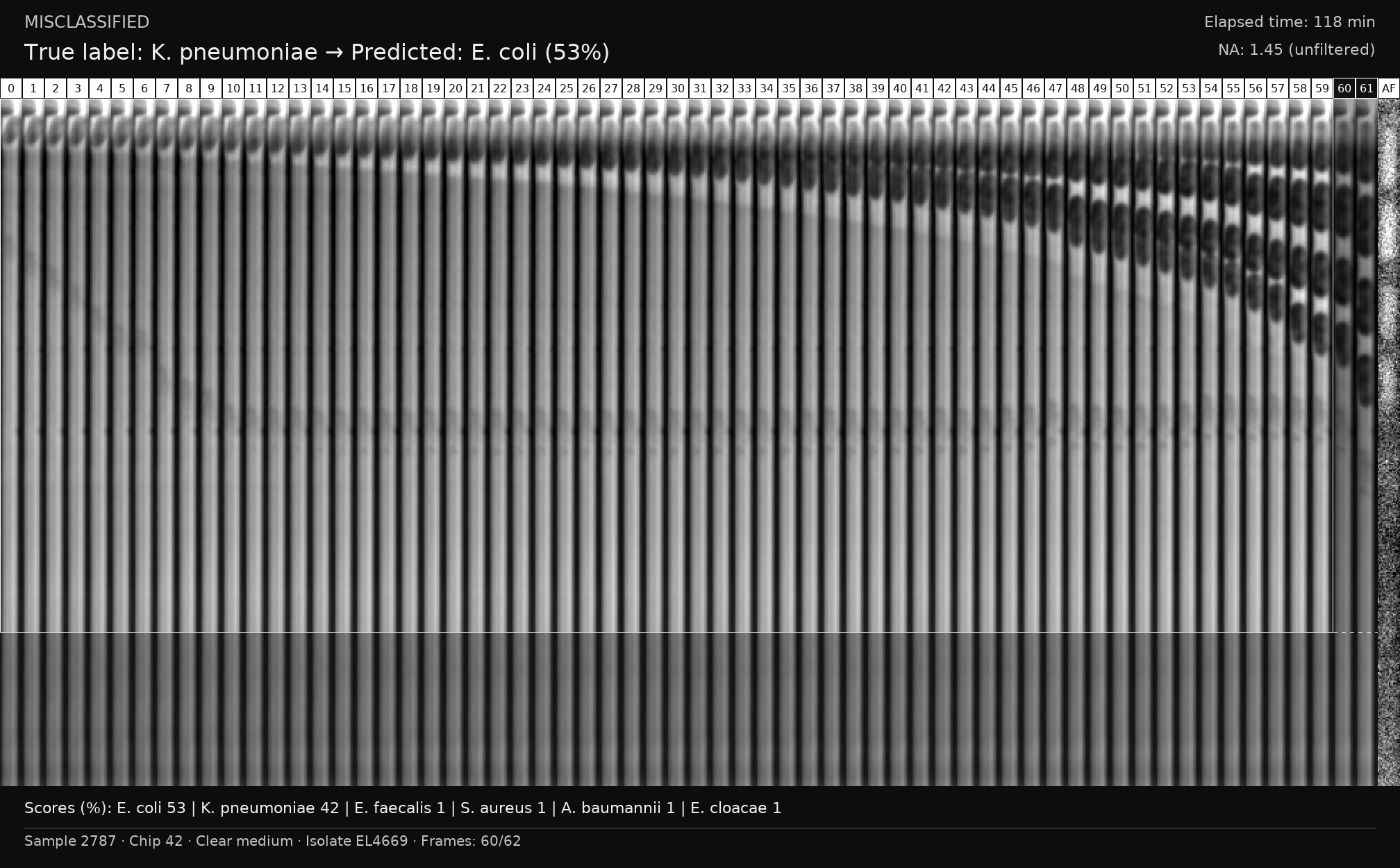

### S27 Fig

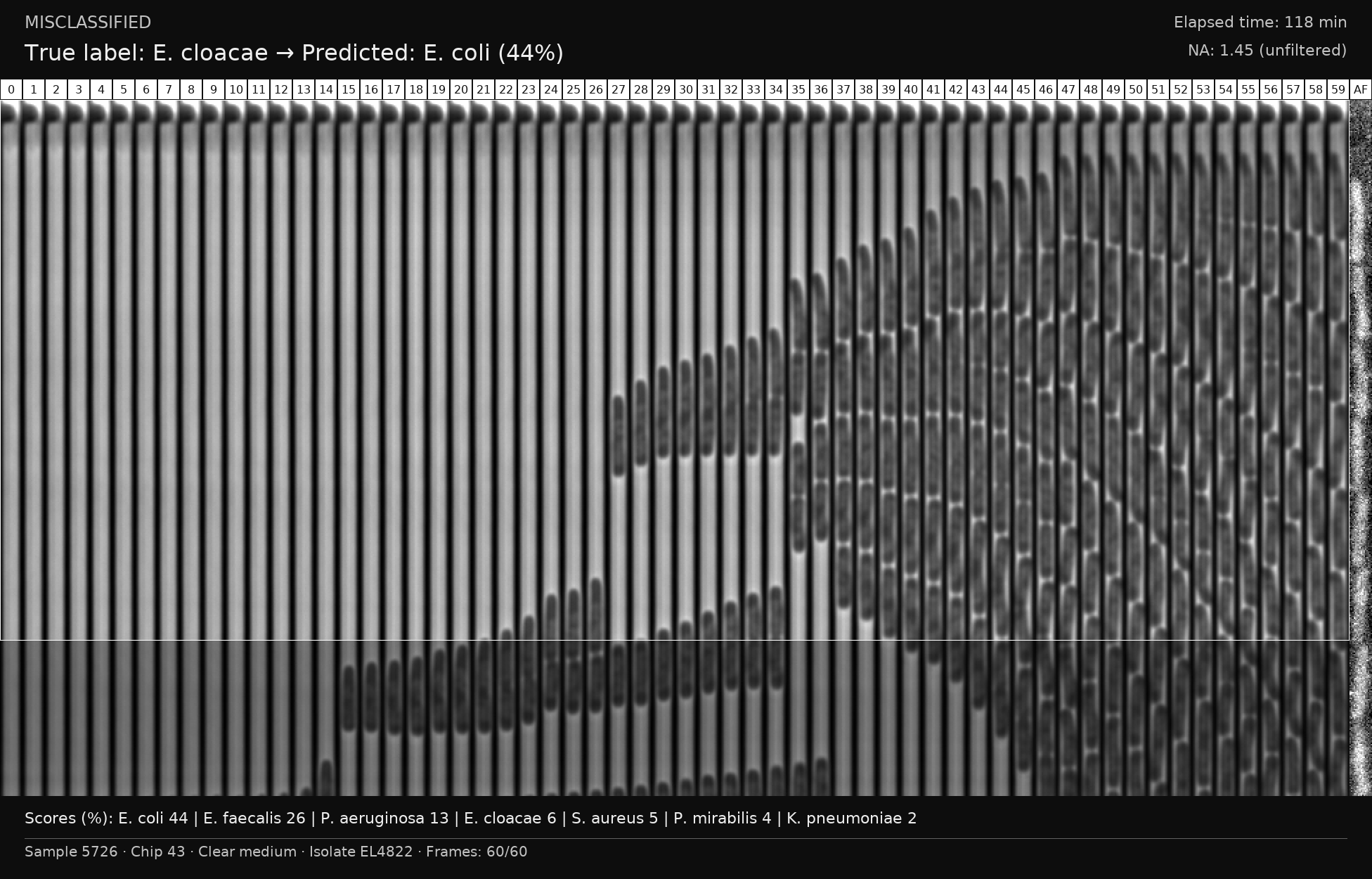

### S28 Fig

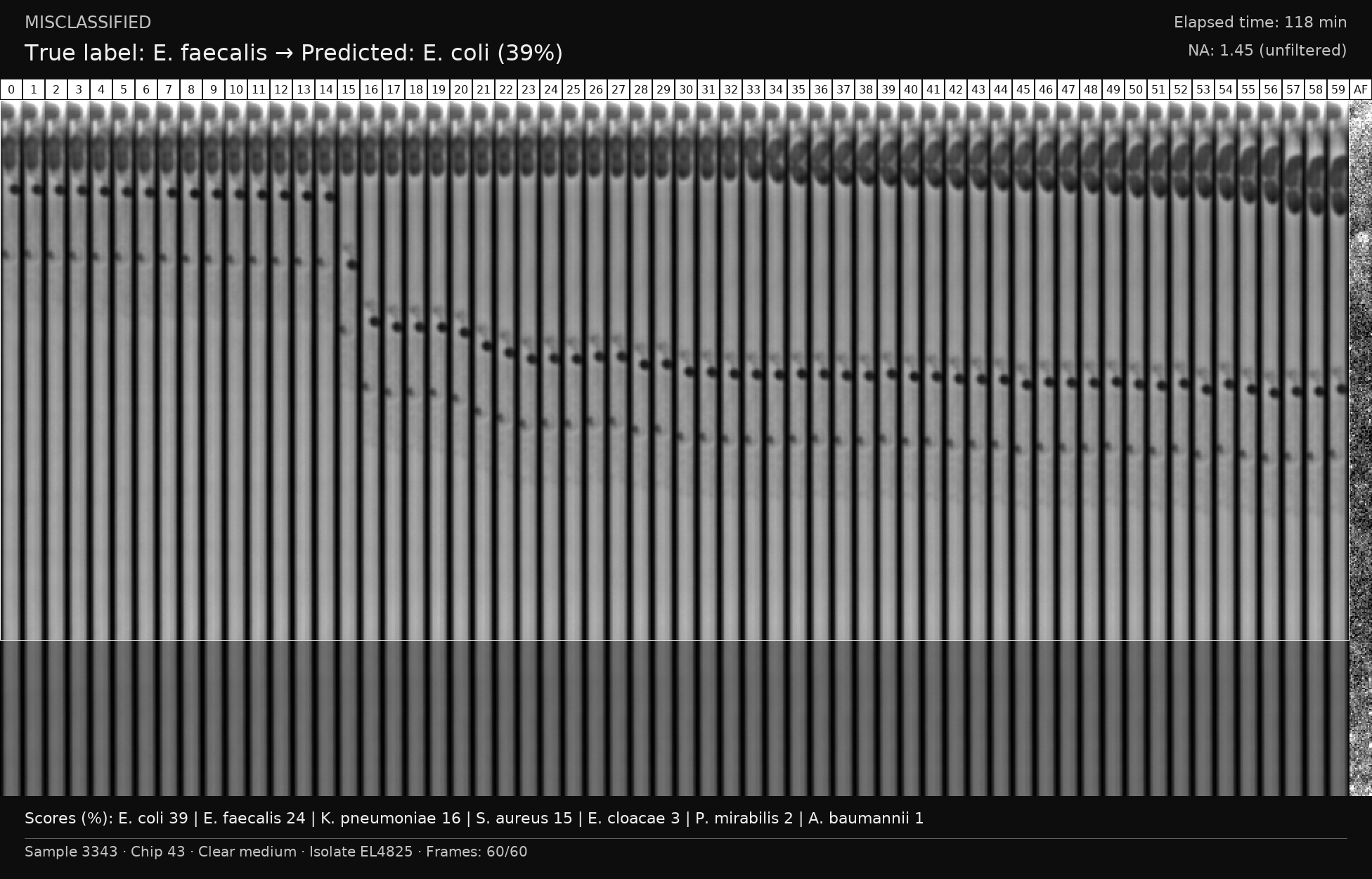

### S29 Fig

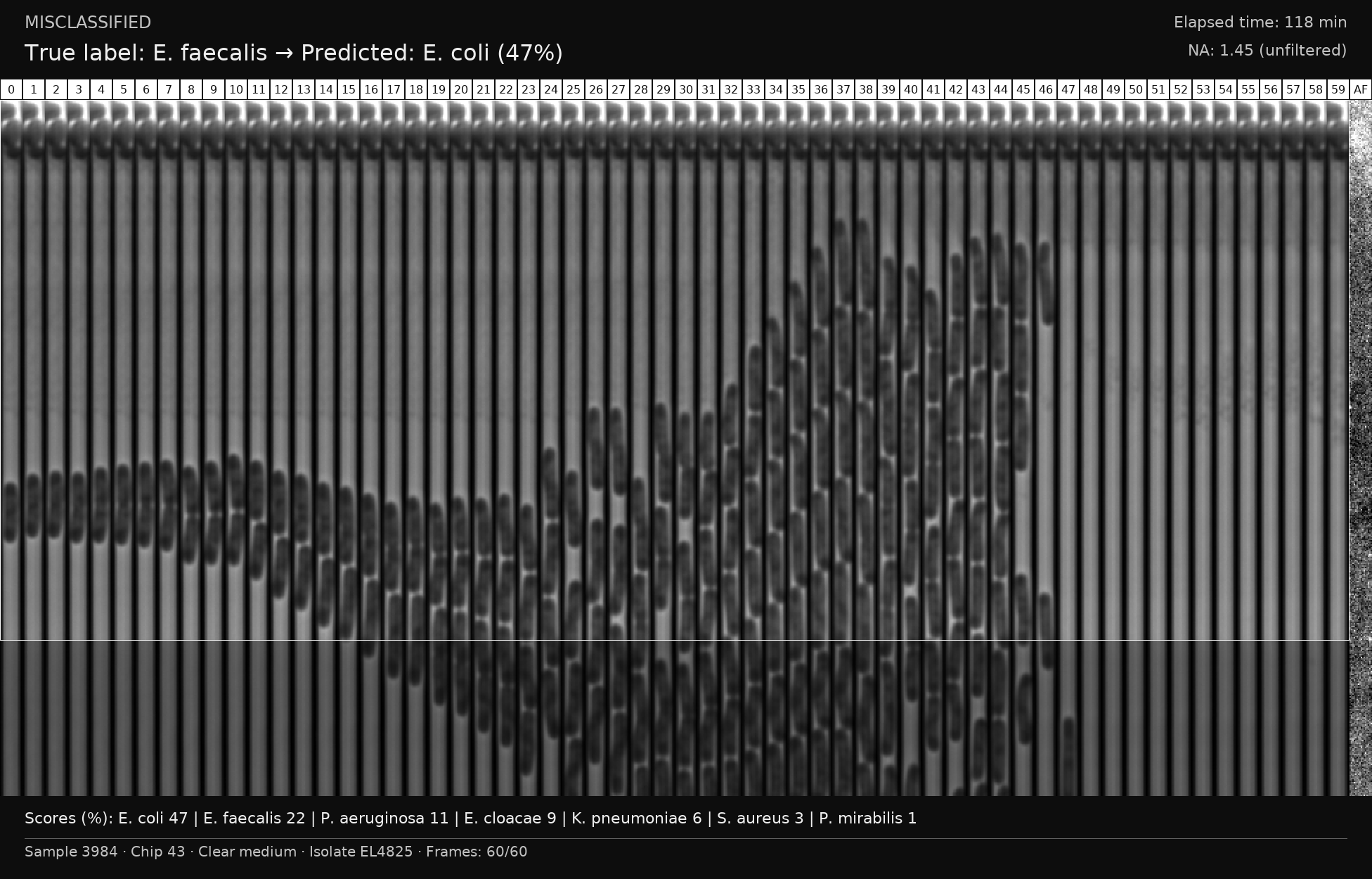

### S30 Fig

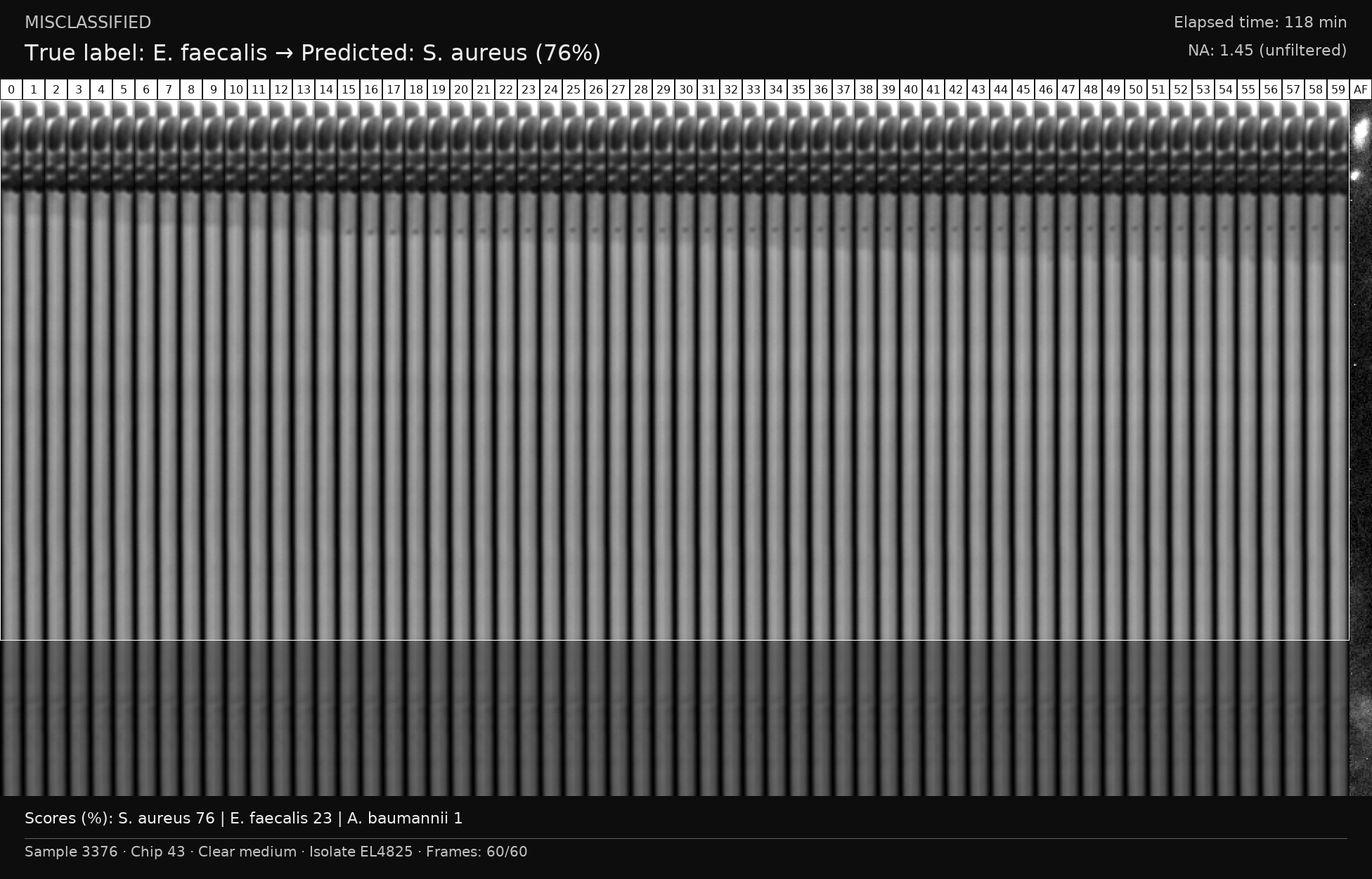

### S31 Fig

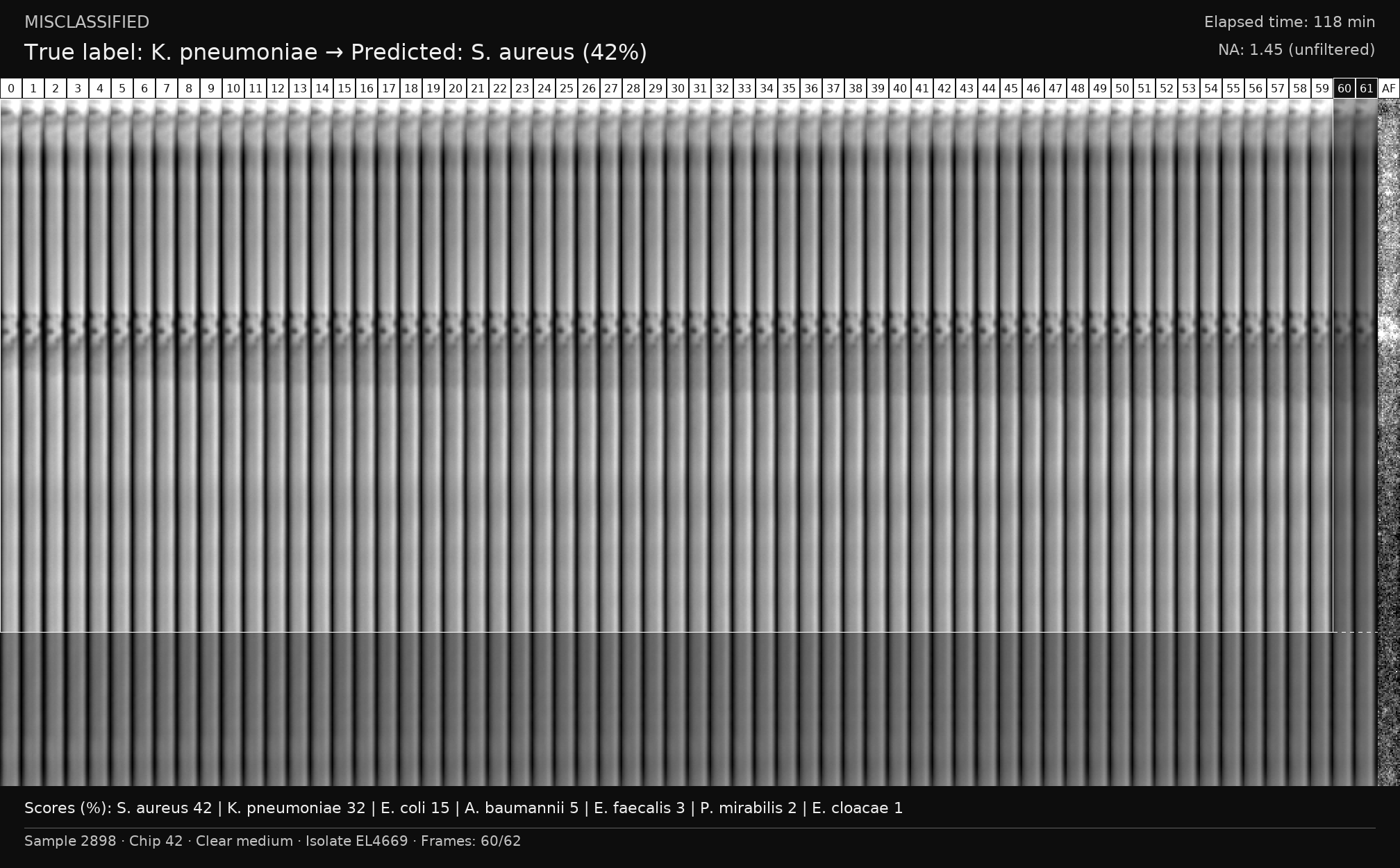

### S32 Fig

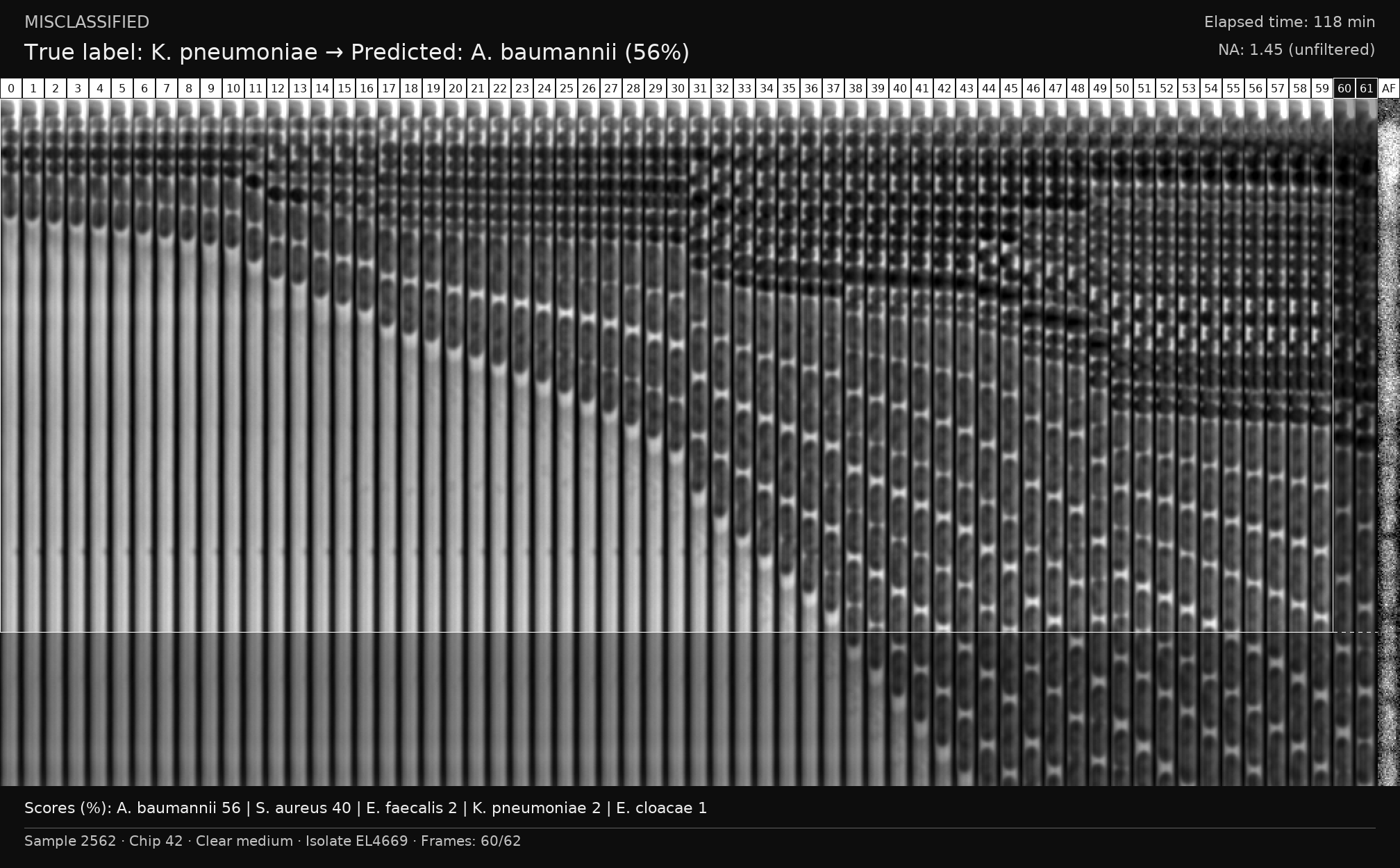

### S33 Fig

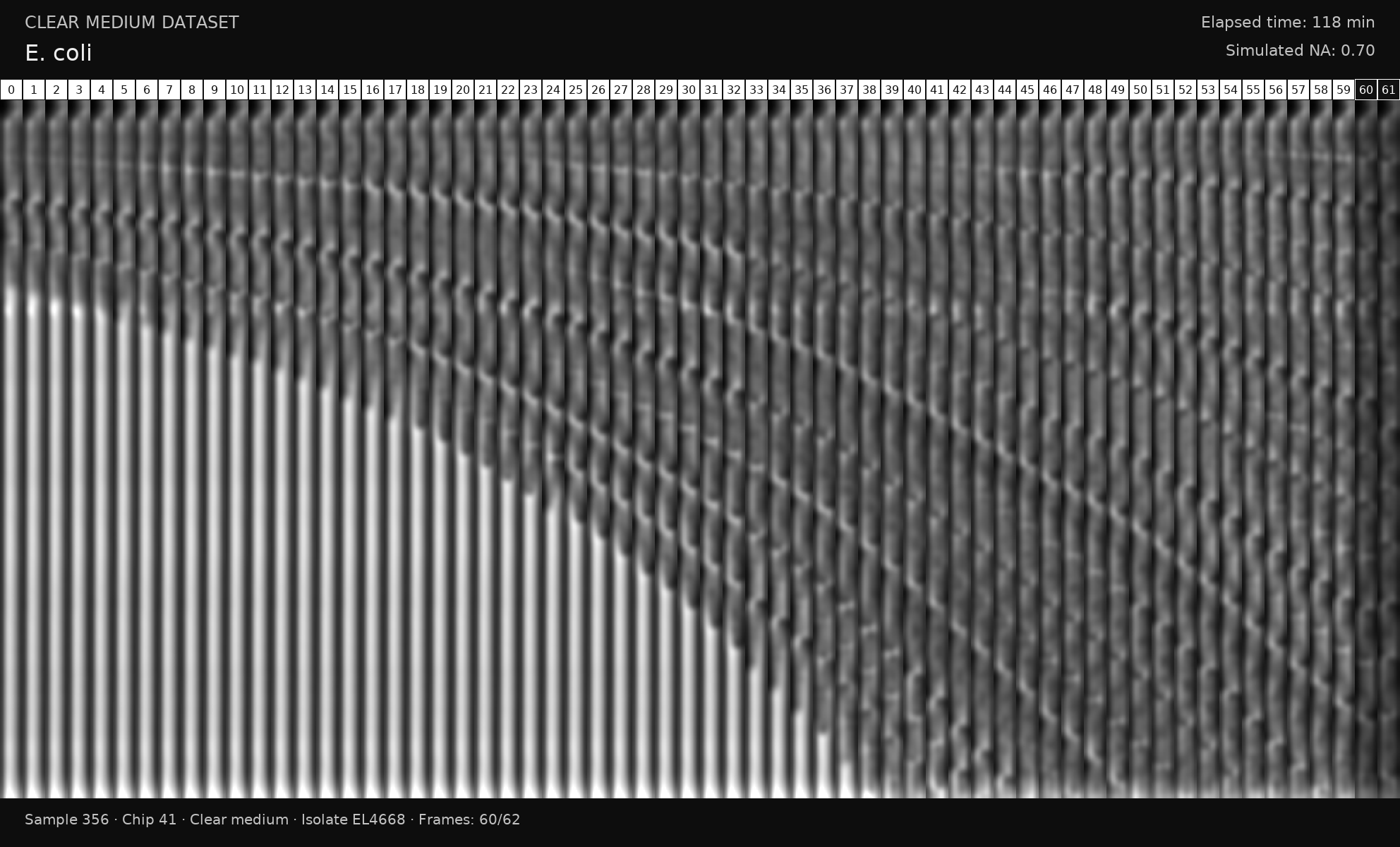

### S34 Fig

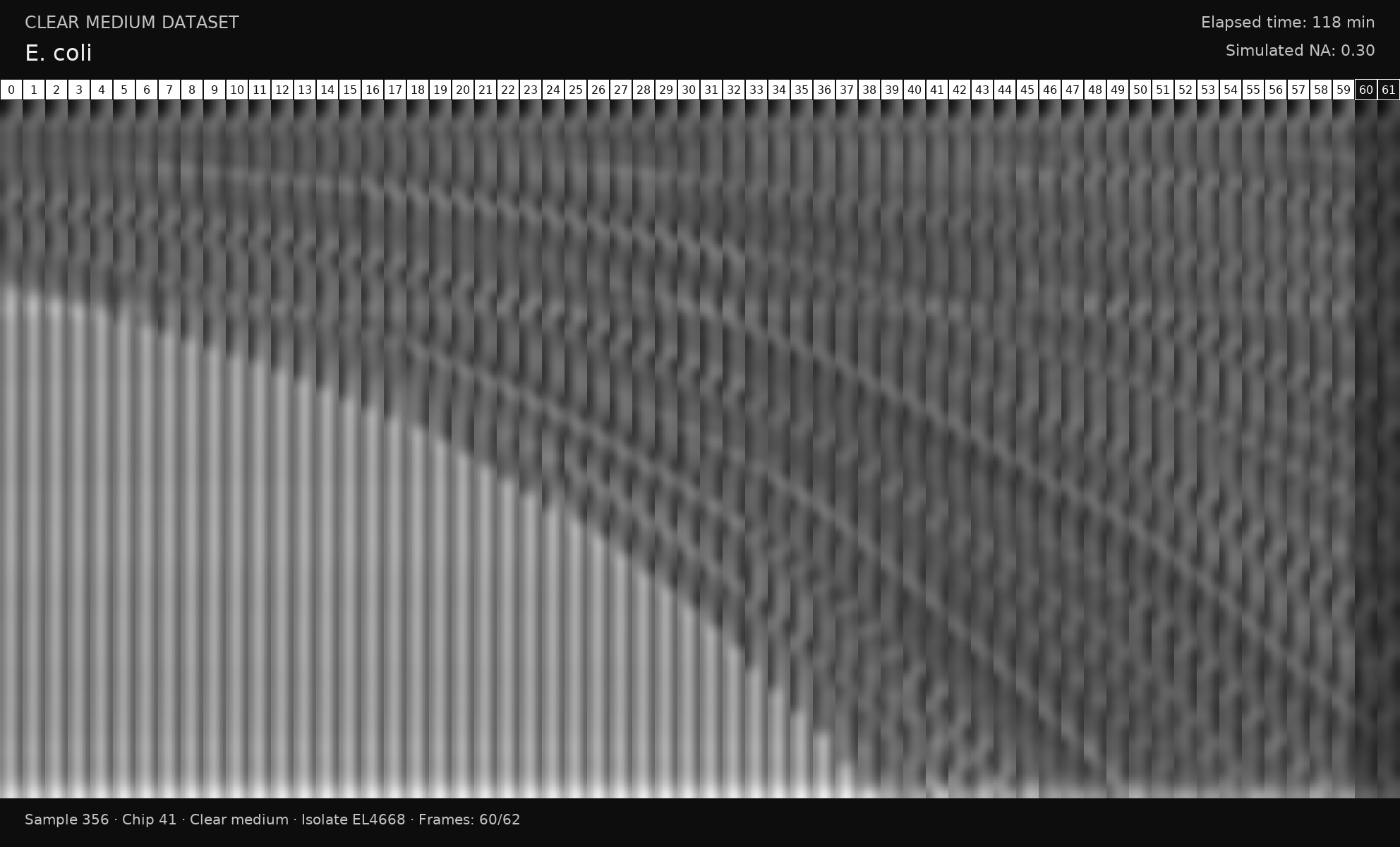

### S35 Fig

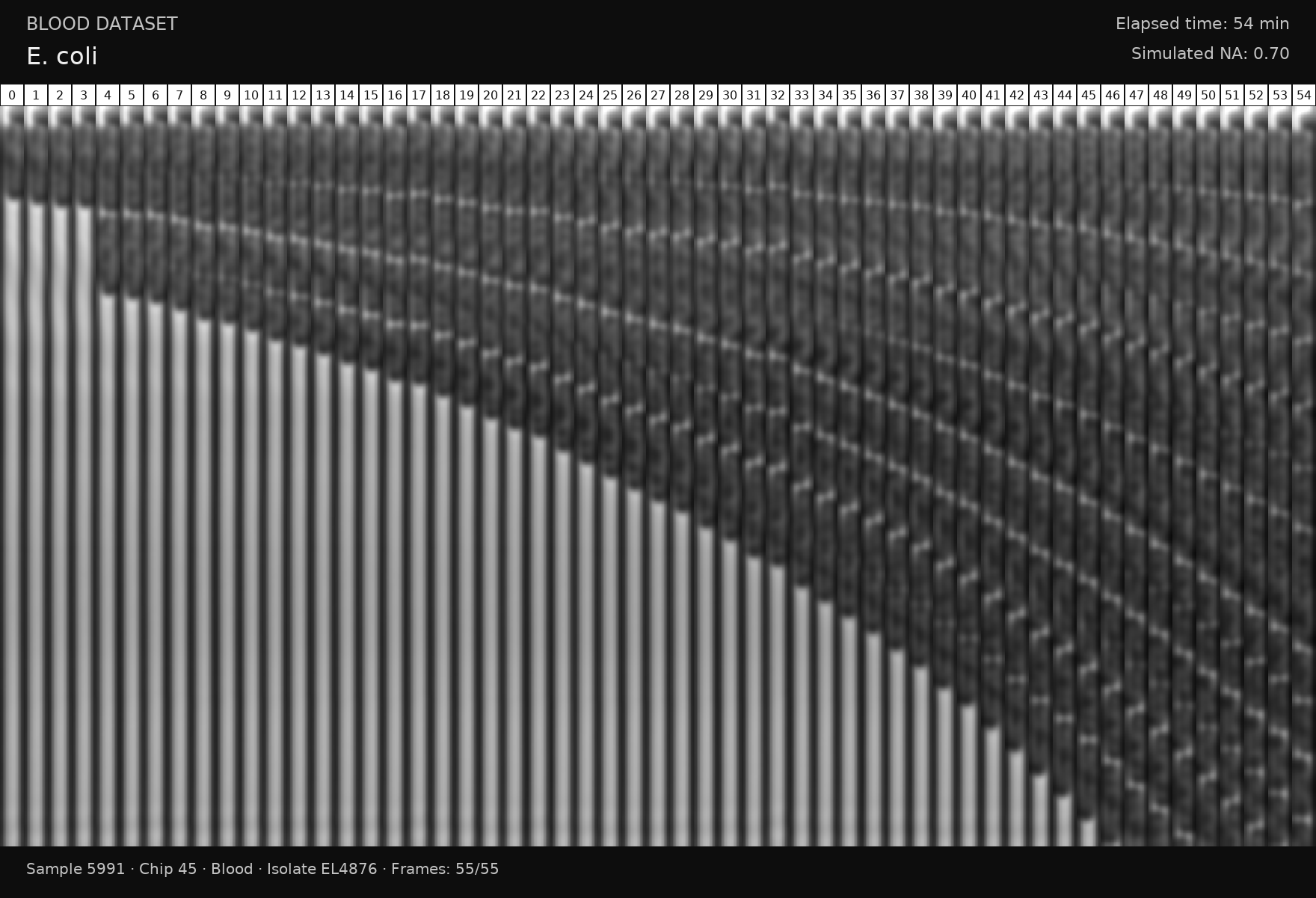

### S36 Fig

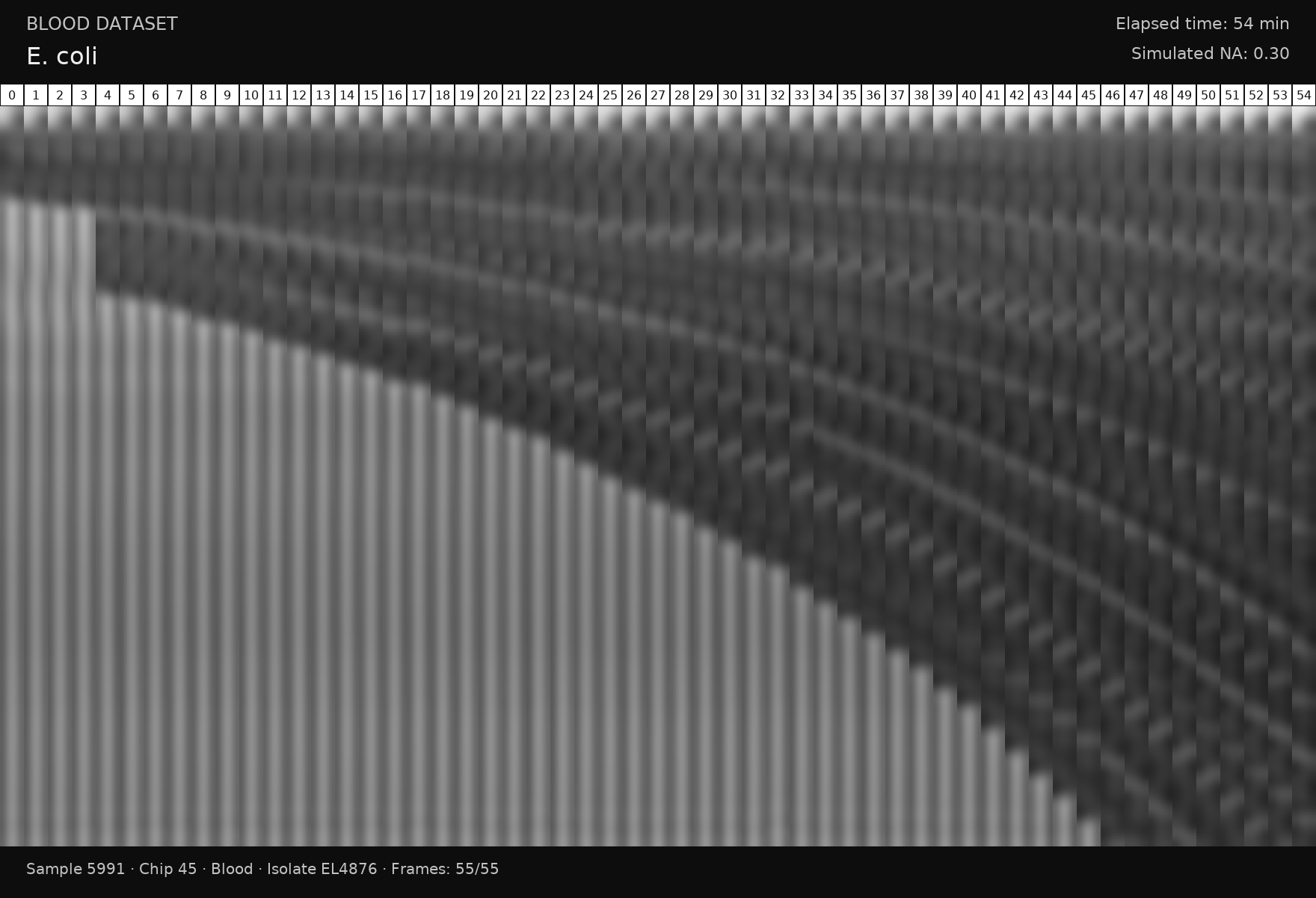

### S37 Fig

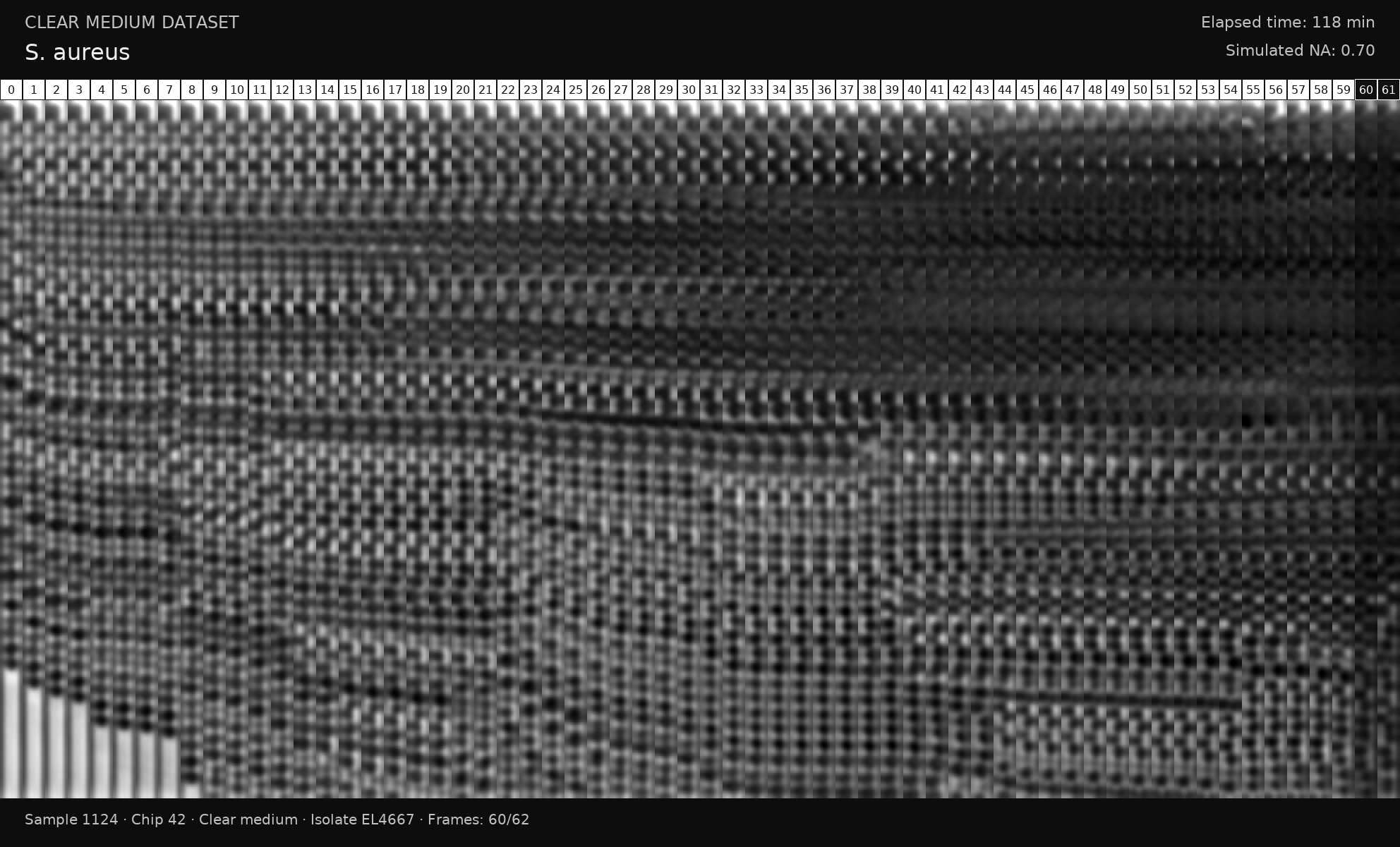

### S38 Fig

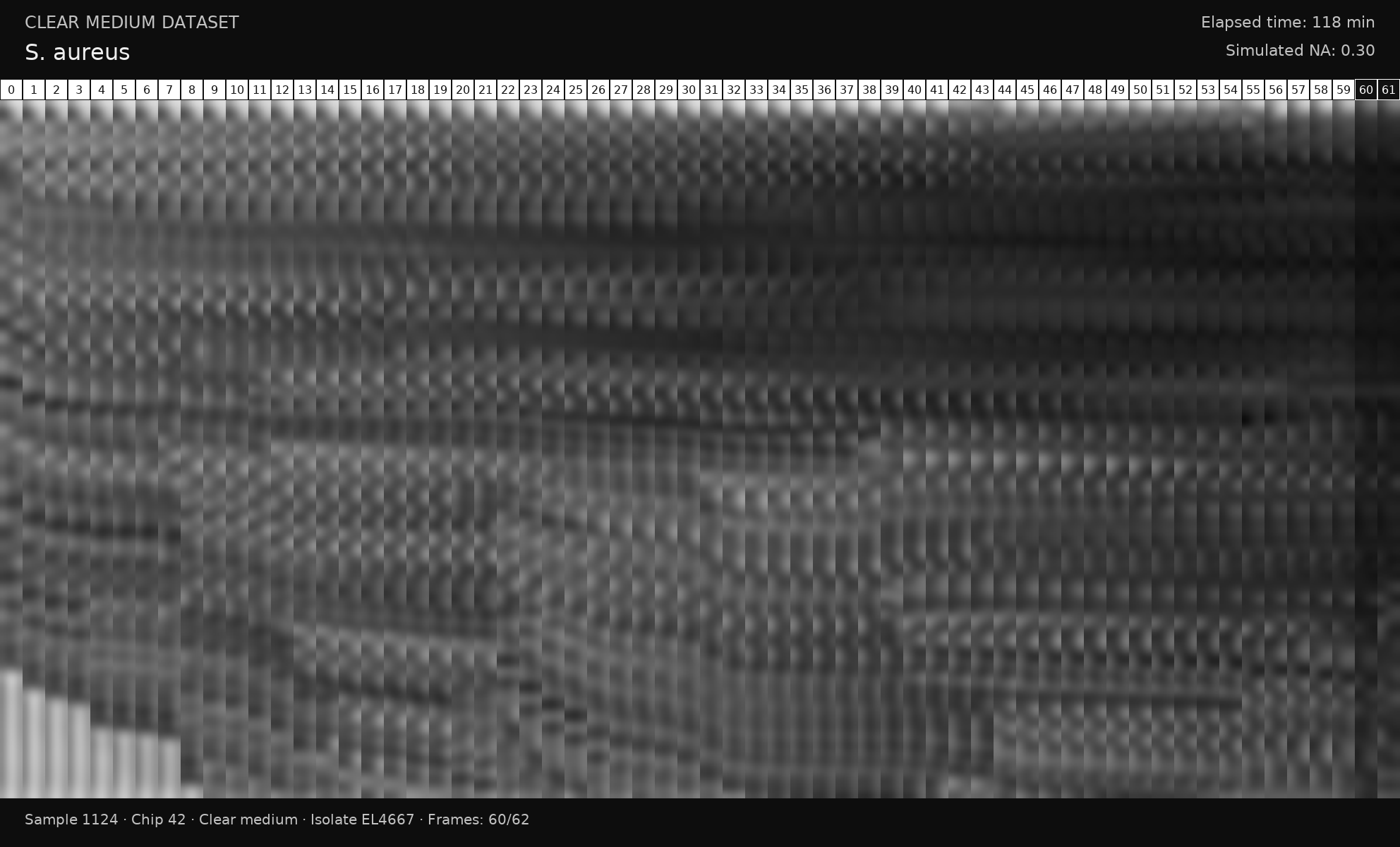

### S39 Fig

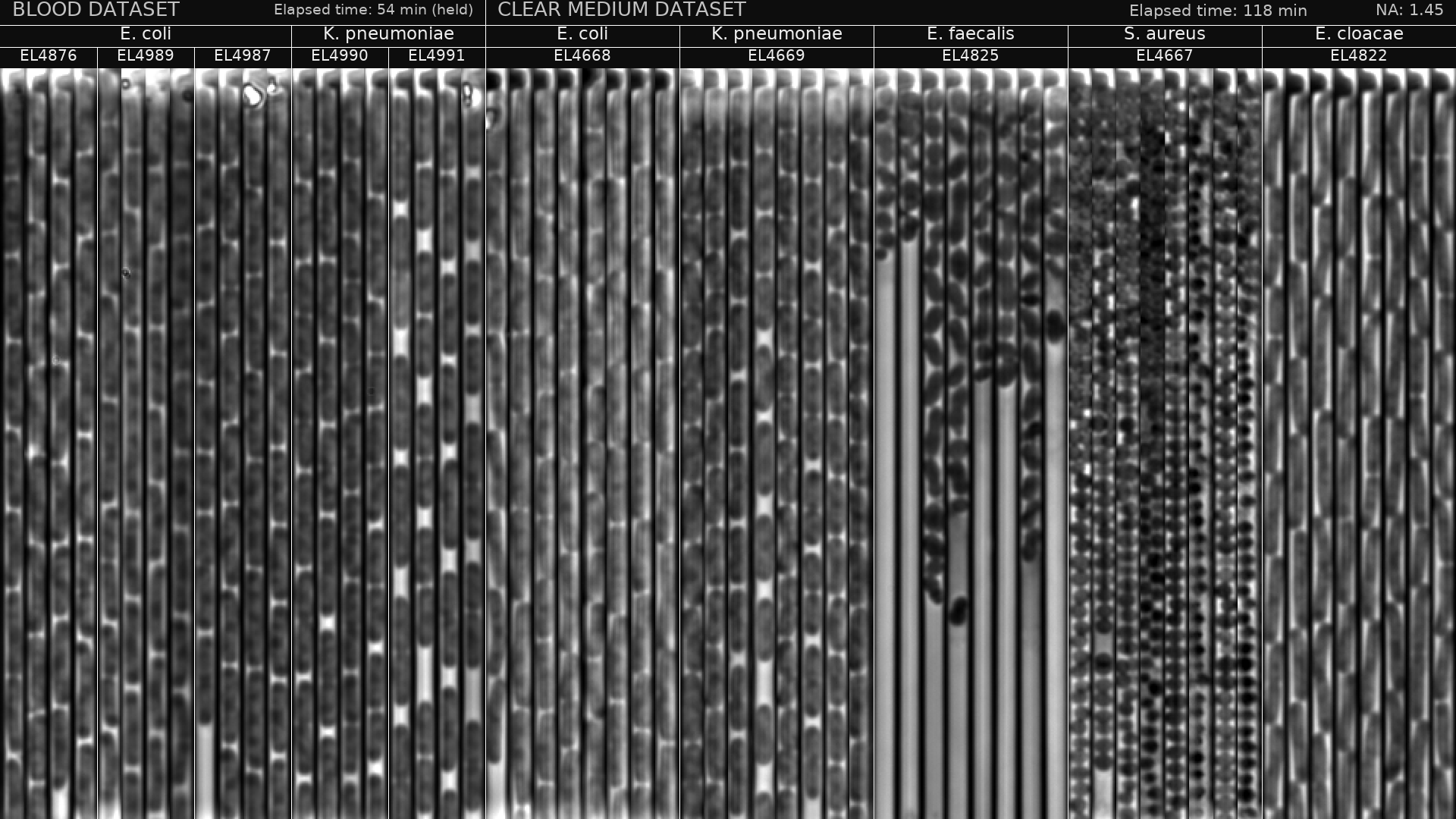

### S40 Fig

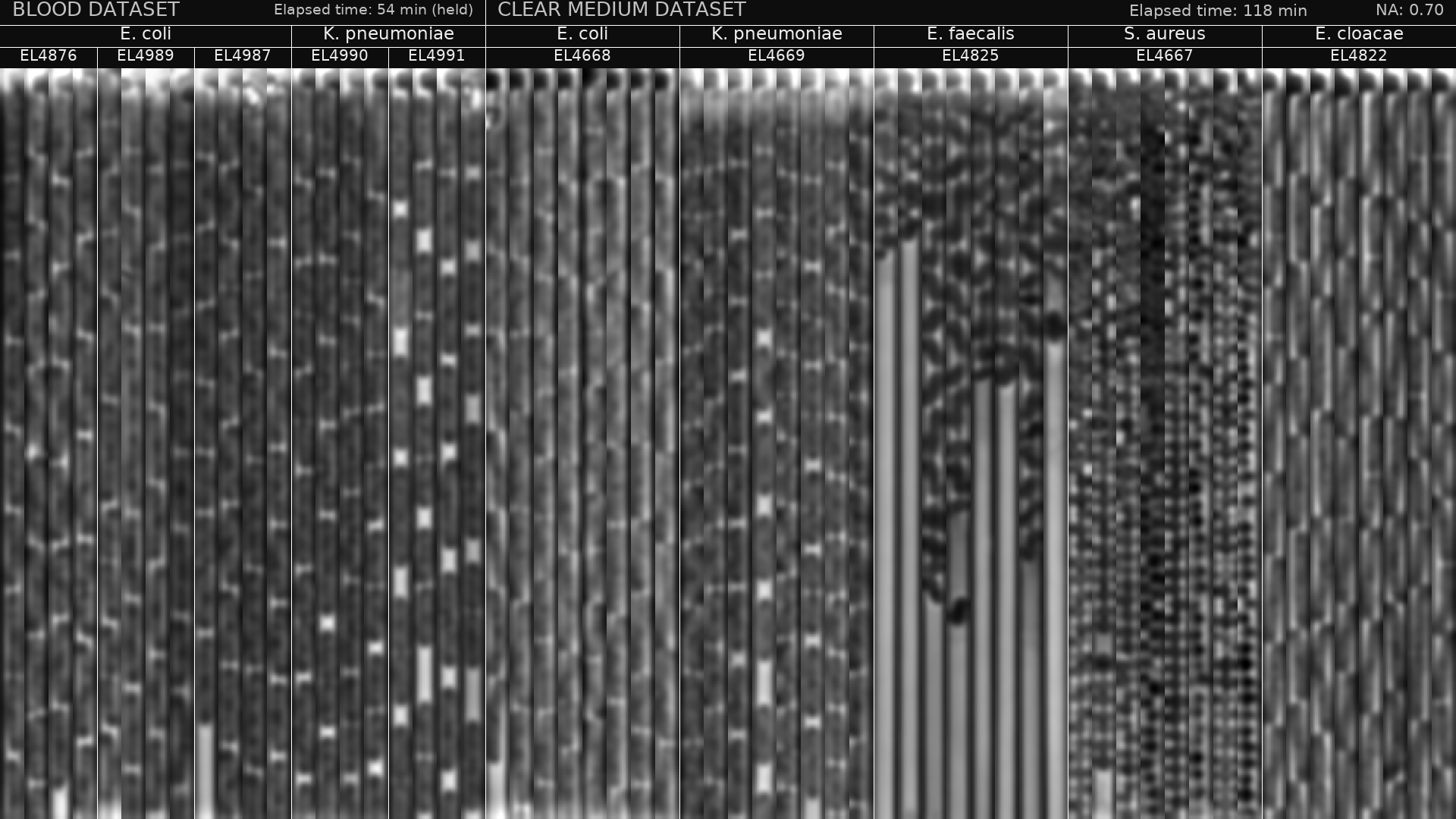

### S41 Fig

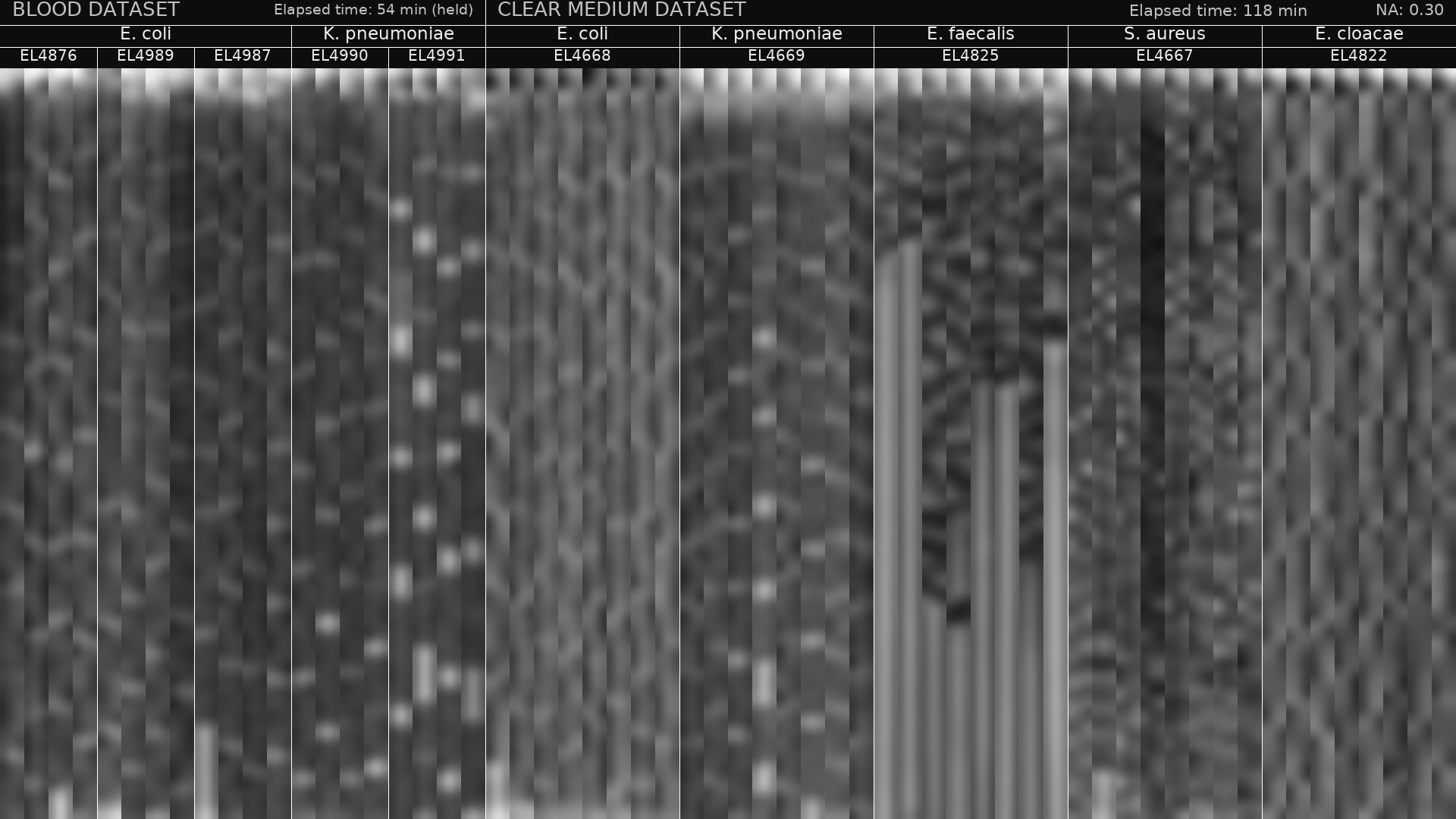
